# Substitution and Concomitant Distractor Intrusions Reveal Different Neural Dynamics of Temporal Selection

**DOI:** 10.64898/2026.11.22.768379

**Authors:** Zong Meng, Qi Chen, Liqin Zhou, Ke Zhou, Antao Chen

## Abstract

Selective attention requires target identification and exclusion of irrelevant information. We distinguished substitution and concomitant distractor intrusions in 29 adults performing rapid serial visual presentation with uncertain target number. Substitution reports, in which only the distractor was reported, showed an attenuated and delayed N2 posterior contralateral (N2pc) component and a smaller P3b; intrusion probability decreased with target-distractor numerical distance. Concomitant reports included both identities without reliable differences in measured averaged event-related potentials from correct reports of the same sequence. However, target-first concomitant reports were distinguishable through distributed activity and showed reduced left-frontal power, predominantly in theta. Comparisons with correct single-and dual-target reports indicated that reporting two identities did not simply reproduce genuine two-target processing. These findings distinguish processing profiles of substitution and concomitant intrusions and extend temporal-selection accounts from attentional engagement to subsequent regulation of which processed identities enter report.

**Impact statement:** Preserving a target and excluding a nearby distractor place different demands on temporal selection, revealing how irrelevant information can enter report even when target identification succeeds.

## Introduction

Selective attention enables observers to extract goal-relevant information from a changing environment. This function is particularly important when visual events occur in rapid succession and temporally adjacent inputs compete for processing (Denison, 2024). Yet target-evoked attentional enhancement is not confined to the target itself. According to attentional episode theory, detecting a target initiates a brief period of enhancement that can extend to neighboring items within the same attentional episode (e.g., Bowman & Wyble, 2007; Callahan-Flintoft et al., 2018; Olivers & Meeter, 2008; Wyble et al., 2009, 2011; Zivony & Eimer, 2022b). This continuity can benefit successive targets. In rapid serial visual presentation (RSVP), identification of a second target is often relatively preserved when it immediately follows the first, compared with the impairment typically observed at slightly longer intervals during the attentional blink (Raymond et al., 1992). This lag-1 sparing (e.g., Chun & Potter, 1995; Visser et al., 1999) can also extend across longer sequences of consecutive targets (e.g., Di Lollo et al., 2005; Olivers et al., 2007). However, neighboring irrelevant items may also benefit from enhancement. An item that lacks the target-defining feature but belongs to the reportable category can consequently be mistaken for the target. For example, when reporting colored digits, observers sometimes report the gray digit immediately following the target—the posttarget distractor (PTD). Such distractor intrusions demonstrate that enhancement supporting consecutive targets can also allow irrelevant identities to enter memory and report (e.g., Zivony & Eimer, 2020, 2021a, 2021b, 2022a).

An important explanation of distractor intrusion is the temporal variability account of attentional engagement (Zivony & Eimer, 2021a, 2022b). This account proposes that target-defining features trigger transient attentional enhancement, but the onset of enhancement varies across trials. When engagement is timely, the target representation receives sufficient enhancement to gain an advantage in subsequent working-memory encoding. When engagement is delayed, backward masking may already have weakened the target representation, whereas the PTD is available to benefit from enhancement and may consequently replace the target in report. Event-related potential (ERP) findings support this account: Relative to correct reports, distractor intrusions are associated with later onset of the N2 posterior contralateral (N2pc) component (Zivony & Eimer, 2020, 2021a, 2021b) and, in some experiments, smaller N2pc amplitudes (Zivony & Eimer, 2021a). Extending this evidence, Hosseini et al. (2024) used dynamic time warping (DTW) to show that intrusion trials exhibited delays in both the N2pc and the subsequent parietal P3b. The latencies of these components were coupled within both correct-report and intrusion conditions, although coupling was weaker during intrusion. These findings link the timing of initial attentional engagement to subsequent encoding, suggesting that variability in engagement timing may influence which information gains access to working memory.

In continuous visual behavior, however, observers often cannot anticipate how many relevant events will occur and must use incoming information to determine whether further targets are present. When target number is uncertain across trials, distractor intrusion may therefore take forms beyond target substitution. Zivony and Eimer (2021b) intermixed single- and dual-target trials, with the two targets appearing consecutively at the same location, and manipulated target-number expectations by varying the relative frequency of these trials. Even on trials containing only one target, observers sometimes reported both the target and the PTD, either in their presentation order or in the reverse order. Such joint reports were more frequent when two targets were expected. Related findings emerged when observers knew that each trial contained only one target: When required to provide two distinct guesses about its identity, they sometimes reported both the target and the PTD, with second reports of the other item exceeding chance (Zivony & Eimer, 2020). Thus, reporting the PTD does not necessarily entail failing to report the target. We accordingly distinguish *substitution intrusion* (DI_D), in which the PTD is reported without the target, from *concomitant intrusion*, in which the PTD is reported in addition to the target. Concomitant intrusion includes target-then-PTD reports in presentation order (DI_TD) and PTD-then-target reports in reverse order (DI_DT). Although previous work has examined joint reports and their relationship to working-memory access, the similarities and differences between these intrusion forms in attentional engagement, subsequent encoding, and maintenance require further clarification. In particular, does concomitant intrusion also exhibit delayed engagement, and is reporting one versus two identities during intrusion associated with neural activity resembling correct processing of one versus two actual targets?

Identifying which aspects of an attentional episode differ between intrusion forms requires combining event-related potentials (ERPs) associated with attentional engagement, encoding, and maintenance. The N2pc can be used to track initial target-evoked attentional engagement (e.g., Callahan-Flintoft et al., 2018; Eimer, 1996; Zivony et al., 2018), whereas the subsequent distractor positivity (Pd) has been associated with attention termination or distractor suppression (e.g., Gaspelin et al., 2023; Hickey et al., 2009; Sawaki et al., 2012), providing information about how target-triggered enhancement is subsequently regulated. Attentional episode theory emphasizes that neighboring items can receive enhancement within the same episode, while forming reportable working-memory representations additionally requires binding identities to specific events. The P3b has been linked to these encoding and binding operations, as well as working-memory updating, informing how enhanced information enters subsequent memory processing (e.g., Craston et al., 2009; Polich, 2007; Verleger, 2020). The late positive complex (LPC), in turn, has been associated with processing devoted to working-memory maintenance (e.g., Gunseli, Meeter, et al., 2014; Ruchkin et al., 1990). Previous RSVP research found that consecutive targets elicited larger and more prolonged parietal positivity than a single target (Dell’Acqua et al., 2016). Examining earlier P3b and later LPC activity separately can therefore help determine whether the effects of a second target and distractor intrusion concern initial encoding, binding, and updating or subsequent maintenance-related processing. Contralateral delay activity (CDA) provides further information about working-memory load: Its amplitude typically increases with the number of maintained items before approaching an asymptote near an individual’s capacity limit (e.g., Luria et al., 2016; Vogel & Machizawa, 2004). Together, these components provide temporal reference points from attentional engagement to memory maintenance for comparing correct single- and dual-target processing with the two intrusion forms.

Beyond these component measures, multivariate pattern and time–frequency analyses can assess whether report outcomes differ in distributed activity and oscillatory dynamics. Time-resolved multivariate pattern analysis uses joint activity across electrodes to determine whether report outcomes exhibit distinguishable neural patterns and whether discriminative information generalizes across time, thereby characterizing the dynamics of processing patterns (King & Dehaene, 2014). Correct single- and dual-target conditions also provide reference patterns for examining features shared with intrusion outcomes, helping assess the relationship between the number of reported identities and overall processing patterns (Diedrichsen & Kriegeskorte, 2017). Time–frequency analysis characterizes changes in activity across time and frequency, including activity attenuated in ERP averages because its phase varies across trials (Makeig et al., 2004). These analyses complement component comparisons within selected electrodes and time windows, providing additional perspectives on the distributed activity and frequency-specific power associated with each intrusion form and its relationship to correct single- and dual-target processing.

The present study recorded electroencephalographic (EEG) activity during a lateralized RSVP task in which target number was uncertain across trials. A colored digit target was immediately followed at the same location by a gray letter, a gray digit PTD, or a second colored digit target, yielding single-target baseline (T), target–distractor (TD), and consecutive dual-target (TT) conditions, respectively. We addressed two related questions. First, how does a target-evoked attentional episode develop as a function of the relevance of subsequent input? Comparisons among the three correct-report conditions characterized changes from attentional engagement and subsequent regulation to encoding and maintenance. Second, which processing changes are associated with substitution and concomitant intrusion? Within identical TD stimulus configurations, we compared correct reports with both intrusion forms to assess the scope of the delayed-engagement account. Correct single- and dual-target processing additionally provided reference conditions for examining neural activity when one versus two identities were reported. By integrating ERP, multivariate pattern, and time–frequency analyses, this study brings target preservation and the exclusion of irrelevant identities into a common framework, helping explain how continuous attentional enhancement supports accurate selection and gives rise to different forms of selection error.

## Results

### Distinct Behavioral Profiles of Distractor Intrusions

#### Substitution and concomitant forms of distractor intrusion

In the TD condition, reports of the PTD produced three intrusion patterns: DI_TD, DI_DT, and DI_D (**Figure 1A**). Intrusion rates were calculated as percentages of all TD trials. A repeated-measures ANOVA revealed a significant effect of intrusion type (*F*(1.23, 34.41) = 30.36, *p*_GG_ < .001, ω^2^ = 0.40). DI_TD (*M* = 27.78, *SD* = 18.22) was more frequent than DI_D (*M* = 6.90, *SD* = 6.25; *t*(28) = 5.22, *p*_Holm_ < .001, 95% CI [10.69, 31.06], *d* = 1.74) and DI_DT (*M* = 6.59, *SD* = 7.88; *t*(28) = 6.56, *p*_Holm_ < .001, 95% CI [12.96, 29.40], *d* = 1.76). DI_DT and DI_D did not differ significantly (*t*(28) = −0.18, *p*_Holm_ = .856, 95% CI [−4.50, 3.90], *d* = −0.03; **Figure 1B**). Each intrusion type was also more frequent than other errors, defined as all TD responses that were neither correct nor one of the three intrusion types (*M* = 3.86, *SD* = 3.88): DI_TD, *t*(28) = 7.17, both *p* and *p*_Holm_ < .001; DI_D, *t*(28) = 2.39, *p* = .024, *p*_Holm_ = .048; and DI_DT, *t*(28) = 2.23, *p* = .034, *p*_Holm_ = .048 (**Table S1**).

**Figure 1.**
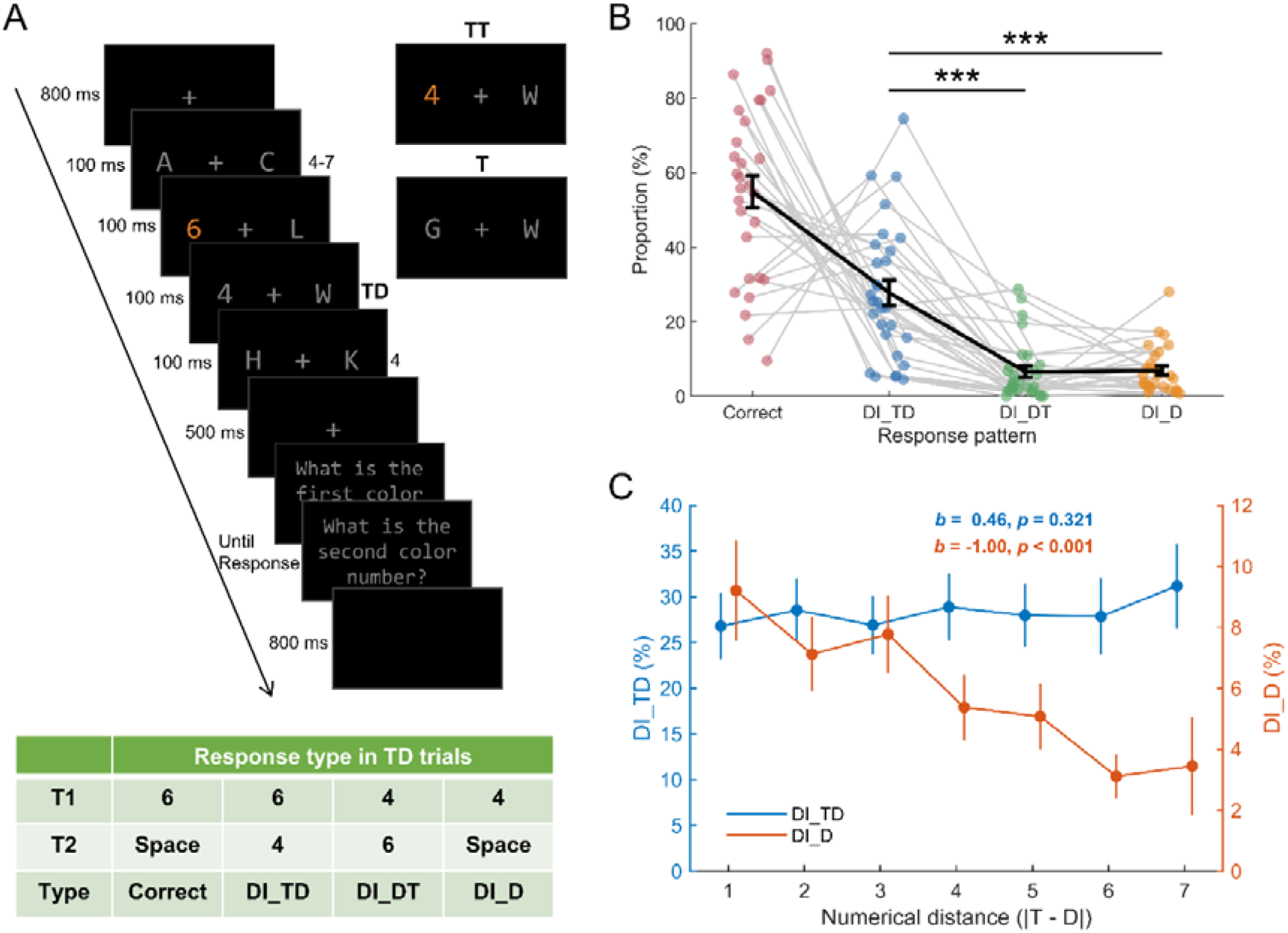
Task design and behavioral signatures of distractor intrusion. (**A**) Participants reported one or two colored digits in two lateral rapid serial visual presentation (RSVP) streams. The first target (T1) was immediately followed at the same location by a gray letter in single-target (T) trials, a gray digit posttarget distractor (PTD) in target–distractor (TD) trials, or a second colored target (T2) in two-target (TT) trials. Participants were informed that T, TD, and TT trials occurred in a 1:4:5 ratio. The table illustrates correct TD reports and three distractor intrusion (DI) outcomes: correct reports identified T1 and indicated that no second target was present; DI_TD reports identified T1 followed by the PTD; DI_DT reports identified the PTD followed by T1; DI_D reports identified only the PTD and indicated that no second target was present. **(B)** Correct-report and intrusion rates as percentages of all TD trials. Colored points and gray lines show paired participant values; black points and error bars show means (*M*) ± standard error of the mean (*SEM*). Brackets indicate Holm-corrected comparisons between intrusion types (*** *p*_Holm_ < .001). **(C)** DI_TD and DI_D rates by absolute T1–PTD numerical distance, shown on the left and right axes, respectively, with *M* ± *SEM*. Annotations show the unstandardized linear slope *b* and its unadjusted *p* value. Slopes express percentage-point changes in intrusion rate per unit increase in distance. Behavioral panels include 29 participants.

#### Substitution decreases with numerical distance

We next examined whether intrusion rates varied with target location and target–PTD numerical distance. No intrusion type differed significantly between left- and right-target trials (DI_D: *p*_Holm_ = .670; DI_TD: *p*_Holm_ = .670; DI_DT: *p*_Holm_ = .670; **Figure 1—figure supplement 1A–C**). Greater numerical distance was associated with lower DI_D rates (*b* = −1.00, *SE* = 0.16, *t*(28) = −6.27, *p*_Holm_ < .001, 95% CI [−1.41, −0.59], *d* = −0.78; **Figure 1C**). The corresponding linear trends were not significant for DI_TD (*b* = 0.46, *SE* = 0.45, *t*(28) = 1.01, *p*_Holm_ = .641, 95% CI [−0.70, 1.61], *d* = 0.12; **Figure 1C**) or DI_DT (*b* = −0.05, *SE* = 0.20, *t*(28) = −0.23, *p*_Holm_ = .817, 95% CI [−0.57, 0.47], *d* = −0.03; **Figure 1—figure supplement 1D**).

Trial-level binomial GLMMs examined location and distance in separate models for each intrusion type (**Figure 1—figure supplement 2**; **Table S2**). Consistent with the participant-level analysis, **greater** distance was associated with lower odds of DI_D (*b* = −0.18, *SE* = 0.03, *z* = −6.28, *p*_Holm_ < .001, OR = 0.84, 95% CI [0.78, 0.90]; **Figure 1—figure supplement 2D**), whereas distance effects were not significant for DI_TD or DI_DT (**both** *p*_Holm_ = .529; **Figure 1—figure supplement 2E–*F***). The location model indicated lower odds of DI_DT on right-than left-target trials (*b* = −0.41, *SE* = 0.17, *z* = −2.49, *p*_Holm_ = .038, OR = 0.66, 95% CI [0.44, 0.98]; **Figure 1—figure supplement 2C**), although the participant-level comparison was not significant. Location effects were not significant for DI_D (*p*_Holm_ = .148) or DI_TD (*p*_Holm_ = .381; **Figure 1—figure supplement 2A–B**).

#### Accuracy differs across posttarget conditions

Overall task accuracy differed across conditions (*F*(1.65, 46.17) = 64.10, *p*_GG_ < .001, ω^2^ = 0.46). Accuracy was highest in T (*M* = 93.60, *SD* = 6.42), followed by TT (*M* = 72.09, *SD* = 17.23) and TD (*M* = 54.87, *SD* = 23.15). Accuracy was higher in T than TT (*t*(28) = 8.29, *p*_Holm_ < .001, 95% CI [14.90, 28.11], *d* = 1.25) and TD (*t*(28) = 10.96, *p*_Holm_ < .001, 95% CI [29.73, 47.73], *d* = 2.25), and higher in TT than TD (*t*(28) = 4.30, *p*_Holm_ < .001, 95% CI [7.03, 27.42], *d* = 1.00). TT trials also produced order reversals (*M* = 11.66, *SD* = 8.97). Averaged across participants, order reversals accounted for 41.82% of TT errors, whereas the three intrusion types together accounted for 91.22% of TD errors. These outcomes therefore accounted for substantial proportions of errors in their respective conditions.

Thus, concomitant target–PTD reports were the most frequent intrusion outcome. Substitution reports were less frequent at greater numerical distances, whereas no reliable linear distance effect was detected for either concomitant report order.

### ERP Signatures of Different Intrusion Outcomes and Posttarget Stimuli

We compared correct TD with DI_TD and DI_D to examine neural differences between correct and intrusion reports on TD trials, in which a colored target digit was immediately followed by a gray PTD. Correct T and TT were compared to characterize processing when the posttarget item was another target. Analyses assessed component amplitudes, N2pc onset latency, and waveform differences. We additionally examined DI_DT, compared correct T, TD, and TT jointly, and used DTW and single-trial models to assess timing and trial-level associations.

#### Concomitant intrusions show no reliable averaged ERP differences

Correct TD and DI_TD trials did not differ significantly in N2pc, Pd, P3b, LPC, or CDA amplitude (all *ps* ≥ .298, all *p*_Holm_ = 1.000; *BF*_01_ = 3.04–4.89), or N2pc onset latency (*p* = .823; **Figure 2—figure supplement 1**; **Table S3**). Waveform permutation tests also detected no significant clusters (**Figure 2A–B**). The supplementary comparison of correct TD and DI_DT likewise revealed no significant amplitude differences (all *ps* ≥ .202, all *p*_Holm_ = 1.000; *BF*_01_ = 1.94–4.11) or onset-latency difference (*p* = .568; **Figure 2—figure supplement 2**; **Table S4**), and no significant waveform clusters (**Figure 2—figure supplement 3**). Thus, neither concomitant report order was associated with a reliable change in the measured condition-averaged ERP amplitudes or N2pc onset.

**Figure 2.**
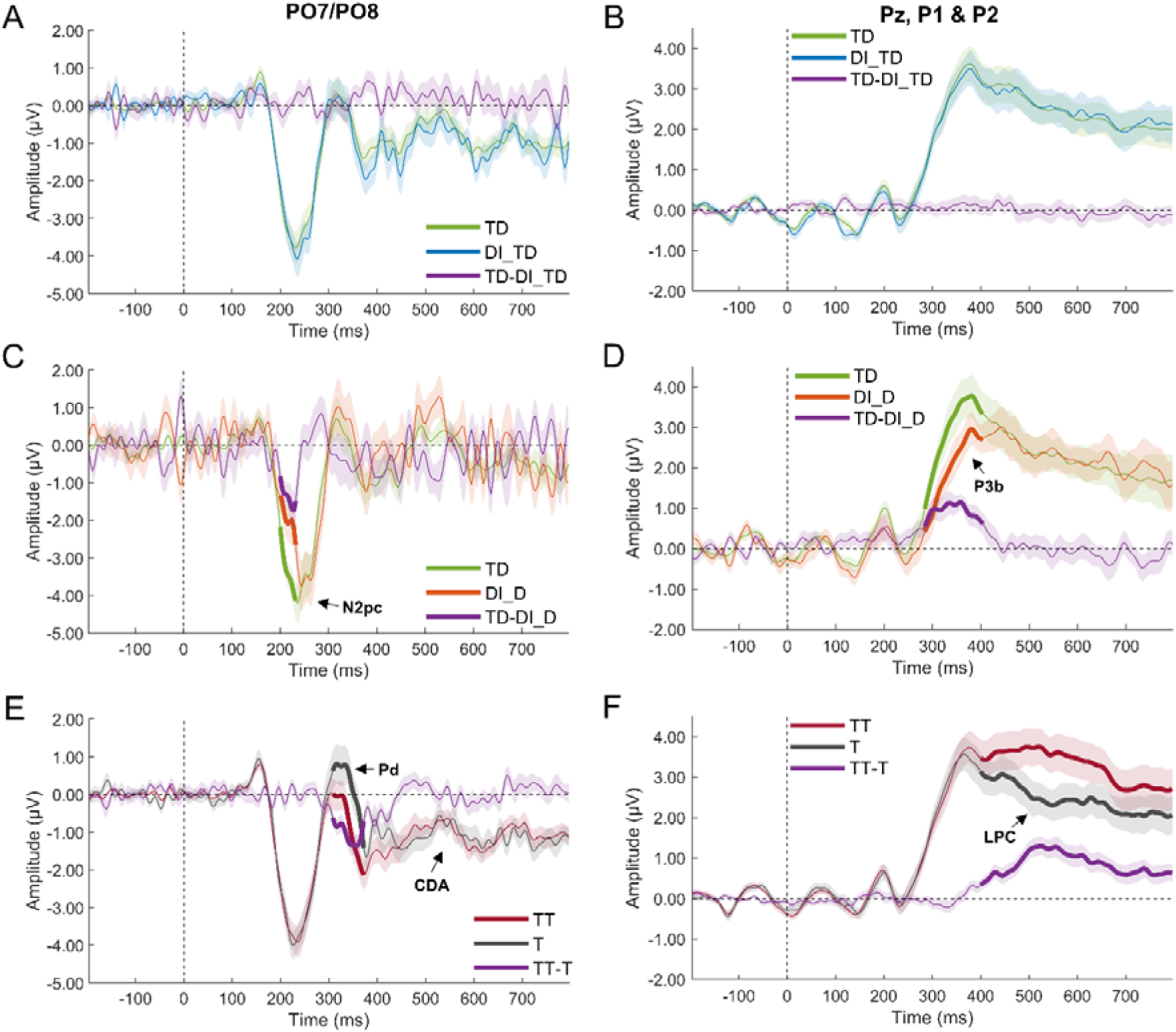
ERP correlates of distractor intrusion and posttarget processing. Grand-average event-related potentials (ERPs) for correct TD versus DI_TD (**A–B**; *n* = 29), correct TD versus DI_D (**C–D**; *n* = 16), and correct TT versus T (**E–F**; *n* = 29). Left-column panels show contralateral-minus-ipsilateral PO7/PO8 waveforms used to measure N2 posterior contralateral (N2pc), distractor positivity (Pd), and contralateral delay activity (CDA). Right-column panels show Pz/P1/P2 averages used to measure P3b and the late positive complex (LPC). Purple lines show condition differences in the order indicated by each legend. Shading indicates ± *SEM*; thick colored segments mark significant waveform clusters. Arrows identify ERP components. Vertical and horizontal dashed lines mark T1 onset and zero amplitude, respectively.

#### Substitution intrusions show delayed N2pc onset and reduced N2pc and P3b amplitudes

Correct TD elicited a more negative N2pc (*M* = −3.54 μV, *SD* = 1.67) than DI_D (*M* = −2.67 μV, *SD* = 1.66; *t*(15) = −2.95, *p* = .010, *p*_Holm_ = .039, *d* = −0.74; **Figure 2—figure supplement 4A**; **Table S5**), with a significant waveform cluster at 200–238 ms (*p*_cluster_ = .012; **Figure 2C**). N2pc onset was earlier for correct TD (*M* = 222.60 ms) than DI_D (*M* = 230.90 ms; *t*_adj_(15) = −3.52, *p* = .003, 95% CI [−13.32, −3.28], *d* = −0.88; **Figure 2—figure supplement 4F**). Correct TD also elicited a larger P3b (*M* = 3.17 μV, *SD* = 1.96) than DI_D (*M* = 2.43 μV, *SD* = 1.50; *t*(15) = 3.35, *p* = .004, *p*_Holm_ = .022, 95% CI [0.09, 1.39], *d* = 0.84; **Figure 2—figure supplement 4C**), with a significant waveform cluster at 284–404 ms (*p*_cluster_ = .008; **Figure 2D**). Pd, LPC, and CDA amplitudes did not differ significantly (all *ps* ≥ .320, all *p*_Holm_ = .959; *BF*_01_ = 2.48, 3.81, and 2.72, respectively; **Figure 2—figure supplement 4B, D–E**). Substitution reports were therefore associated with changes in N2pc amplitude and onset and with reduced P3b amplitude.

#### Posttarget input modulates Pd and LPC

Correct T and TT showed a different pattern of component differences. T elicited a larger Pd (*M* = −0.12 μV, *SD* = 2.12) than TT (*M* = −1.01 μV, *SD* = 1.87; *t*(28) = 5.90, *p* < .001, *p*_Holm_ < .001, 95% CI [0.50, 1.26], *d* = 0.45), with a significant waveform cluster at 308–372 ms (*p*_cluster_ < .001; **Figure 2E**). TT elicited a larger LPC (*M* = 3.44 μV, *SD* = 2.39) than T (*M* = 2.45 μV, *SD* = 2.03; *t*(28) = 4.68, *p* < .001, *p*_Holm_ < .001, 95% CI [0.45, 1.52], *d* = 0.46), with a significant waveform cluster at 400–800 ms (*p*_cluster_ < .001; **Figure 2F**). N2pc, P3b, and CDA amplitudes did not differ significantly (all *ps* ≥ .136, all *p*_Holm_ ≥ .273; *BF*_01_ = 5.06, 1.78, and 4.87, respectively), nor did N2pc onset latency (*p* = .785; **Figure 2— figure supplement 5**; **Table S6**).

The joint comparison of correct T, TD, and TT extended these findings to a digit PTD (**Figure 2—figure supplement 5**; **Table S6**). Condition effects were significant for Pd (*F*(1.65, 46.18) = 12.47, *p*_GG_ < .001, ω^2^ = 0.03) and LPC (*F*(1.50, 41.90) = 22.76, *p*_GG_ < .001, ω^2^ = 0.04). In addition to the T–TT differences, TD elicited a larger Pd than TT (*p* = .009, *p*_Holm_ = .018), and TT elicited a larger LPC than TD (both *p* and *p*_Holm_ < .001). The P3b difference between TD and TT did not survive Holm correction (*p* = .020, *p*_Holm_ = .060). Omnibus condition effects were not significant for N2pc (*p* = .666), P3b (*p*_GG_ = .129), or CDA amplitude (*p* = .102), or for N2pc onset latency (*p* = .426). Across correct reports, the clearest differences associated with posttarget input therefore involved Pd and LPC.

#### DTW identifies later N2pc and P3b timing in substitution intrusions but no reliable coupling

To complement the N2pc onset analysis, we used DTW to compare the timing of the full N2pc and P3b waveforms within their analysis windows and assess whether their timing variations were coupled. DI_D showed later N2pc and P3b timing than correct TD (*p* = .025 and .001, respectively; **Figure 2—figure supplement 6G–H; Table S7A).** These tests evaluated signed DTW area; descriptive median path displacements were 8.00 and 32.00 ms, respectively. Neither DI_TD (N2pc: *p* = .539; P3b: *p* = .659) nor DI_DT (N2pc: *p* = .874; P3b: *p* = .364) showed a reliable timing difference (**Figure 2—figure supplement 6A–B, D–E**; **Table S7A**). Within each comparison sample, N2pc–P3b coupling was not reliably detected for correct TD or any intrusion condition, and coupling strength did not differ reliably between correct TD and any intrusion type. All within-condition Pearson and Spearman correlations and their between-condition differences had 95% CIs spanning zero (**Figure 2—figure supplement 6C, *F*, I**; **Table S7B–C**).

#### Single-trial ERPs show component-specific associations with report outcomes

Single-trial GLMMs revealed component-specific associations with report outcomes (**Figure 2—figure supplement 7**; **Table S8**). None of the five amplitudes was reliably associated with DI_TD rather than correct TD responses (all *ps* ≥ .052, all *p*_Holm_ ≥ .260; **Figure 2—figure supplement 7B**; **Table S8B**). For DI_D, a more positive N2pc was associated with greater intrusion odds (*b* = 0.16, *SE* = 0.04, *z* = 3.76, *p* < .001, *p*_Holm_ < .001, OR = 1.17, 95% CI [1.08, 1.27]), whereas a larger P3b was associated with lower odds (*b* = −0.15, *SE* = 0.05, *z* = −3.39, *p* < .001, *p*_Holm_ = .003, OR = 0.86, 95% CI [0.79, 0.94]; **Figure 2—figure supplement 7D**; **Table S8D**). For DI_DT, a larger P3b was associated with lower intrusion odds (*b* = −0.12, *SE* = 0.05, *z* = −2.50, *p* = .012, *p*_Holm_ = .049, OR = 0.89, 95% CI [0.81, 0.98]), as was a more positive CDA voltage (*b* = −0.16, *SE* = 0.05, *z* = −3.31, *p* < .001, *p*_Holm_ = .005, OR = 0.85, 95% CI [0.77, 0.94]; **Figure 2—figure supplement 7C**; **Table S8C**). Among correct trials, a larger Pd was associated with lower TT odds relative to T (*b* = −0.15, *SE* = 0.02, *z* = −6.65, *p* < .001, *p*_Holm_ < .001, OR = 0.86, 95% CI [0.82, 0.90]), whereas a larger LPC was associated with higher TT odds (*b* = 0.23, *SE* = 0.02, *z* = 10.02, *p* < .001, *p*_Holm_ < .001, OR = 1.26, 95% CI [1.20, 1.32]; **Figure 2—figure supplement 7A; Table S8A).**

Because numerical distance predicted DI_D behaviorally, follow-up GLMMs tested whether distance moderated the ERP associations, with target location also included. No component-by-distance interaction was significant (N2pc, Pd, P3b, LPC, and CDA: χ^2^(1) = 0.08–1.09, all *p*_Holm_ = 1.000; **Figure 2—figure supplement 8**; **Table S9**). These analyses therefore provided no reliable evidence that the ERP–DI_D associations varied with numerical distance.

In summary, substitution reports were associated with attenuated and later N2pc activity, together with a smaller P3b and later P3b waveform timing. At the group level, neither concomitant intrusion type (DI_TD or DI_DT) showed reliable differences from correct TD in the measured ERP amplitudes or timing. Across correct reports, Pd was larger for T and TD than TT, and LPC was larger for TT than either T or TD.

### Distributed EEG Patterns Distinguish Intrusion Outcomes and Relate Them to Target Processing

Time-resolved decoding tested whether distributed EEG activity distinguished intrusion outcomes from correct processing. Comparisons with correct T and TT then assessed whether intrusion-related activity resembled processing of one or two genuine targets, using pairwise decoding, the T–TT reference axis, and PCM.

#### Different reports within the TD condition are decodable

These trials all presented a colored target digit immediately followed by a gray PTD and were grouped by the subsequent report. Correct TD and DI_TD were discriminable over 230–800 ms (*p*_cluster_ < .001; **Figure 3A**), despite the absence of reliable differences in the measured ERP components. This information generalized across time: classifiers trained at one time point also distinguished the conditions at other times, consistent with a relatively sustained format of discriminating information (King & Dehaene, 2014; **Figure 3—figure supplement 2B**). No significant scalp-pattern cluster was detected for this comparison (**Figure 3—figure supplement 3B**).

**Figure 3.**
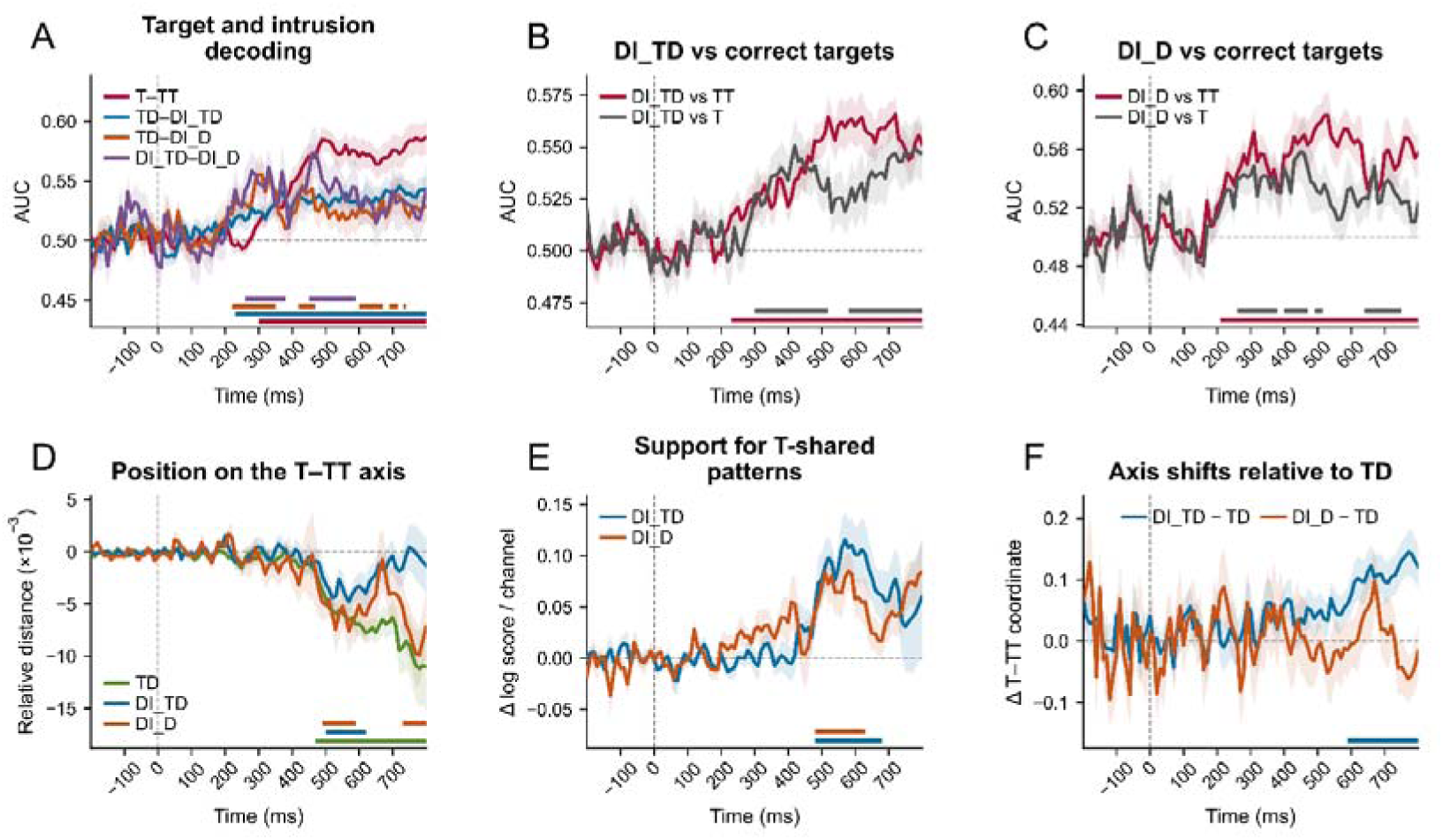
Distributed electroencephalographic patterns and their relation to target processing. **(A)** Time-resolved decoding of correct T versus TT, correct TD versus DI_TD, correct TD versus DI_D, and DI_TD versus DI_D. Performance is the area under the receiver operating characteristic curve (AUC). **(B–C)** Decoding of each intrusion type versus correct T and TT. **(D)** Crossvalidated distance to T minus distance to TT; negative values indicate relative proximity to T. **(E)** Differences in held-out predictive log scores between T-shared and TT-shared pattern component modeling (PCM) models; positive values favor T-shared patterns. **(F)** Intrusion-minus-correct-TD differences in T–TT axis coordinates; positive values indicate a TT-directed shift relative to correct TD. Shading shows ± *SEM*. Solid colored bars mark significant clusters; dashed bars mark marginal clusters (.05 ≤ *p*_cluster_ < .10). Gray dashed horizontal lines mark chance decoding or zero difference. DI_TD analyses included 29 participants; comparisons involving DI_D included 16.

Correct TD and DI_D were discriminable at 220–350 ms (*p*_cluster_ = .005), 420–470 ms, and 600–670 ms (both *p*_cluster_ = .045; **Figure 3A**). Their temporal-generalization map showed higher AUC values mainly near the diagonal (**Figure 3—figure supplement 2C**), and significant scalp-pattern clusters included central, parietal, and right frontocentral electrodes (**Figure 3—figure supplement 3C**). DI_TD and DI_D were also discriminable at 260–380 ms (*p*_cluster_ = .012) and 450–590 ms (*p*_cluster_ = .010), with generalization across times (**Figure 3—figure supplement 2D**). Thus, DI_TD was distinguishable from correct TD through distributed activity even when the selected ERP measures were not, and its temporal-generalization and scalp-pattern profiles differed descriptively from those of TD–DI_D.

#### Correct-report patterns vary with posttarget input

Correct T and TT were discriminable over 300–800 ms (*p*_cluster_ < .001; **Figure 3A**), and correct TD and TT over 210– 800 ms (*p*_cluster_ < .001; **Figure 3—figure supplement 1B**). Correct T and TD, which contained the same number of targets but differed in the posttarget distractor, were discriminable at 340– 510 ms (*p*_cluster_ = .003; **Figure 3—figure supplement 1A; Table S10**). Distributed activity therefore distinguished both the presence of a second target and the presence of a digit PTD.

#### Both intrusion outcomes differ from correct single- and two-target activity

We next asked whether substitution and concomitant intrusions differed from accurate processing of one or two genuine targets. DI_TD differed from correct TT over 230–800 ms and DI_D from correct TT over 210–800 ms (both *p*_cluster_ < .001). DI_TD differed from correct T at 300–520 ms and 580–800 ms (both *p*_cluster_ = .003; **Figure 3B**). DI_D differed from correct T at 260–380 ms (*p*_cluster_ = .006), 400–470 ms (*p*_cluster_ = .019), and 640–750 ms (*p*_cluster_ = .012; **Figure 3C**). For each intrusion type, discrimination from TT generalized across times and yielded significant scalp-pattern clusters **(Figure 3—figure supplement 2E, G and Figure 3—figure supplement 3E, G).** Discrimination of DI_TD from T also generalized across times **(Figure 3—figure supplement 2F);** discrimination of DI_D from T showed more restricted generalization and significant scalp-pattern clusters (**Figure 3—figure supplement 2H and Figure 3—figure supplement 3H**; **Table S10**). Matching the number of reported identities therefore did not yield indistinguishable activity: DI_TD differed from correct TT, and DI_D from correct T.

#### Intrusion activity is relatively closer to single-target activity at middle-to-late times

Pairwise decoding establishes discriminability but does not determine which reference a condition more closely resembles. We therefore used the crossvalidated T–TT reference axis to test whether DI_TD was closer to correct TT and DI_D to correct T (**Figure 3D; Table S11**). The distance-to-T minus distance-to-TT contrast was negative for DI_TD at 500–620 ms (*p*_cluster_ = .007), and for DI_D at 490–590 ms (*p*_cluster_ = .016) and 730–800 ms (*p*_cluster_ = .032). Correct TD showed the same direction over 470–800 ms (*p*_cluster_ < .001). Both intrusion outcomes were therefore relatively closer to T during these intervals, including DI_TD despite its two-identity report.

PCM provided converging evidence for this relative relationship (**Figure 3E**). The T-shared model predicted held-out data better than the TT-shared model for DI_TD at 480–680 ms (*p*_cluster_ = .008) and DI_D at 480–630 ms (*p*_cluster_ = .027). The TD-shared model also outperformed the TT-shared model, whereas T-shared and TD-shared models were not reliably differentiated (**Figure 3—figure supplement 4A–B**; **Table S12**). Model comparisons therefore favored pattern sharing with correct T or TD over correct TT.

#### Both concomitant intrusion types show late TT-directed shifts relative to correct TD

For DI_TD, relative proximity to T coexisted with a later shift toward TT compared with correct TD: the DI_TD-minus-TD axis-coordinate contrast was positive over 590–800 ms (*p*_cluster_ < .001; **Figure 3F; Table S11**). DI_DT showed a similar directional change in the cross-condition classifier analysis. The classifier trained on correct T and TT assigned higher TT-directed scores to DI_DT than correct TD at 650–700 ms (*p*cluster = .034), with a further marginal cluster at 730–770 ms (*p*_cluster_ = .076; **Figure 3—figure supplement 5C; Table S14**). No significant axis-coordinate difference was detected for DI_D versus correct TD.

#### Other DI_DT analyses showed no consistent preference for either target reference

The proportion of DI_DT trials classified as TT did not reliably differ from 50% (**Figure 3— figure supplement 5B**), and the reference-axis, RSA, and direct PCM contrasts provided no reliable preference for T or TT (**Figure 3—figure supplement 5D–F**; **Table S14**). Thus, DI_DT showed a late TT-directed change relative to correct TD without a consistent preference for the TT reference.

Together, distributed EEG patterns distinguished correct processing of different posttarget inputs and separated DI_TD and DI_D from correct TD. Both intrusion outcomes were relatively closer to correct T at middle-to-late times, with PCM also favoring sharing with T or TD over TT. DI_TD nevertheless showed a later TT-directed shift relative to correct TD, and DI_DT showed a related late classifier-score change.

### Reduced Frontal Theta Power and Lateralization in Target–Distractor Reports

Time–frequency analyses examined whether oscillatory activity distinguished DI_TD from correct TD. We first assessed regional power and its phase-locking properties, then frontal lateralization and single-trial associations. Comparisons with other intrusion outcomes and correct-report conditions assessed the broader distribution of these effects.

#### Reduced induced left frontal theta power in DI_TD

Correct TD elicited greater lFC power than DI_TD in a cluster concentrated predominantly in theta and extending into alpha frequencies (*p*_cluster_ = .004, *p*_Holm_ = .031; **Figure 4A**). The cluster spanned 4–12 Hz and 232–928 ms at its outer bounds. No clusters in the other regions survived correction (all *p*_Holm_ ≥ .410; **Figure 4—figure supplement 1**). At the single-trial level, greater lFC power within this cluster was associated with lower odds of DI_TD rather than correct TD responses (*b* = −0.10, *SE* = 0.03, *z* = −3.78, *p* < .001, OR = 0.91, 95% CI [0.86, 0.95]; **Figure 4C**; **Table S15**).

**Figure 4.**
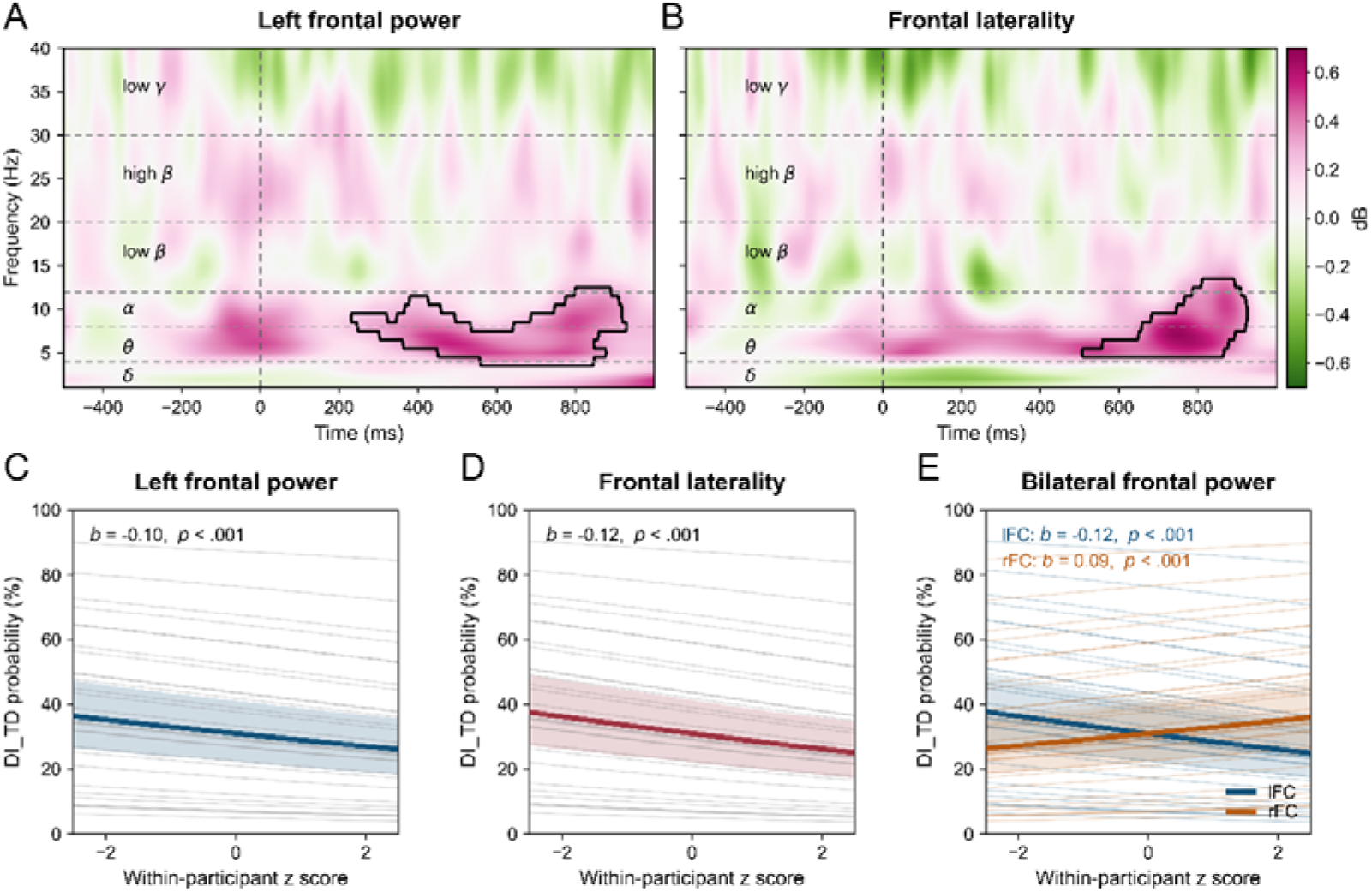
Frontal oscillatory power and lateralization are associated with distractor intrusion. (**A**) Correct-TD-minus-DI_TD power at left frontal electrodes (lFC: F1, F3, F5, F7, FC1, FC3, FC5). (**B**) Condition difference in frontal lateralization: (left − right)_correct TD_ − (left − right)_DI_TD_, using F3, F5, F7, FC3, FC5 and their right frontal (rFC) counterparts. Power is expressed in decibels (dB) relative to the −500 to −200 ms baseline. Magenta indicates positive differences and green negative differences; black outlines enclose significant clusters. Dashed lines mark T1 onset and frequency-band boundaries. (**C–E**) Single-trial binomial generalized linear mixed-effects model (GLMM) predictions for DI_TD versus correct TD (*n* = 29). Panel C uses the cluster in A; D and E use the cluster in B. D includes left-minus-right power, bilateral mean power, and target location; E includes separate lFC and rFC predictors and target location. Predictors are centered within participant and expressed in pooled within-participant *SD* units, with a common scale for both hemispheres in E. Thick curves and shading show fixed-effect predictions and 95% confidence intervals (CIs); thin curves show participant-specific predictions. Bilateral mean power in D and the other hemisphere’s power in E are held at zero. Predictions in D and E are averaged over target location. Displayed *b* values are log-odds coefficients; *p* values are from unadjusted two-sided Wald tests.

The induced-power analysis identified an overlapping cluster in which correct TD exceeded DI_TD, predominantly in theta and adjacent frequencies (outer bounds: 4–19 Hz and 236–928 ms; *p*_cluster_ = .006; **Figure 4—figure supplement 2A**). No significant clusters were detected for lFC evoked power, lFC ITPC, or ITPC averaged across all analyzed electrodes (all *p*_cluster_ ≥ .168; **Figure 4—figure supplement 2B–D**). The frontal power difference was therefore accompanied by an induced-power difference, with no reliable difference detected in the phase-locked measures.

#### Reduced frontal lateralization in DI_TD

The correct-TD-minus-DI_TD power difference was greater over left than right frontal electrodes in a significant condition-by-hemisphere cluster (*p*_cluster_ = .030; **Figure 4B**), involving mainly theta and adjacent alpha frequencies (outer bounds: 5–13 Hz and 508–924 ms). Greater single-trial left-minus-right power within this cluster was associated with lower DI_TD odds after controlling for bilateral mean power and target location (*b* = −0.12, *SE* = 0.03, *z* = −4.63, *p* < .001, OR = 0.89, 95% CI [0.85, 0.93]; **Figure 4D**; **Table S15**). An equivalent model with separate left and right predictors associated greater lFC power with lower DI_TD odds (*b* = −0.12, *SE* = 0.03, *z* = −4.38, *p* < .001, OR = 0.89, 95% CI [0.84, 0.94]) and greater rFC power with higher odds (*b* = 0.09, *SE* = 0.03, *z* = 3.31, *p* < .001, OR = 1.10, 95% CI [1.04, 1.16]; **Figure 4E**). Adding rFC power improved model fit beyond lFC power and target location (χ**^2^**(1) = 10.93, *p* < .001); adding lFC power likewise improved fit beyond rFC power and target location (χ**^2^**(1) = 19.22, *p* < .001; **Table S16**). Each hemisphere therefore contributed information about report outcome beyond the other.

#### Control analyses of intrusion type and posttarget processing

We first tested whether the frontal effects extended to other intrusion outcomes. Neither TD–DI_DT nor TD–DI_D showed frontal power or lateralization differences that survived correction (**Figure 4—figure supplement 3A–F; Table S17)**.

We next compared correct T, TD, and TT to assess differences associated with target number and PTD presence. None of these contrasts reproduced the frontal theta pattern that remained significant after correction (**Figure 4—figure supplement 4**; **Table S17**). Instead, TT–T showed a late, beta-centered increase in the all-electrode average (*p*_cluster_ = .0039; **Figure 4—figure supplement 5A**), with increases over rFC and centroparietal regions surviving regional correction (**Figure 4—figure supplement 5B, D**). The observed theta pattern was therefore not reproduced by the correct-report contrasts for target number or PTD presence.

Finally, comparing DI_TD with correct TT revealed a left frontal power difference (*p*_cluster_ = .0025, *p*_Holm_ = .017; **Figure 4—figure supplement 3G**; **Table S17**). The corresponding frontal-lateralization comparison was not significant after correction (**Figure 4—figure supplement 3I**). T–DI_TD showed no corrected-significant frontal power or lateralization difference (**Figure 4—figure supplement 3J–L**). Thus, reduced left frontal power also characterized DI_TD relative to correct two-target reports, whereas the reliable lateralization difference was observed relative to correct TD.

Thus, DI_TD was associated with reduced left frontal power predominantly in theta, an overlapping induced-power difference, and reduced left-relative-to-right frontal power. The left frontal difference occurred both relative to TD, which had the same actual target count, and relative to TT, which had the same reported identity count. This pattern was therefore not explained solely by whether one or two targets were presented or one or two identities were reported.

### Joint Neural Correlates of Intrusion Reports

To examine neural associations with intrusion reports while accounting for covariation among the measures, we fitted a multinomial GLMM jointly including five ERP amplitudes, lFC power, and rFC power, with numerical distance and target location as covariates. Correct TD served as the reference category (*n* = 29; **Figure 5**; **Table S18A**). More positive N2pc voltage was associated with greater DI_D odds (*b* = 0.17, *p* < .001, *p*_Holm_ = .014, OR = 1.19, 95% CI [1.08, 1.31]). Larger P3b amplitude was associated with lower DI_D odds (*b* = −0.37, both *p* and *p*_Holm_ < .001, OR = 0.69, 95% CI [0.62, 0.78]), whereas larger LPC amplitude was associated with greater odds (*b* = 0.32, both *p* and *p*_Holm_ < .001, OR = 1.38, 95% CI [1.23, 1.55]). Matched-trial sensitivity analyses showed that this LPC association was reliable only after P3b adjustment; omitting P3b yielded a nonsignificant LPC coefficient (*b* = 0.055, *p* = .181; **Table S19**). Greater numerical distance was also associated with lower DI_D odds (*b* = −0.31, both *p* and *p*_Holm_ < .001, OR = 0.73, 95% CI [0.67, 0.80]; **Figure 5B**).

**Figure 5.**
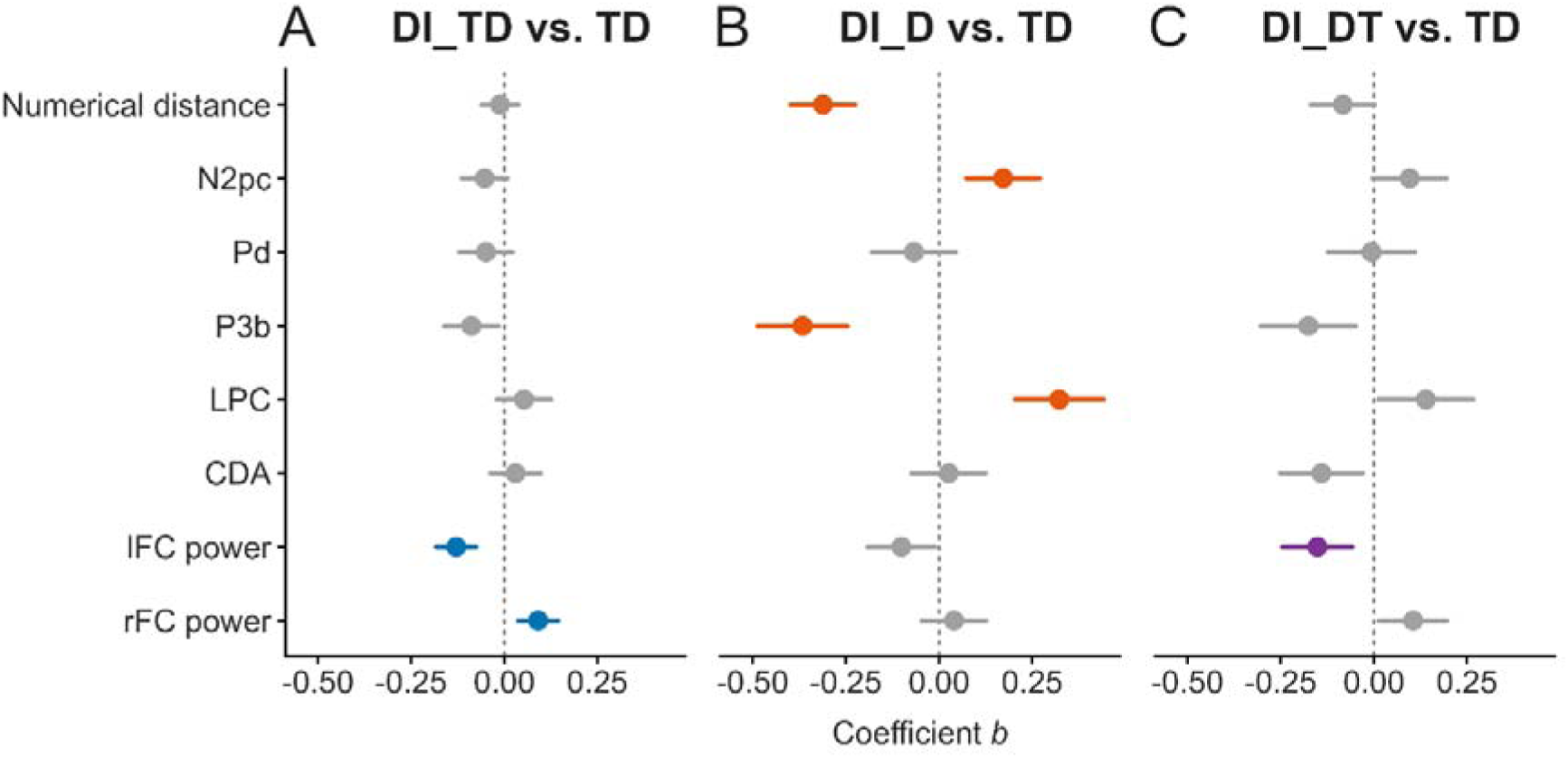
Joint ERP and frontal-power associations with intrusion outcomes. Multinomial GLMM coefficients for DI_TD **(A)**, DI_D **(B)**, and DI_DT **(C)**, each relative to correct TD (*n* = 29). All seven neural predictors, numerical distance, and target location entered the same model; location is controlled but not shown. Neural effects are per pooled within-participant *SD*; distance effects are per full-sample *SD*. The lFC feature uses the regional time–frequency mask; the rFC feature uses right-sided power from the frontal-lateralization mask. Points show *b* and lines show unadjusted 95% CIs. Colored estimates survive Holm correction across all 27 slopes; gray estimates do not. The dashed vertical line marks zero. Full statistics are reported in **Table S18A**.

Frontal power was associated with concomitant intrusions after accounting for ERP amplitudes and the behavioral covariates. Greater lFC power was associated with lower odds of both DI_TD (*b* = −0.13, both *p* and *p*_Holm_ < .001, OR = 0.88, 95% CI [0.83, 0.93]) and DI_DT (*b* = −0.15, *p* = .002, *p*_Holm_ = .031, OR = 0.86, 95% CI [0.78, 0.94]) relative to correct TD. Greater rFC power was additionally associated with higher DI_TD odds (*b* = 0.09, *p* < .001, *p*_Holm_ = .018, OR = 1.10, 95% CI [1.04, 1.15]; **Figure 5A, C**). No ERP coefficient for either concomitant intrusion survived correction (**Table S18A**).

The 11-participant intersection-sample analysis retained the DI_D associations with P3b, LPC, and numerical distance, the DI_TD associations with lFC and rFC power, and the DI_DT association with lFC power. The N2pc association with DI_D did not survive correction in this subset (*p* = .045, *p*_Holm_ = .712; **Figure 5—figure supplement 1**; **Table S18B**). Together, the joint analysis identified ERP and numerical-distance associations with substitution intrusions and frontal-power associations with concomitant intrusions.

## Discussion

Combining ERP, multivariate pattern, and time–frequency analyses, the present study examined how attentional episodes develop with subsequent input and how different forms of distractor intrusion relate to this process when target number is uncertain. First, differences among correct-report conditions emerged primarily after initial attentional engagement: Pd was larger for T and TD than for TT, whereas LPC was larger for TT than for either single-target condition. Multivariate analyses likewise showed that the nature and relevance of subsequent input altered distributed neural activity. Second, relative to correct TD reports under identical stimulus conditions, DI_D was associated with smaller and later N2pc and P3b activity, linking target substitution to changes in attentional engagement and subsequent encoding. By comparison, the most frequent intrusion outcome, DI_TD, showed no corresponding reliable differences in averaged ERPs but exhibited distinguishable distributed activity and lower left-frontal low-frequency power. Moreover, both intrusion forms were distinguishable from correct single- or dual-target conditions with the same number of reported identities. Reporting one or two identities therefore did not simply correspond to correctly processing one or two actual targets. These findings suggest that accounts of temporal selection must explain both why targets sometimes go unreported and why irrelevant identities can enter report even when the target is preserved.

### Posttarget Stimuli Shape the Dynamics of Attentional Episodes

Comparisons among the three correct-report conditions located the main differences in attentional episodes after initial engagement, during subsequent regulation and sustained processing. N2pc amplitude and onset latency did not differ reliably, consistent with related findings by Tan and Wyble (2015) and Callahan-Flintoft et al. (2018). Akyürek et al. (2017), however, reported a larger N2pc for two targets, suggesting dependence on RSVP settings. Our findings are compatible with attentional episode theory: Enhancement triggered by the first target can extend to subsequent input, allowing adjacent targets to be processed within one episode (e.g., Wyble et al., 2009, 2011).

The present Pd increased with the incompatibility of posttarget input with task requirements. At 300–400 ms, the Pd was larger for T and TD than for TT, with a marginal T– TD difference (*p*_Holm_ = .053). Ordinary letters matched neither the target color nor the reportable digit category and elicited the largest Pd. Gray digit PTDs retained reportable identities and elicited an intermediate Pd, whereas consecutive targets elicited the smallest Pd. An exploratory association between greater target–PTD numerical distance and larger Pd did not survive correction (*p* = .014, *p*_Holm_ = .071; **Figure 2—figure supplement 9**; **Table S20**). This provides weak evidence for sensitivity to target–PTD compatibility, although numerical distance cannot be equated with task compatibility. This gradient links posttarget relevance to regulation of enhancement after the initial selection stage.

Our Pd gradient is compatible with accounts of enhancement maintenance and termination in attentional episode and boost-and-bounce theories. Successive targets can sustain an episode, whereas the absence of further target input and encoding-related feedback can help terminate it (e.g., Wyble et al., 2009, 2011); boost-and-bounce theory emphasizes inhibitory feedback triggered by an enhanced distractor (Olivers & Meeter, 2008). Together with evidence linking the Pd to attentional termination (Sawaki et al., 2012), our findings may reflect relatively automatic termination or suppression under the current task set. Such regulation can follow stimulus processing without requiring participants to deliberately reject each distractor.

The present Pd modulation suggests that late lateralized effects of categorization matching may also involve attentional termination or suppression. In Zivony and Eimer’s (2022a) experiment with three PTD types, target identities were restricted to three letters specified for each block. Matching PTDs belonged to that set, nonmatching PTDs were letters outside it, and category-irrelevant PTDs were digits; all lacked the external shape cue defining a target. Both EEG experiments showed late matching effects, interpreted as a prolonged N2pc. Our Pd modulation at a similar stage suggests that these late negative differences could partly reflect variation in Pd amplitude across PTD types. This interpretation requires direct testing and does not establish a different component attribution. Both sustained enhancement and variation in suppression therefore remain possible contributors to the observed lateralized waveform.

Target number differentiated the LPC more consistently than the P3b in the present study. The omnibus P3b effect at 300–450 ms was nonsignificant, and the TD–TT comparison did not survive correction. At 450–700 ms, the LPC was larger for TT than for T or TD, which did not differ reliably. The cluster test identified a T–TT difference in parietal positivity at 400–800 ms (*p*_cluster_ < .001). Spanning the P3b tail and subsequent sustained positivity, this effect emphasizes later processing without assigning the difference exclusively to one component.

Our predominantly late effect of target number highlights the need to distinguish earlier P3b activity from subsequent sustained positivity. Dell’Acqua et al. (2016) and Akyürek et al. (2017) reported larger P3/P3b responses to two targets, but broad measurement ranges complicate separating encoding from maintenance. During encoding and brief retention of one to four faces, Morgan et al. (2008) found no reliable load effect on the earlier P3b, whereas the later P3b at 450–700 ms decreased with load. Although opposite in direction to our effect, this suggests different load sensitivities across time. Ruchkin et al. (1990) likewise observed a larger posterior positive slow wave without a corresponding P3b change. Gunseli, Olivers, and Meeter (2014) found a larger LPC with greater anticipated search difficulty despite constant template number, indicating sensitivity to the subsequent use of maintained information. These findings establish sensitivity of parietal positivity to load, but leave its attribution to a specific processing stage open.

The more consistent LPC effect in our study supports a distinction in the relative functional contributions of P3b and LPC. Earlier P3b activity may emphasize encoding, binding, and updating within an episode, whereas later LPC activity may emphasize maintenance and continued use. Attentional episode theory allows consecutive targets to share enhancement without encoding activity increasing additively with their number (e.g., Wyble et al., 2009, 2011). Related work links P3b activity to working-memory binding and updating (e.g., Craston et al., 2009; Polich, 2007). Its amplitude may depend on the strength of selected identity representations and the organization of their encoding. Our larger LPC for TT is compatible with the additional demands of retaining two targets until report. This interpretation concerns relative contributions, without assuming sharply separated stages or an exclusive maintenance function for LPC.

The present CDA did not reliably distinguish one- and two-target processing at PO7/PO8. Supplementary P7/P8 and combined-electrode analyses likewise revealed no reliable differences within the CDA window (**Figure 2—figure supplement 10**; **Table S21**). This differs from Akyürek et al.’s (2017) late set-size effect, although their targets were abstract square-corner symbols and ours were familiar digits. Other studies found no reliable CDA difference between two and four complex stimuli, or among one, two, and four orientations under severely restricted presentation times (Gao et al., 2009; Hao et al., 2018). These findings suggest dependence on processing requirements and encoding conditions, without directly explaining our null result. The stimulus difference identifies a possible boundary condition, although it cannot by itself account for the discrepancy.

Our multivariate analyses distinguished distributed activity patterns among the three correct-report conditions. T and TT were discriminable at 300–800 ms, TD and TT at 210–800 ms, and T and TD at 340–510 ms, indicating differences across electrode activity patterns. Multivariate methods can extract working-memory load information (Adam et al., 2020), but our conditions also differed in posttarget color and category. Decoding therefore distinguishes overall processing conditions and cannot directly identify representations of target number or identity. This interpretation is consistent with condition information being distributed across electrodes, beyond effects measured by individual ERP components.

Our higher late beta power for TT than for T at approximately 700–1000 ms is compatible with additional retention and internal processing demands. This difference occurred after sequence offset and before the report prompt. Beta load effects depend on task: Palva et al. (2011) observed increasing frontoparietal beta amplitude during retention. However, a systematic review documented substantial heterogeneity (Pavlov & Kotchoubey, 2022). Our late beta increase thus provides complementary evidence of greater subsequent processing demands, without constituting a task-general index of target number.

The main differences in Pd, LPC, and late beta power outline how attentional episodes develop with subsequent input. Following initial engagement, posttarget relevance may regulate the continuation, termination, or suppression of enhancement. Selected information then undergoes encoding, binding, and updating, with subsequent maintenance and continued use guided by report requirements.

### Attentional Engagement and Identity Competition in Substitution Intrusions

The present association of DI_D with attenuated and delayed N2pc activity extends the link between delayed attentional engagement and target substitution to situations in which target number is uncertain. Relative to correct TD reports, DI_D showed an approximately 8-ms delay in N2pc onset and a smaller amplitude, with a reliable waveform cluster at 200–238 ms. This pattern agrees with findings from single-target studies (Zivony & Eimer, 2020, 2021a). In the delayed-engagement account (Zivony & Eimer, 2021a, 2022b), enhancement triggered by the target-defining feature acts on the sensory representations available at that time. When engagement is delayed, backward masking may already have weakened the target representation, whereas the subsequent PTD can receive more enhancement and gain an advantage in competition for working-memory access. Observing this pattern with intermixed one- and two-target trials indicates that uncertainty about target number does not eliminate the association between substitution and altered engagement.

Our smaller and later P3b, together with decreasing DI_D rates at greater target–PTD numerical distances, further implicates identity competition and subsequent encoding in substitution intrusions. P3b amplitude was lower for DI_D than for correct TD, with a waveform cluster at 284–404 ms; DTW also indicated later P3b waveform timing. The association between smaller P3b amplitude and higher DI_D odds remained after jointly controlling for other ERPs, frontal power, and behavioral covariates. Given links between P3b and encoding, binding, and updating, these findings are compatible with altered encoding-related processing following weaker target enhancement (e.g., Meng et al., 2023; Polich, 2007; Xu et al., 2026). Targets and PTDs at smaller numerical distances were also more likely to produce substitution reports, and this association remained in the joint model, suggesting that identity discriminability should be incorporated into the competition account (Zivony & Eimer, 2020). However, numerical distance was not separated from visual similarity or report-related factors, and no reliable distance-by-ERP interaction emerged. The processing stage underlying the distance effect therefore remains unresolved, and P3b alone cannot identify whether the target or PTD was ultimately encoded.

Although both N2pc and P3b were delayed in DI_D, we detected no reliable temporal coupling in correct TD or any of the three intrusion conditions, nor reliable differences in coupling strength between correct TD and any intrusion condition. Reanalyzing single-target intrusion data, Hosseini et al. (2024) found correlated N2pc and P3 timing within both correct and intrusion reports, with a weaker relationship during intrusions, providing evidence for a temporal link between engagement and subsequent encoding. We did not obtain comparable coupling evidence, indicating that average delays in both components do not establish stable propagation of delay across processing stages. Uncertainty about target number may introduce additional variability after initial engagement because participants must also determine whether a second target occurred. Category-expectation research has shown that expectations can modulate P3b without corresponding N2pc changes, providing indirect evidence that additional factors can influence later processing (Zivony & Eimer, 2024). However, category expectations differ from uncertainty about target number, and this explanation requires a direct manipulation of target-number certainty.

Our multivariate findings further show that DI_D differs from correct single-target processing despite involving a report of only one identity. Correct TD and DI_D were discriminable at 220–350 ms and during some later intervals, and DI_D also differed from correct T, indicating processing differences around initial selection and at later stages. Conversely, middle-to-late relative-distance and pattern-component analyses favored shared activity with T or TD over TT. DI_D therefore retained some features of single-target processing alongside changes associated with target substitution. Because the reference conditions also differed in stimulus features and task relevance, this sharing cannot be equated directly with identical identity representations or encoding processes.

The present findings support the delayed-engagement account of substitution intrusions and extend it to uncertainty about target number. DI_D was associated with attenuated and delayed N2pc activity, a smaller and later P3b, and a lower occurrence rate at greater target–PTD numerical distances. Explaining why a distractor replaces the target therefore requires considering the timing of engagement, the relationship between competing identities, and subsequent encoding-related processing together.

### Frontal Theta and Lateralization in Concomitant Intrusions

Neither DI_TD nor DI_DT showed the ERP abnormalities observed for substitution intrusions. Relative to correct TD reports, neither form of concomitant intrusion showed reliable differences in N2pc amplitude or latency, and neither showed reliable differences in the mean amplitudes of the subsequent Pd, P3b, LPC, or CDA. The corresponding waveform cluster tests likewise revealed no reliable differences. Delayed attentional engagement therefore appears insufficient, on its own, to explain intrusions in which the target is retained and the PTD is also reported. Although these null effects do not establish identical processing, the findings still require a further theoretical explanation: Why is the PTD accepted as an additional target when the target identity is preserved?

Our multivariate results further indicate that concomitant intrusions involve changes in distributed activity beyond those captured by averaged ERPs. DI_TD and correct TD reports were distinguishable at approximately 230–800 ms, with cross-temporal generalization suggesting that their differences persisted over time. Comparisons with correct single- and dual-target reports did not assign DI_TD to a single processing pattern: its activity was closer to correct T during some intervals and later shifted along the TD-to-TT discriminant direction. DI_DT also showed a late shift in this direction. Reporting two identities thus need not reproduce the processing of two genuine targets. These analyses, however, distinguish condition-related activity patterns rather than directly identifying the contents or binding status of target and PTD representations.

Our clearest time–frequency finding was lower left-frontal power for DI_TD than for correct TD and TT reports, primarily in the theta band and extending into lower alpha frequencies. The difference between correct TD and DI_TD was also present in induced power, indicating that it was not confined to phase-consistent evoked activity. Control comparisons among correct T, TD, and TT reports did not reproduce the same corrected theta pattern, and substitution intrusions did not show a corresponding group-level power difference. This pattern is difficult to attribute solely to the number of physical targets or the presence of a PTD and instead points toward how information is ultimately selected for report.

Our theta associations also showed a lateralized pattern. Correct TD reports had a larger left-minus-right frontal power difference than DI_TD, predominantly in the theta band, with some extension into the alpha band, at approximately 508–924 ms; the association remained after controlling for bilateral mean power and stimulus location. In the joint model, lower left-frontal power was associated with greater odds of both DI_TD and DI_DT relative to correct TD, whereas higher right-frontal power was associated with greater DI_TD odds. Concomitant intrusions therefore appear to involve different contributions from left- and right-frontal activity, rather than a uniform bilateral reduction. These conditional associations do not establish that the two sides respectively facilitate and impair accurate selection.

Our time–frequency and lateralization results complement averaged ERPs and suggest that control, representational coordination, and task monitoring may contribute to concomitant intrusions. In general, frontal-midline theta is associated with recognizing and responding to cognitive-control demands and may communicate the need for increased control to task-relevant networks (Cavanagh & Frank, 2014). Perturbation evidence also links prefrontal theta to prioritizing working-memory contents (Riddle et al., 2020). The review by de Vries et al. (2020) further proposed that frontal theta activity may contribute to establishing and switching the priority states of working-memory representations. Regarding hemispheric differences, Vallesi (2021) reviewed evidence that setting and selecting rules or criteria preferentially recruit left lateral prefrontal regions and their networks, whereas continuously monitoring whether rules are being applied appropriately preferentially recruits right-sided networks. These accounts provide a theoretical background for considering our theta and lateralization associations in terms of control and task monitoring.

Our results suggest that concomitant intrusions may involve insufficient regulation and monitoring of information that has benefited from target-triggered enhancement. Because the PTD and target are both digits, the PTD may benefit from target-elicited enhancement despite failing to meet the color criterion for report. Correct TD performance would then require preserving the target identity while distinguishing the target from the PTD according to the color rule and preventing the PTD from acquiring reportable target status; correct TT performance would require coordinating two eligible targets. We therefore speculate that lower left-frontal theta during DI_TD may reflect insufficient recognition of control demands or insufficient recruitment of rule-guided regulation and monitoring, allowing the PTD to be accepted as an additional target while the target identity is preserved. This mechanistic hypothesis, based on the observed associations, highlights the possibility that retaining the target identity and excluding an irrelevant identity depend on different processing operations.

Our findings therefore extend the account of concomitant intrusions from the timing of attentional enhancement to the subsequent selection and coordination of information. Differences in distributed activity and frontal theta suggest that the inclusion of an irrelevant identity in report may depend on how task rules constrain information that continues to be processed. Such regulation offers a further perspective on how attentional episodes preserve relevant content while limiting irrelevant content.

## Summary

To our knowledge, this study is the first to systematically distinguish substitution from concomitant distractor intrusions within a single experimental framework and characterize their different processing profiles using complementary neural measures. Comparisons among correct T, TD, and TT reports further characterized the dynamics of target-triggered attentional episodes. These conditions showed no reliable differences in the initial N2pc, but differed in subsequent processing: Pd amplitudes were larger for T and TD than for TT, whereas LPC amplitudes were larger for TT than for either single-target condition. Distributed activity also differentiated the correct-report conditions. Together, these findings indicate that the nature of posttarget input shapes how processing develops following initial engagement.

Against these correct-report references, the two intrusion types showed different neural profiles. Substitution intrusions were associated with an attenuated and delayed N2pc and a reduced P3b, implicating changes in attentional engagement and subsequent encoding. Neither form of concomitant intrusion showed the same condition-averaged ERP abnormalities, yet concomitant intrusions were associated with differences in distributed activity or frontal power. These findings support different processing mechanisms for the two intrusion types: reporting an irrelevant identity instead of the target and accepting that identity as an additional target while preserving the target are not simply different degrees of the same selection failure. Accurate temporal selection therefore appears to depend on both target-triggered engagement and subsequent regulation and monitoring of processed information.

## Limitations and Future Directions

Our classification of intrusion types was based on final reports and therefore cannot fully determine the processing stage at which errors arose. Reporting both the target and the PTD demonstrates that the PTD identity influenced behavior, but this outcome alone cannot distinguish the contributions of feature binding, selection among memory representations, and report decisions. Similarly, associations between frontal theta and intrusion probability support a control-related account but do not establish that insufficient regulation or monitoring causes intrusions. Future studies could independently manipulate the color rule and category match and incorporate separate feature reports, target-number judgments, or confidence ratings to distinguish information processing, assignment of target status, and selection for report.

Our distinction between forward-order and reverse-order concomitant intrusions also warrants further investigation. DI_TD reports the target before the PTD, preserving their physical presentation order, whereas DI_DT reports the PTD before the target. Both preserve the same two identities, but DI_DT additionally reverses their order. Complementary analyses of the same dataset found a more negative posterior ERP response near 400 ms for DI_TD than DI_DT (Meng et al., 2026), providing an additional association with report order rather than an independent replication. The present analyses primarily characterized DI_TD, with DI_DT serving as a supplementary comparison, and did not provide a comprehensive direct test of differences between the two outcomes. Future studies should obtain sufficient observations of both forms and directly compare their behavioral and neural profiles. Manipulating the target– PTD interval and independently varying identity-report and order-report requirements could help establish whether their differences arise during information selection or during the subsequent organization of selected identities for report.

Our account of frontal theta also requires direct intervention. Future studies could use theta-frequency transcranial alternating current stimulation or rhythmic transcranial magnetic stimulation to modulate prefrontal activity and compare their effects on substitution and concomitant intrusions. Sham stimulation, control frequencies or stimulation sites, and EEG measurements before and after stimulation would help establish whether changes in intrusions accompany the intended theta modulation. Crucially, such studies should distinguish improved exclusion of the PTD from general changes in target identification or report criteria, thereby testing more directly the causal contribution of frontal theta to selective regulation.

## Constraints on Generality

Our findings primarily concern healthy young adults who are familiar with Arabic digits and can discriminate the target colors. The sample comprised 29 right-handed adults aged 19–24 years recruited from a university and its surrounding community. We expect the distinction between preserving a target identity and excluding irrelevant identities to inform comparable adults’ performance in rapid visual-report tasks because these constitute different requirements for accurate selection. This expectation has not been tested in other age groups, clinical populations, or individuals with different levels of numerical familiarity. The task used colored digit targets and gray digit PTDs in lateralized RSVP streams, with consecutive critical items presented at 100 ms per frame and single- and dual-target trials intermixed. The present evidence therefore primarily constrains accounts of temporal selection when target number is uncertain and neighboring items have reportable identities. Whether the specific intrusion rates, numerical-distance association, and timing and lateralization of neural activity generalize to fixed target-number tasks, other stimulus categories or sensory modalities, or naturally unfolding continuous events remains to be established.

## Materials and methods

### Participants

Thirty-two participants were recruited from the university and surrounding community. Three were excluded because of excessive EEG artifacts, using the criteria described below. The final sample comprised 29 participants (16 female; age range: 19–24 years, *M* = 20.36, *SD* = 1.45). No a priori power analysis was conducted. All participants were right-handed and reported normal or corrected-to-normal vision, normal color vision, and no history of psychiatric, neurological, or other major medical conditions.

EEG comparisons of correct TD reports with DI_D and DI_DT reports included 16 and 18 participants, respectively, who met the condition-specific trial-count criteria below. Related EEG experiments using the distractor intrusion paradigm analyzed final samples of 11–15 participants (Zivony & Eimer, 2020, 2021a, 2021b, 2022a). All participants provided written informed consent and received ¥60 per hour. The study was approved by the Ethics Committee of the Faculty of Psychology at Southwest University (approval no. H22049) and conducted in accordance with the Declaration of Helsinki.

### Transparency and Openness

We report the sample-size rationale, experimental manipulations, measures, and participant- and trial-exclusion criteria in the Materials and methods section. Neither the study nor its analysis plans were preregistered. The data and code sharing plan is described under Data and code availability.

### Shared dataset and related work

The behavioral and EEG recordings analyzed here were also used in a companion preprint primarily examining correctly ordered and reversed reports of two genuine targets (Meng et al., 2026). That report additionally included descriptive behavioral summaries and selected control analyses involving single-target trials and distractor intrusions. The present study focuses on substitution and concomitant distractor intrusions, with correct T, TD, and TT trials serving as reference conditions. Thus, correct-TT observations, descriptive behavior, and selected control comparisons overlap between the reports. Condition-specific artifact-related participant exclusions yielded 29 retained participants here and 27 in the companion report; the two reports do not represent independent samples or replications.

### Apparatus, Stimuli and Design

#### Task design

Participants reported, in temporal order, the identities of one or two colored digits embedded in two concurrent lateral RSVP streams. Color distinguished targets from gray distractors. Three conditions differed in the frame immediately following the first target (T1). In T, this frame contained two gray letters. In TD, a gray digit PTD appeared in the T1 stream and a gray letter in the opposite stream. In TT, a second colored target (T2) appeared in the T1 stream and a gray letter in the opposite stream. The two TT targets were different digits presented consecutively at the same location and in the same color.

Participants were informed of these stimulus properties and of the 1:4:5 ratio of T, TD, and TT trials before the experiment. Thus, one-target trials (T and TD combined) and two-target trials (TT) were equally frequent overall.

#### Apparatus and stimuli

Participants were tested individually in a sound-attenuated laboratory. Stimuli were presented against a black background on a 1,024 × 768-pixel LED monitor (60 Hz) at a viewing distance of approximately 60 cm. E-Prime 2.0 (Psychology Software Tools, Pittsburgh, PA) controlled presentation and recorded keyboard responses. Characters were approximately 1° high, and the centers of the two streams were separated by 3.5°. The stimulus set comprised digits 2–9 and 22 uppercase letters, excluding B, I, O, and Z. Digits and letters were sampled without replacement within each trial. If the letter pool was exhausted, it was replenished while preventing letter repetitions across consecutive frames. Distractors were gray (RGB: 118, 118, 118). Target color was randomly selected on each trial from blue (0, 84, 200), green (0, 176, 80), and orange (225, 125, 40).

#### Trial sequence

Each trial began with an 800-ms central fixation cross (0.75° × 0.75°), followed by 10–13 RSVP frames (**Figure 1A**). Each frame contained one character in each stream and lasted 100 ms, with no interstimulus interval. T1 appeared equally often in the left and right streams and at serial positions 5–8. The condition-specific frame appeared immediately after T1 and was followed by four distractor frames. Fixation remained visible throughout the sequence and for 500 ms after the last frame. Two successive prompts then requested the first and second colored digits. Participants responded using keys 2–9 and pressed the space bar at the second prompt if no second target had appeared. Both prompts required a response, with no response deadline. Each trial ended with an 800-ms blank interval.

#### Practice and Main-Task Structure

Participants completed 20 practice trials before the main task. The main task was designed to comprise 1,000 trials divided into ten 100-trial blocks, with breaks between blocks. The planned trial counts were 100 T, 400 TD, and 500 TT trials, presented in a randomly intermixed order. T1 location (left or right visual field) and serial position were balanced within each condition, and target colors were distributed as evenly as possible. Technical recording issues resulted in missing data for one or two blocks in three participants. Consequently, data were available for 1,000 trials in 26 of the 29 retained participants, 900 trials in two participants, and 800 trials in one participant.

### Behavioral Analysis

We compared the rates of DI_D, DI_TD, and DI_DT in the TD condition and examined whether each intrusion type varied with target location and absolute target–PTD numerical distance (1–7). Intrusion rates were expressed as percentages of all TD trials; location- and distance-specific rates used the corresponding TD trials as denominators. For each participant and intrusion type, an unstandardized linear slope was fitted across the seven distance-specific rates, weighting each distance equally. The group mean slope *b* therefore represented the percentage-point change in intrusion rate per unit increase in distance. Each intrusion type was also compared with other TD errors, defined as incorrect reports outside the three intrusion categories. Task accuracy and TT order-reversal rates were calculated relative to all trials in the respective conditions. Behavioral analyses included all available trials, irrespective of EEG artifact status.

### EEG Recording and Preprocessing

#### EEG recording

EEG was recorded continuously from 64 Ag/AgCl electrodes mounted in a Neuroscan Quik-Cap according to the extended international 10–20 system. Additional electrodes below the left eye and at the outer canthus of the right eye recorded vertical and horizontal electrooculographic activity, respectively. Signals were referenced online to an electrode near the vertex, sampled at 1,000 Hz, and filtered online between 0.1 and 100 Hz. Electrode impedances were maintained below 10 kΩ.

#### Continuous preprocessing

Offline preprocessing was performed in MATLAB R2023b (The MathWorks, Inc.) using EEGLAB (Delorme & Makeig, 2004). Line noise was first removed from the continuous recordings using Zapline-plus (De Cheveigné, 2020; Klug & Kloosterman, 2022) and CleanLine (Mullen, 2012). The data were then high-pass filtered at 0.25 Hz and subsequently low-pass filtered at 70 Hz using separate zero-phase finite impulse response filters. The clean_rawdata plugin (Kothe et al., 2019) identified channels with flat signals lasting longer than 5 s, line-noise contamination exceeding 4 *SD*s, or correlations with neighboring channels below .80. After these channels were removed, the data were downsampled to 250 Hz.

#### Independent component analysis (ICA)

ICA was estimated on a copy of the continuous data obtained after the preceding preprocessing steps, with an additional 1-Hz high-pass filter. The resulting weights were then applied to the corresponding preprocessed data without this additional filter. Components were classified using ICLabel (Pion-Tonachini et al., 2019), and those assigned a probability greater than .90 of belonging to the eye, muscle, or channel-noise categories were removed.

#### Epoching and artifact rejection

Following component removal, the data were segmented relative to T1 onset. Epochs spanning −200 to 800 ms were extracted for time-domain analyses and baseline-corrected using −200 to 0 ms. Longer epochs spanning −2,000 to 3,000 ms were extracted for time–frequency analyses to reduce edge effects during decomposition. The same rejection interval was used for both datasets: trials were excluded if voltage exceeded ±75 μV at any retained EEG channel during −200 to 800 ms. Previously removed channels were then reconstructed using spherical interpolation, and the data were re-referenced to the average of the scalp EEG channels. Subsequent analyses used 60 scalp channels.

#### EEG quality criteria and retained trials

Participant-level EEG quality criteria required an artifact-rejection rate no greater than 40% and no more than 10 bad channels. Among the 29 retained participants, the number of bad channels was *M* = 2.66 (*SD* = 1.72, range = 0–8), and the overall artifact-rejection rate was *M* = 4.89% (*SD* = 6.21%, range = 0–24.50%). DI_D and DI_DT analyses included the 16 and 18 participants, respectively, who had at least 15 artifact-free trials in the corresponding condition.

In the full sample, the numbers of artifact-free trials were *M* (*SD*) = 87.55 (8.02) for correct T, 336.31 (83.61) for correct TT, 203.52 (82.33) for correct TD, 106.52 (70.46) for DI_TD, 26.00 (23.01) for DI_D, and 25.03 (30.42) for DI_DT. In the DI_D subset, the corresponding correct TD and DI_D counts were 191.31 (79.62) and 40.31 (22.10). In the DI_DT subset, the correct TD and DI_DT counts were 156.83 (65.84) and 37.89 (32.50), respectively.

### ERP Analysis

#### ERP amplitude measurements

Artifact-free epochs were averaged separately for each participant and condition. ERP amplitudes and N2pc onset latency were quantified using ERPLAB (Lopez-Calderon & Luck, 2014). N2pc, Pd, and CDA were measured from contralateral-minus-ipsilateral difference waveforms at PO7/PO8, with laterality defined relative to T1 location. P3b and LPC were measured from waveforms averaged across Pz, P1, and P2. Mean-amplitude windows, selected from grand-average waveforms and previous research (Kappenman et al., 2021; Meng et al., 2023), were 200–260 ms for N2pc, 300–400 ms for Pd, 300–450 ms for P3b, and 450–700 ms for LPC and CDA. No additional filtering was applied for amplitude measurement.

#### N2pc onset latency

Onset was estimated using a jackknife procedure, with equally weighted leave-one-participant-out grand averages constructed separately for each comparison sample (Miller et al., 1998). Before measurement, the full N2pc waveforms were low-pass filtered at 10 Hz using ERPLAB’s second-order Butterworth filter. After prestimulus baseline correction, positive values were set to zero for area calculation. Onset was defined as the time at which cumulative negative area reached 30% of the total negative area within 180–300 ms. Tenfold temporal interpolation provided a 0.4-ms calculation grid from the original 4-ms sampling intervals. To correct for the reduction in variance introduced by jackknifing, paired *t* values were divided by *N* − 1, repeated-measures *F* values by (*N* − 1)², and standard errors were multiplied by *N* − 1, where *N* was the comparison sample size. These corrected quantities were used to calculate *p* values, confidence intervals (CIs), and jackknife-corrected standard errors (*SE*s) for the latency plots.

#### Dynamic time warping (DTW)

DTW assessed relative waveform timing between correct TD and each intrusion outcome (Hosseini et al., 2024). Analyses used PO7/PO8 N2pc waveforms within 150–350 ms and Pz/P1/P2 P3b waveforms within 250–550 ms. For DTW only, P3b waveforms were low-pass filtered at 25 Hz using a fourth-order Butterworth filter applied forward and backward before window selection; N2pc received no additional filtering. Participant waveforms were averaged with equal weights and *z*-standardized within each window. MATLAB’s *dtw* function aligned the intrusion waveform to its correct-TD reference. The normalized signed area between the alignment path and the identity diagonal quantified relative timing for inference. The median displacement between paired time indices was converted to milliseconds to describe delay, with positive values indicating later intrusion activity. Significance was assessed using 20,000 within-participant condition-label permutations, repeating grand averaging, standardization, and alignment for each permutation. Two-sided *p* values tested signed area, without Holm adjustment across components.

#### N2pc–P3b temporal coupling

Coupling was assessed separately for correct TD and each intrusion condition within the corresponding comparison sample. We generated 10,000 participant bootstrap resamples, using the same participant indices for both components. Each resampled grand average was z-standardized and aligned to the original grand average for that condition and component. Pearson and Spearman correlations between the paired signed-area estimates quantified covariation in N2pc and P3b timing. Between-condition differences were defined as the correct-TD correlation minus the intrusion correlation. Correlations and their differences were evaluated using 95% percentile CIs from a nested bootstrap with 2,000 outer participant resamples and 5,000 inner resamples per outer sample. Joint resampling across components and conditions preserved the paired structure.

### Time-Resolved Multivariate EEG Analyses

#### Data preparation

Multivariate analyses used artifact-free epochs spanning −200 to 800 ms from all 60 scalp electrodes. Voltages were summarized with 20-ms moving-average windows evaluated every 10 ms, yielding a common 100-Hz analysis grid. Preparation for pairwise decoding and the T–TT reference axis additionally included anti-alias filtering during resampling. Folds and repetitions were averaged to obtain one estimate per participant for each analysis.

#### Time-resolved decoding and temporal generalization

Pairwise decoding tested whether distributed EEG patterns distinguished conditions. We used shrinkage-regularized linear discriminant analysis (LDA), implemented with the custom MATLAB function *fit_lda*, with equal class priors and oracle-approximating shrinkage (OAS) estimation of within-class covariance. Features were centered and scaled using training data only. Stratified fivefold cross-validation was repeated 20 times with new random partitions. In each repetition, four folds trained the classifier and the remaining fold tested it, rotating through all five folds. Within each training fold, the larger class was randomly subsampled to match the smaller class; test trials retained their original class counts. Performance was quantified as the area under the receiver operating characteristic curve (AUC; chance = .50) and averaged across the 100 held-out evaluations. Temporal generalization applied classifiers trained at one time point to held-out trials at every test time point (King & Dehaene, 2014).

#### Activation-pattern topographies

To describe the spatial distribution of decoding information, classifier weights were transformed into activation patterns using the training-data covariance matrix (Haufe et al., 2014). Patterns were averaged across folds and repetitions, *z*-standardized over electrodes within each participant and time point, and averaged over the displayed windows. The resulting topographies describe scalp activity associated with the classifier decision variable.

#### Cross-condition classification

Correct T and TT trials trained ensembles of 20 classifiers on independently balanced samples. The fitted T–TT classifiers were then applied to correct TD and intrusion trials, which never entered source training. Outputs included the proportion of trials classified as TT and transfer AUCs distinguishing an intrusion from correct TD along the trained direction. Source-validation controls used stratified fivefold cross-validation repeated five times, with 20 balanced classifiers per training fold. Correct T and TT were scored only when held out. Continuous scores were divided by the training-data pooled within-class *SD* of projected scores and compared with the midpoint of held-out T and TT scores. The DI_DT extension applied the saved source ensemble to independent DI_DT and TD trials. Classification proportions and projected scores were interpreted relative to the trained decision boundary.

#### Reference-axis and representational-distance analyses

Correct T and TT training trials alone defined a noise-normalized spatial direction. Shrinkage covariance was estimated from their within-condition residuals pooled over 0–800 ms. At each time point, the training TT-minus-T pattern defined the axis, and held-out patterns were projected relative to the held-out T origin. Intrusion-minus-TD coordinate differences therefore quantified changes along the same independently trained T–TT direction. Relative proximity was also estimated from crossvalidated Mahalanobis distances (crossnobis): each held-out partition was divided into two independent halves, and noise-normalized difference patterns were multiplied across halves and averaged over electrodes (Walther et al., 2016). Distance to T minus distance to TT was negative for relative T proximity. Fivefold cross-validation was repeated 20 times.

#### Representational similarity analysis (RSA)

RSA assessed the distances between DI_DT and correct T, TT, and TD activity. Noise metrics were estimated from independent correct T/TT trials; these trials were excluded from reference-pattern estimation. Within each of five noise-metric folds, the remaining trials were assigned to five pattern folds, with 20 repetitions. For each pattern fold, its intrusion-minus-reference pattern was multiplied by the independently estimated pattern from the other four folds, weighted by their trial counts. Negative crossnobis estimates were retained. Same-time distances were evaluated along the diagonal; cross-time analyses assessed reference-relative distances and generalization of condition-difference patterns. Absolute distance from zero and relative proximity to T versus TT were tested separately.

#### Pattern component modeling (PCM)

PCM compared candidate accounts of shared and distinct activity patterns using the pattern component modeling toolbox (Diedrichsen et al., 2011, 2018; Diedrichsen & Kriegeskorte, 2017). Each analysis included correct T, TT, TD, and one intrusion condition. Participant-level patterns were estimated from independent trial partitions and expressed as T-minus-TD, TT-minus-TD, and intrusion-minus-TD contrasts. Seven models comprised a no-signal null model; three models constraining the intrusion contrast to match T, TT, or TD; rank-one and rank-two covariance models; and a free covariance model. Spatial whitening used only training-set residuals from correct T/TT/TD trials. Condition-specific residual variances and partition trial counts determined observation noise. Models were fitted by maximum likelihood at each time point and evaluated by conditional predictive log likelihood on held-out partitions, using fivefold cross-validation repeated three times. Scores were summed over held-out folds and averaged over repetitions. Higher scores indicated better held-out prediction; differences between scores were not interpreted as Bayes factors.

### Time-Frequency Analysis

#### Spectral decomposition and baseline normalization

Artifact-free long epochs were decomposed using the adaptive additive superlet transform (Moca et al., 2021) in FieldTrip (Oostenveld et al., 2011). This approach combines wavelets of different cycle lengths to improve the joint temporal and frequency localization of transient activity. The base wavelet width was three cycles, with a Gaussian width of three. Analyses covered 1–40 Hz in 1-Hz steps and −500 to 1,000 ms, with superlet orders increasing linearly from 1 to approximately 6. The same settings were used for Fourier-based and single-trial analyses. For condition-level total power, trial power was first averaged on the linear scale and then converted to decibels relative to the mean −500 to −200 ms power at each electrode and frequency: 10 × log_10_(power/baseline power). Electrode averaging was performed after normalization.

#### Electrode regions of interest (ROIs)

To characterize regional differences, power was averaged within seven ROIs: left frontal (lFC: F1, F3, F5, F7, FC1, FC3, FC5), right frontal (rFC: F2, F4, F6, F8, FC2, FC4, FC6), left central (C1, C3, C5, CP3, CP5), right central (C2, C4, C6, CP4, CP6), centroparietal (CP1, CPz, CP2, P1, Pz, P2, P3, P4), left posterior (P5, P7, PO3, PO7, O1), and right posterior (P6, P8, PO4, PO8, O2).

#### Induced and evoked activity

To assess the contribution of phase-locked activity, evoked power, induced power, and intertrial phase coherence (ITPC) were examined. Evoked power was obtained by transforming each participant’s condition-average ERP. Induced power was obtained by subtracting that ERP from each trial before decomposition and then averaging trial power. Both measures were normalized to their respective −500 to −200 ms baselines. ITPC was calculated as the magnitude of the mean unit-length complex phase vector at each electrode, frequency, and time point. To account for unequal trial counts, the larger condition was randomly subsampled to match the smaller condition within each participant; estimates were averaged over 1,000 repetitions. ITPC magnitudes were then averaged across lFC and, separately, all scalp electrodes. ITPC was not normalized to baseline power.

#### Frontal lateralization

Lateralization was assessed using five homologous electrode pairs: F3/F4, F5/F6, F7/F8, FC3/FC4, and FC5/FC6. Power was averaged within each five-electrode hemisphere group. The condition-by-hemisphere contrast was calculated as the left-minus-right difference in the first condition minus that in the second condition.

### Single-Trial Analysis

#### Model framework

Binomial and multinomial generalized linear mixed-effects models (GLMMs) related trial outcomes to behavioral or neural predictors using a logit link. Binomial models were fitted in R (version 4.5.2; R Core Team, 2025) with the *glmer* function from lme4 (Bates et al., 2015), using maximum likelihood with the Laplace approximation. Target location contrasted right with left; numerical distance was |T1 − PTD|, centered at 4 when used in interaction models. Neural predictors were centered within participant and divided by the *SD* of centered values pooled across participants. For the separate ERP models, scaling was established from artifact-free data across all 29 participants before selecting comparison subsets. ERP coefficients therefore represented the effect of a one-*SD* increase in signed amplitude.

#### Behavioral models

Separate models tested target location and numerical distance for each of DI_D, DI_TD, and DI_DT using all TD trials. The relevant intrusion was coded 1 and every other TD response 0. Each model included participant random intercepts and predictor slopes, with their correlation fixed at zero. The outcome therefore represented intrusion occurrence among all TD trials.

#### ERP models

Single-trial N2pc, Pd, P3b, LPC, and CDA amplitudes were extracted using the same electrodes and windows as the averaged ERPs, without latency-specific filtering. Each component was entered into a separate random-intercept model without location or distance covariates. Models contrasted each intrusion with correct TD for DI_TD (*n* = 29), DI_DT (*n* = 18), and DI_D (*n* = 16), and correct TT with T (*n* = 29). Follow-up DI_D models tested component-by-distance interactions while including the main effects of component amplitude, numerical distance, and target location. Each interaction model was compared with its additive counterpart using identical observations and random-effects structure.

#### Time–frequency models

Trial-level total power was converted to decibels relative to each trial’s own baseline before feature extraction. lFC power was averaged across its seven electrodes and the samples comprising the complete significant lFC time–frequency mask. For lateralization, left and right power were extracted from the same samples in the condition-by-hemisphere mask. Models coded DI_TD as 1 and correct TD as 0. One model included lFC power alone. A second included left-minus-right power, bilateral mean power, and target location. An equivalent parameterization included left and right power plus location, using a common *SD* scale for both hemispheres. Nested model comparisons assessed the additional contribution of each hemisphere.

#### Distance effects on ERP amplitudes

Supplementary Gaussian linear mixed-effects models tested numerical-distance effects across all artifact-free TD trials, including correct reports, all intrusion types, and other errors. Separate models for each ERP component included within-participant-centered distance, participant mean distance, target location, and report category. Participant random intercepts and distance slopes were retained when estimable. If a fit was singular, intercept–slope correlations were removed first, followed by random slopes if singularity persisted. Final models were fitted by restricted maximum likelihood with lmerTest, using Satterthwaite degrees of freedom (Kuznetsova et al., 2017).

#### Joint ERP and time–frequency models

A multinomial GLMM jointly related ERP amplitudes and frontal power to correct TD, DI_TD, DI_D, and DI_DT reports, with correct TD as the reference. The main analysis included all 29 participants and 10,093 trials with valid values for every predictor. The model included N2pc, Pd, P3b, LPC, CDA, lFC power, rFC power, numerical distance, and target location. ERP electrodes and windows were unchanged. The lFC feature used its seven-electrode regional mask, whereas the rFC feature used right-sided power from the five-electrode group within the frontal-lateralization mask. These were total-power features and used different time–frequency masks. Each neural predictor was centered within participant across the four retained TD report categories and divided by the pooled centered *SD* in the full sample. Numerical distance was standardized across retained trials; location was coded right = 0.5 and left = −0.5. Participant mean neural values were not entered as predictors. Models were fitted with the *mblogit* function in the mclogit package (Elff, 2025) using penalized quasi-likelihood with a maximum-likelihood approximation and category-specific participant random intercepts whose between-category covariances were fixed at zero. A sensitivity analysis used the 11 participants who met both condition-specific trial-count criteria, retaining the full-sample centering and scaling.

#### Model inference

GLMM fixed effects were evaluated with two-sided Wald *z* tests and reported as *b*, *SE*s, and odds ratios (ORs). Likelihood-ratio tests evaluated interactions and nested GLMMs; convergence and singularity were checked. For behavioral models, Holm-adjusted *p* values and Bonferroni-adjusted OR CIs covered the three intrusion types separately for each predictor. ERP main effects were Holm-adjusted across five components within each comparison; DI_D distance interactions formed a separate five-component family. ERP and power GLMMs used unadjusted 95% Wald OR CIs, and full-cluster power models used unadjusted *p* values. Distance–ERP models used Holm correction across five components and unadjusted 95% CIs. Fixed-effect prediction curves set random effects to zero, whereas participant curves included fitted random effects. Displayed intervals were pointwise. For the joint multinomial models, Holm correction covered all 27 slope tests (nine predictors across three intrusion-versus-correct contrasts); 95% Wald CIs were unadjusted. Convergence, fixed-effect covariance matrices, and predictor collinearity were checked.

### Statistical Analysis

#### Parametric tests

Parametric analyses were performed in JASP (JASP Team, 2025). Repeated-measures analyses of variance (ANOVAs) tested intrusion-type differences, accuracy differences across T, TD, and TT, and ERP condition effects among correct T, TD, and TT trials. Greenhouse–Geisser correction was applied when Mauchly’s test indicated a violation of sphericity; otherwise, uncorrected degrees of freedom and *p* values were reported. Paired *t* tests compared conditions, and one-sample *t* tests evaluated participant-level distance slopes against zero. Effect sizes were expressed as ω^2^ for repeated-measures ANOVAs (Bakeman, 2005; Olejnik & Algina, 2003) and Cohen’s *d* for *t* tests. For amplitude comparisons, *BF*_01_ quantified evidence for the null relative to the alternative hypothesis where reported.

Holm correction was applied separately within the three intrusion-type comparisons, three accuracy comparisons, three intrusion-versus-other-error comparisons, three location tests, and three distance-slope tests. ERP amplitude tests were adjusted across five components within each TD–intrusion comparison; post hoc comparisons among correct T, TD, and TT were adjusted across three pairs within each component. Amplitude, location, and distance-slope CIs used Bonferroni correction within the corresponding families. Inference for single-trial models and DTW is described in the respective sections.

#### Cluster-based nonparametric permutation tests

ERP waveform, multivariate, and time– frequency effects were evaluated using cluster-based permutation tests in FieldTrip (Maris & Oostenveld, 2007; Oostenveld et al., 2011). Sample-level *t* statistics were thresholded at *p* < .05, and neighboring suprathreshold samples were grouped into clusters by summing their *t* values. Within-participant condition swaps or sign flips generated a null distribution of the maximum cluster statistic. Comparing observed clusters with this distribution controlled multiplicity across the samples searched, including time points, electrode samples, time–frequency samples, or cells of temporal-generalization matrices. Analyses used 20,000 randomizations. AUC was tested against .50 using one-sided tests, whereas condition differences used two-sided tests. Clusters with *p*_cluster_ < .05 were considered significant; .05 ≤ *p*_cluster_ < .10 was displayed as marginal where specified. Where correction across multiple ROIs or contrasts was applied, it used Holm adjustment or a joint maximum-cluster distribution, as specified for each analysis.

## Additional information

### Funding and acknowledgements

This work was supported by grants from the National Natural Science Foundation of China to Antao Chen (grant numbers: 32171040 and 32371105).

### Author contributions

Zong Meng: Conceptualization, Methodology, Investigation, Formal analysis, Writing – original draft, Writing – review & editing.

Qi Chen: Investigation, Formal analysis.

Liqin Zhou: Conceptualization, Writing – review & editing.

Ke Zhou: Conceptualization, Writing – review & editing.

Antao Chen: Conceptualization, Methodology, Funding acquisition, Writing – review & editing.

### Ethics

The study was approved by the Ethics Committee of the Faculty of Psychology at Southwest University (approval no. H22049) and was conducted in accordance with the Declaration of Helsinki. All participants provided written informed consent before participation.

### Data and code availability

De-identified data, analysis code, and experimental materials supporting the analyses reported in this manuscript will be made publicly available in a trusted repository upon publication. Repository details will be provided in the published article.

