## Supplementary Material for "Substitution and Concomitant Distractor Intrusions Reveal Different Neural Dynamics of Temporal Selection"

### Figure supplement index

Figure 1—figure supplements 1–2: participant-level and trial-level behavior.

Figure 2—figure supplements 1–8: ERP amplitudes, waveforms, timing, and single-trial analyses; figure supplements 9–10: additional ERP analyses near the end of this file.

Figure 3—figure supplements 1–7: decoding, temporal generalization, covariance patterns, and representational comparisons.

Figure 4—figure supplements 1–5: regional power, induced and evoked activity, phase coherence, and control comparisons.

Figure 5—figure supplement 1: joint neural associations in the intersection sample.

Supplementary statistical tables retain the labels Table S1–Table S21. This file combines figure supplements and editable tables for review.

### 1. Supplementary behavior results

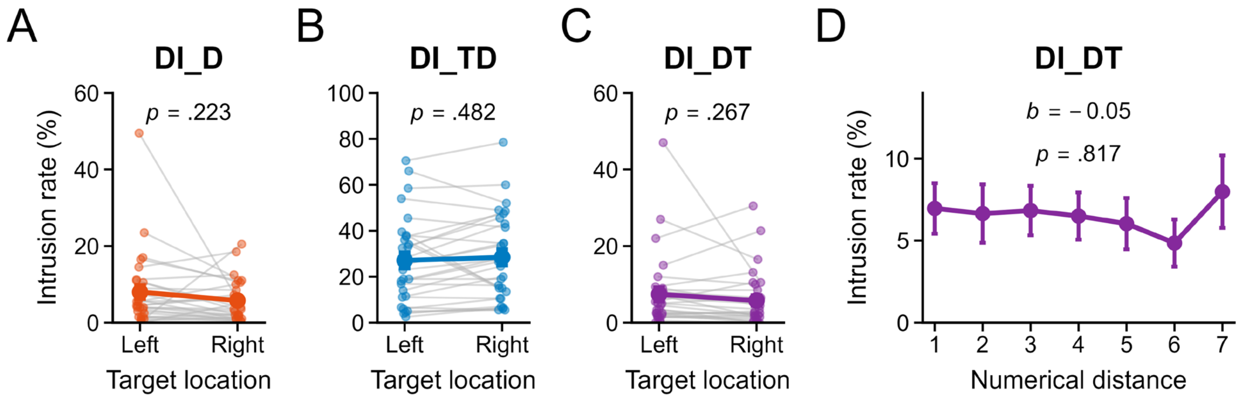


**Figure 1—figure supplement 1. Participant-level effects of target location and numerical distance on distractor intrusion.**

Distractor intrusion (DI) on target–distractor (TD) trials comprised DI_D (only the posttarget distractor [PTD] reported), DI_TD (target then PTD), and DI_DT (PTD then target). (**A–C**) Rates for left- and right-target trials. Faint points and gray lines show paired participant values; larger points and error bars show means (*M*) ± standard error of the mean (*SEM*). Left-minus-right comparisons were nonsignificant: DI_D (**A**), $t(28) = 1.25$, $p = .223$, $p_{\mathrm{Holm}} = .670$, 95% confidence interval (CI) [−2.31, 6.75], Cohen's $d = 0.23$; DI_TD (**B**), $t(28) = -0.71$, $p = .482$, $p_{\mathrm{Holm}} = .670$, 95% CI [−6.27, 3.53], $d = -0.13$; DI_DT (**C**), $t(28) = 1.13$, $p = .267$, $p_{\mathrm{Holm}} = .670$, 95% CI [−2.02, 5.25], $d = 0.21$. (**D**) DI_DT by absolute target–PTD numerical distance (*M* ± *SEM*): $b = -0.05$, standard error (*SE*) = 0.20, $t(28) = -0.23$, $p = .817$, $p_{\mathrm{Holm}} = .817$, 95% CI [−0.57, 0.47], $d = -0.03$. *b* is the percentage-point change per unit distance. Rates use TD trials in the corresponding category as the denominator ($n = 29$). Holm-adjusted *p* values and Bonferroni CIs cover three intrusion types separately for location and distance; plot annotations show raw *p* values.

**Table S1. Intrusion rates relative to other errors.**

| Intrusion minus other errors | Difference [95% CI] | $t(28)$ | $p$ | $p_{\text{Holm}}$ | $d$ | $BF_{01}$ |
|---|---|---|---|---|---|---|
| DI_TD | 23.91 [15.42, 32.40] | 7.17 | **< .001** | **< .001** | 1.33 | < 0.01 |
| DI_D | 3.03 [−0.20, 6.26] | 2.39 | **.024** | **.048** | 0.44 | 0.45 |
| DI_DT | 2.73 [−0.38, 5.84] | 2.23 | **.034** | **.048** | 0.41 | 0.60 |

***Note.*** $n = 29$. Other errors are incorrect TD reports outside DI_TD, DI_D, and DI_DT ($M = 3.86$, standard deviation [$SD$] = 3.88). Rates are percentages of all TD trials; differences are in percentage points. Tests are two-sided and paired. $p_{\text{Holm}}$ and Bonferroni CIs cover the three intrusion-versus-other-error comparisons. $d$ uses the $SD$ of paired differences; $BF_{01}$ is the Bayes factor favoring the null over the alternative hypothesis. Holm tests and Bonferroni CIs use different procedures, so a Holm-significant comparison may have a CI spanning zero. Significant raw or adjusted $p$ values are bold.

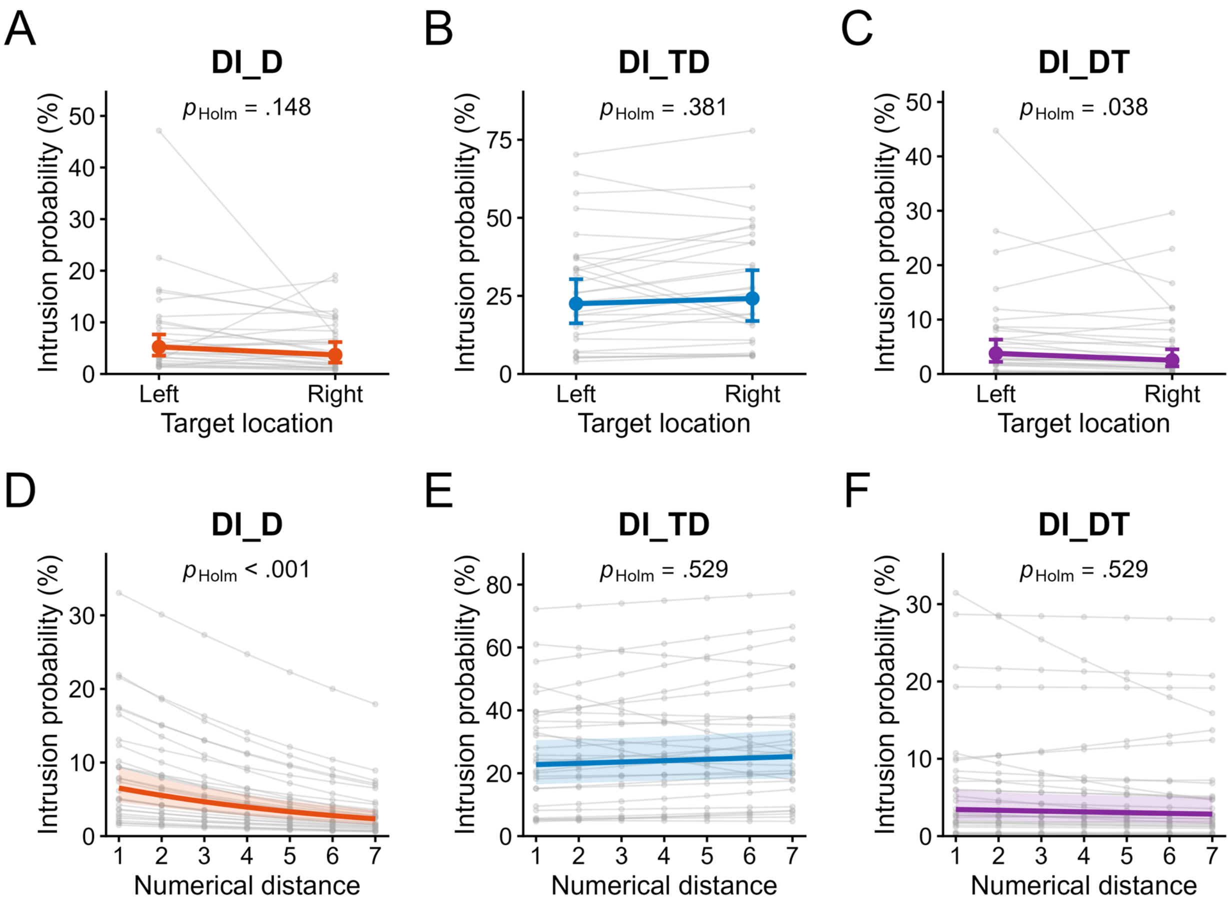


**Figure 1—figure supplement 2. Trial-level effects of target location and numerical distance on distractor intrusion.**

Binomial generalized linear mixed-effects models (GLMMs) assessed target location (**A–C**) and numerical distance (**D–F**) separately for DI_D (**A, D**), DI_TD (**B, E**), and DI_DT (**C, F**). Colored points and lines show fixed-effect predictions; pale lines show participant-specific predictions. Error bars and shading are unadjusted 95% CIs. Annotated $p_{Holm}$ values adjust the three intrusion types separately for each predictor ($n$ = 29; **Table S2**).

**Table S2. Trial-level effects of location and numerical distance on intrusion (Figure 1—figure supplement 2).**

| Intrusion | Predictor | *b* | *SE* | *z* | *p* | $p_{Holm}$ | *OR* | 95% CI for *OR* |
|---|---|---|---|---|---|---|---|---|
| DI_D | Right vs left | −0.36 | 0.20 | −1.79 | .074 | .148 | 0.70 | [0.43, 1.13] |
| DI_TD | Right vs left | 0.09 | 0.11 | 0.88 | .381 | .381 | 1.10 | [0.85, 1.42] |
| DI_DT | Right vs left | −0.41 | 0.17 | −2.49 | **.013** | **.038** | 0.66 | [0.44, 0.98] |
| DI_D | Numerical distance | −0.18 | 0.03 | −6.28 | **< .001** | **< .001** | 0.84 | [0.78, 0.90] |
| DI_TD | Numerical distance | 0.02 | 0.02 | 1.12 | .264 | .529 | 1.02 | [0.97, 1.08] |
| DI_DT | Numerical distance | −0.03 | 0.03 | −1.03 | .305 | .529 | 0.97 | [0.90, 1.04] |

***Note.*** Each row is a separate binomial GLMM ($n = 29$): intrusion = 1, other TD responses = 0. Models included participant random intercepts and predictor slopes, with zero correlation. *b* is the log-odds coefficient; OR is the odds ratio; *z* is the Wald statistic. Location contrasts right with left; distance is in digit units. Two-sided Wald *p* values are raw or Holm-adjusted across three intrusion types per predictor. OR CIs use Bonferroni correction across the same tests; plot CIs remain pointwise. Significant raw or adjusted *p* values are bold.

### 2. Supplementary ERP results

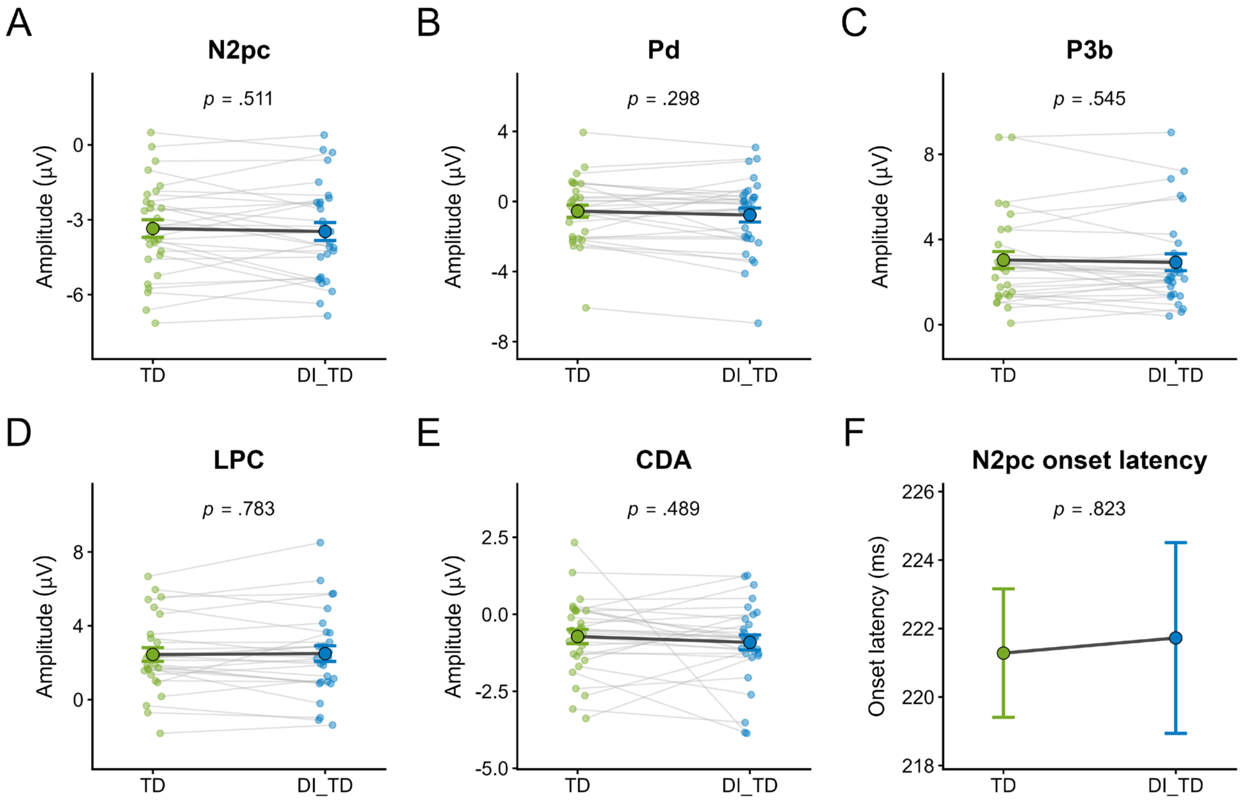


**Figure 2—figure supplement 1. Event-related potential measures for correct TD and DI_TD trials.**

Event-related potential (ERP) measures comprise N2 posterior contralateral (N2pc), distractor positivity (Pd), P3b, late positive complex (LPC), and contralateral delay activity (CDA) amplitudes (**A–E**), and N2pc onset latency (**F**; $n = 29$). N2pc, Pd, and CDA use contralateral-minus-ipsilateral PO7/PO8 waveforms; P3b and LPC use Pz/P1/P2 averages. Green denotes correct TD; blue denotes DI_TD. Faint points and gray lines show paired amplitudes; larger points and error bars show *M* ± *SEM*. Latency points show *M* leave-one-participant-out estimates with jackknife-corrected *SEM*. Annotations give raw two-sided paired-test *p* values; latency tests are jackknife-corrected (**Table S3**).

**Table S3. ERP comparisons between TD and DI_TD (Figure 2—figure supplement 1).**

| Measure | Correct TD | DI_TD | $t$ | $p$ | $p_{Holm}$ | 95% CI | $d$ | $BF_{01}$ |
|---|---|---|---|---|---|---|---|---|
| N2pc | −3.35 (1.89) | −3.47 (1.93) | 0.67 | .511 | 1.000 | [−0.37, 0.61] | 0.12 | 4.13 |
| Pd | −0.55 (1.87) | −0.78 (2.15) | 1.06 | .298 | 1.000 | [−0.36, 0.80] | 0.20 | 3.04 |
| P3b | 3.03 (2.15) | 2.93 (2.14) | 0.61 | .545 | 1.000 | [−0.37, 0.57] | 0.11 | 4.26 |
| LPC | 2.44 (2.00) | 2.49 (2.29) | −0.28 | .783 | 1.000 | [−0.55, 0.45] | −0.05 | 4.89 |
| CDA | −0.72 (1.25) | −0.91 (1.31) | 0.70 | .489 | 1.000 | [−0.56, 0.94] | 0.13 | 4.04 |
| N2pc onset latency | 221.28 | 221.72 | −0.23 | .823 | — | [−4.44, 3.56] | −0.04 | — |

***Note.*** $n = 29$. Amplitudes are $M$ (standard deviation [$SD$]), in µV; onset latencies are $M$ leave-one-participant-out estimates, in ms. Differences are TD minus DI_TD. Raw $p$ values are two-sided paired tests. $p_{Holm}$ and Bonferroni CIs cover the five amplitudes; latency $p$ values and CIs are unadjusted for multiplicity. Latency $t$, CI, and $d$ are jackknife-corrected. $d$ uses the $SD$ of paired differences. $BF_{01}$ is the Bayes factor for the null relative to the alternative, reported for amplitudes only. Significant $p$ values are bold.

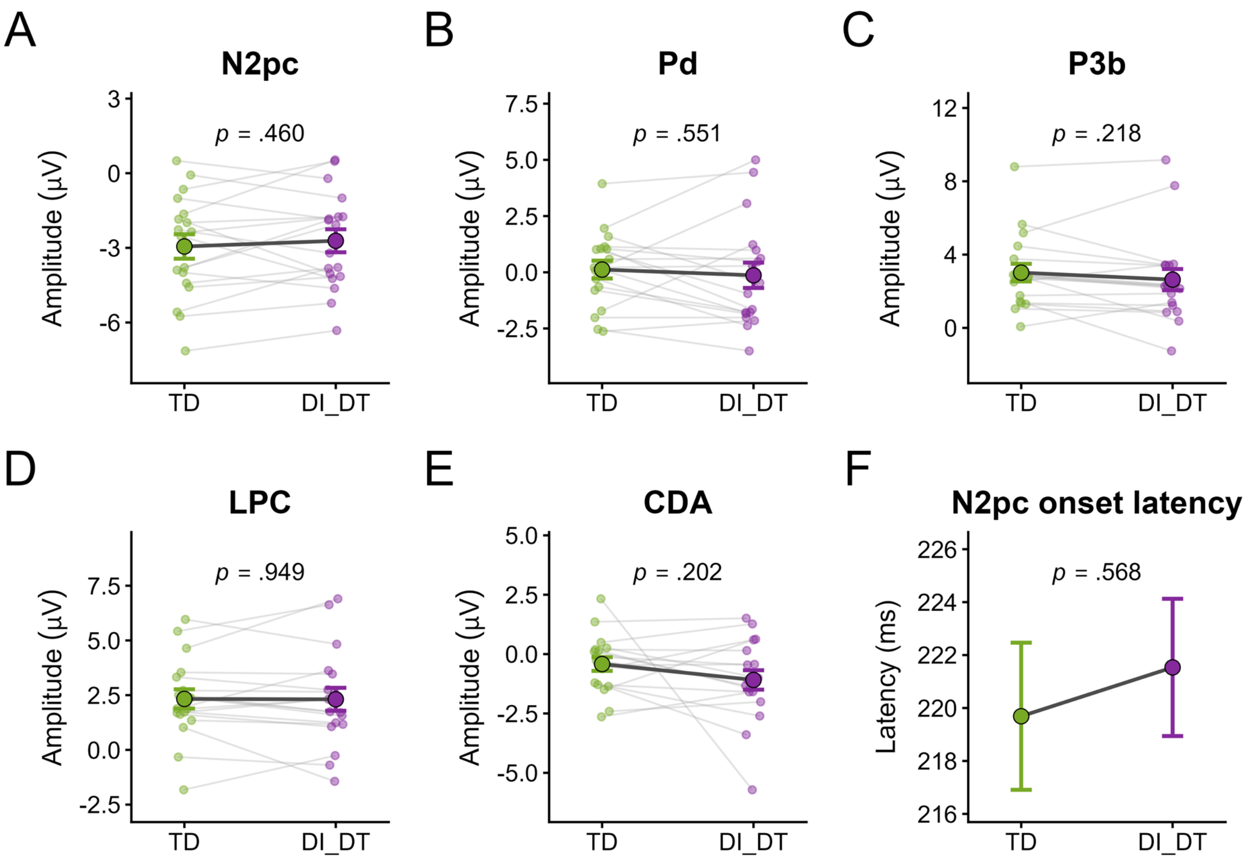


**Figure 2—figure supplement 2. ERP measures for correct TD and DI_DT trials.**

Correct TD (green) and DI_DT (purple), $n$ = 18. Measures and plotting conventions follow **Figure 2—figure supplement 1**; statistics are in **Table S4**.

**Table S4. ERP comparisons between TD and DI_DT (Figure 2—figure supplements 2–3).**

| Measure | Correct TD | DI_DT | $t$ | $p$ | $p_{Holm}$ | 95% CI | $d$ | $BF_{01}$ |
|---|---|---|---|---|---|---|---|---|
| N2pc | −2.95 (2.09) | −2.72 (1.96) | −0.76 | .460 | 1.000 | [−1.11, 0.65] | −0.18 | 3.19 |
| Pd | 0.12 (1.68) | −0.13 (2.39) | 0.61 | .551 | 1.000 | [−0.96, 1.47] | 0.14 | 3.49 |
| P3b | 3.02 (2.04) | 2.63 (2.46) | 1.28 | .218 | 1.000 | [−0.48, 1.25] | 0.30 | 2.04 |
| LPC | 2.33 (1.87) | 2.31 (2.23) | 0.06 | .949 | 1.000 | [−0.77, 0.81] | 0.02 | 4.11 |
| CDA | −0.42 (1.22) | −1.09 (1.73) | 1.33 | .202 | 1.000 | [−0.79, 2.13] | 0.31 | 1.94 |
| N2pc onset latency | 219.69 | 221.53 | −0.58 | .568 | — | [−8.52, 4.83] | −0.14 | — |

***Note.*** $n$ = 18; TD minus DI_DT. Reporting conventions follow **Table S3**.

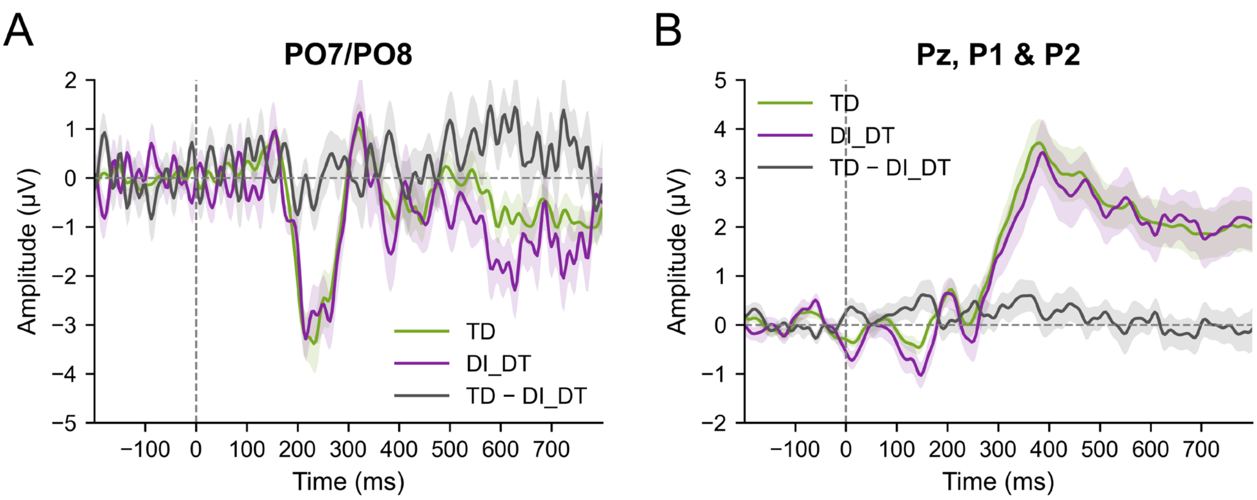


**Figure 2—figure supplement 3. ERP waveforms for correct TD and DI_DT trials.**

Grand-average ERPs for correct TD and DI_DT (*n* = 18; ≥10 valid DI_DT trials). **A**: contralateral-minus-ipsilateral PO7/PO8 waveforms relative to the first target (T1). **B**: Pz/P1/P2 averages. Green denotes correct TD, purple DI_DT, and gray TD minus DI_DT; shading shows ± *SEM*, including for paired differences. Dashed lines mark T1 onset and zero amplitude. Neither panel contained a significant cluster (two-sided permutation tests, −200–800 ms, 20,000 permutations). Component statistics are in **Table S4**.

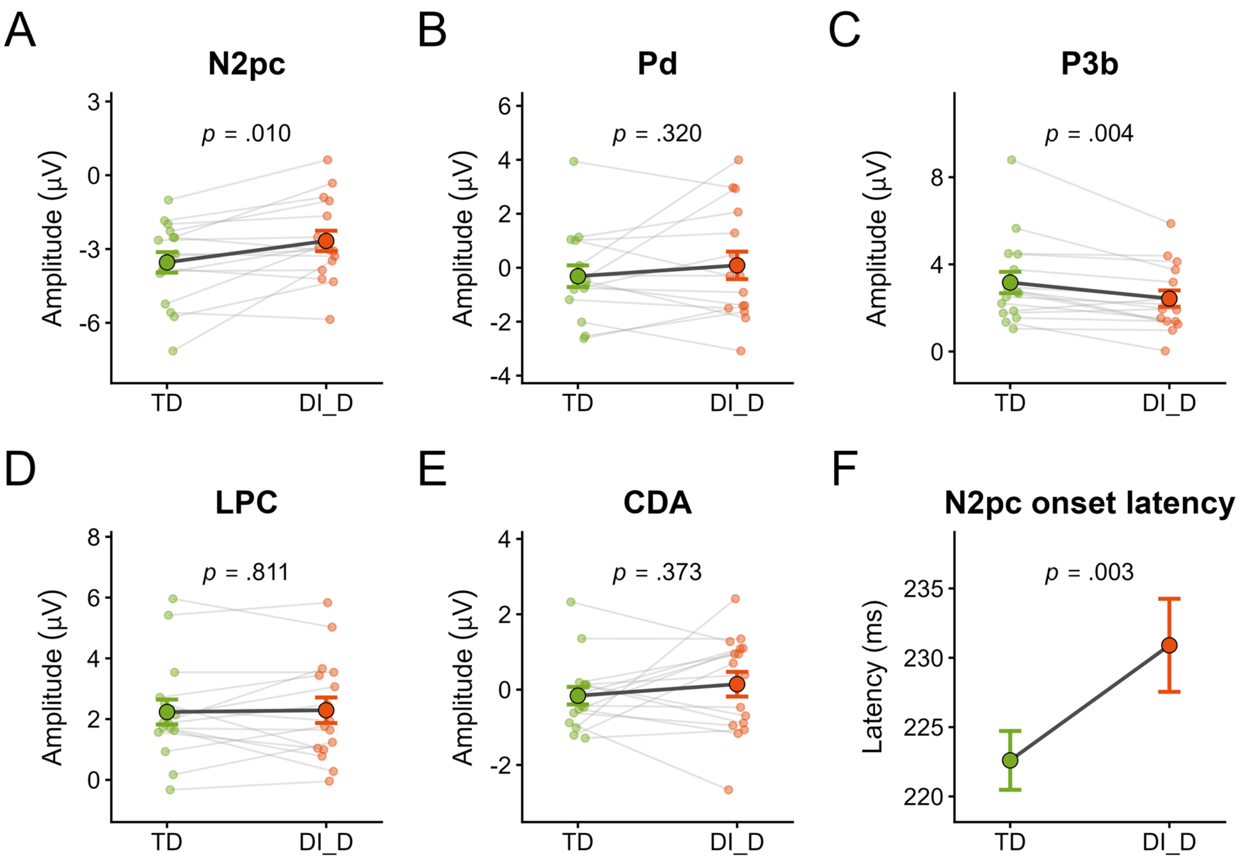


**Figure 2—figure supplement 4. ERP measures for correct TD and DI_D trials.**

Correct TD (green) and DI_D (orange), $n = 16$. Measures and plotting conventions follow **Figure 2—figure supplement 1**; statistics are in **Table S5**.

**Table S5. ERP comparisons between TD and DI_D (Figure 2—figure supplement 4).**

| Measure | Correct TD | DI_D | $t$ | $p$ | $p_{\text{Holm}}$ | 95% CI | $d$ | $BF_{01}$ |
|---|---|---|---|---|---|---|---|---|
| N2pc | −3.54 (1.67) | −2.67 (1.66) | −2.95 | **.010** | **.039** | [−1.74, 0.00]† | −0.74 | 0.18 |
| Pd | −0.32 (1.60) | 0.09 (2.04) | −1.03 | .320 | .959 | [−1.55, 0.75] | −0.26 | 2.48 |
| P3b | 3.17 (1.96) | 2.43 (1.50) | 3.35 | **.004** | **.022** | [0.09, 1.39] | 0.84 | 0.09 |
| LPC | 2.23 (1.64) | 2.29 (1.68) | −0.24 | .811 | .959 | [−0.75, 0.63] | −0.06 | 3.81 |
| CDA | −0.16 (0.93) | 0.14 (1.31) | −0.92 | .373 | .959 | [−1.28, 0.67] | −0.23 | 2.72 |
| N2pc onset latency | 222.60 | 230.90 | −3.52 | **.003** | — | [−13.32, −3.28] | −0.88 | — |

***Note.*** $n$ = 16; TD minus DI_D. Reporting conventions follow **Table S3**. † The N2pc CI upper limit is below zero before rounding.

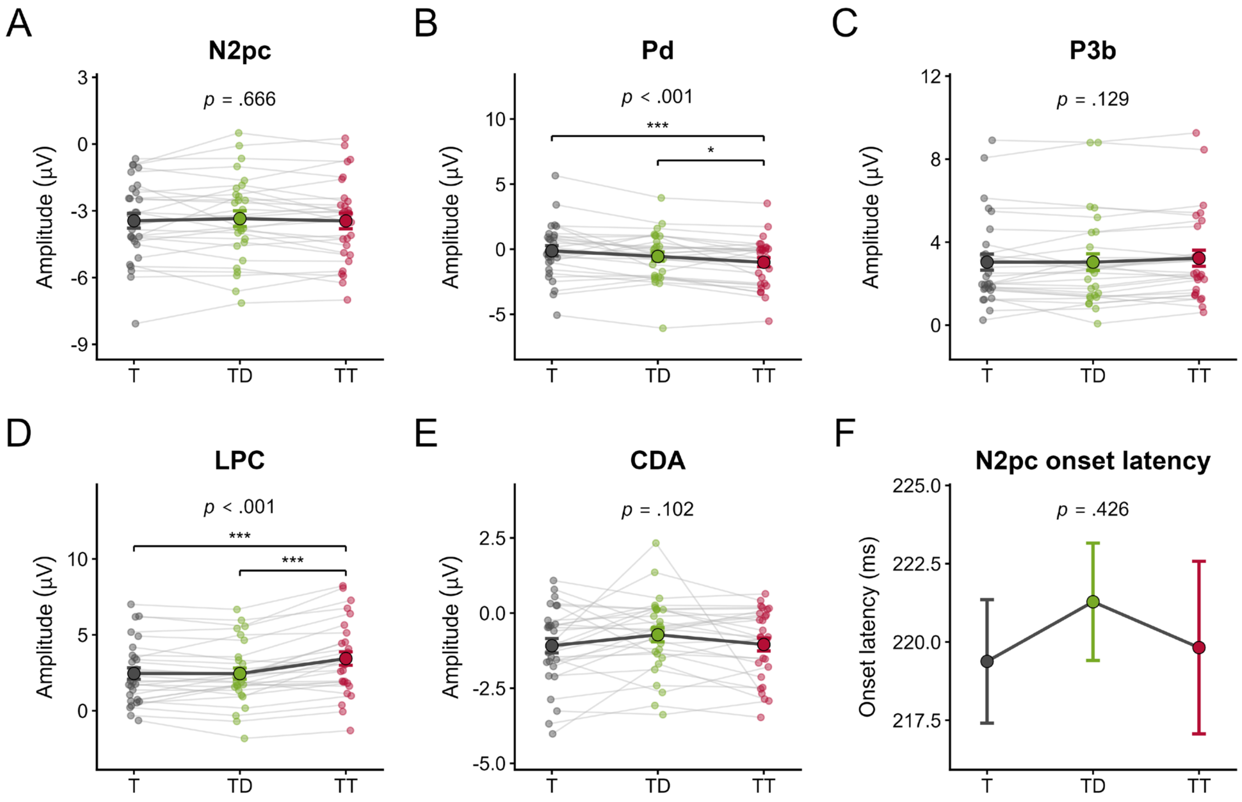


**Figure 2—figure supplement 5. ERP amplitudes and N2pc onset latency across correct-report conditions.**

Correct single-target (T), TD, and dual-target (TT) trials, $n = 29$. Panels show N2pc (A), Pd (B), P3b (C), LPC (D), CDA (E), and N2pc onset latency (*F*). Dark gray denotes T, green TD, and red TT; measurement and plotting follow **Figure 2—figure supplement 1**. Annotations give repeated-measures analysis of variance (ANOVA) *p* values: Greenhouse–Geisser (GG) correction was applied to Pd, P3b, and LPC after significant Mauchly tests; N2pc, CDA, and onset latency use uncorrected degrees of freedom. Latency statistics are jackknife-corrected. Brackets mark amplitude comparisons with Holm correction across three pairs per component (* $p_{\text{Holm}} < .05$, ** < .01, *** < .001). Latency comparisons are unadjusted for multiplicity (**Table S6**).

**Table S6. ERP measures across correct T, TD, and TT trials (Figure 2—figure supplement 5).**

**A. Descriptive statistics**

| Measure | T | TD | TT |
|---|---|---|---|
| N2pc | −3.45 (1.78) | −3.35 (1.89) | −3.46 (1.88) |
| Pd | −0.12 (2.12) | −0.55 (1.87) | −1.01 (1.87) |
| P3b | 3.03 (2.09) | 3.03 (2.15) | 3.22 (2.09) |
| LPC | 2.45 (2.03) | 2.44 (2.00) | 3.44 (2.39) |
| CDA | −1.09 (1.29) | −0.72 (1.25) | −1.05 (1.16) |
| N2pc onset latency | 219.38 (1.97) | 221.28 (1.88) | 219.82 (2.76) |

**B. Omnibus condition effects**

| Measure | $F$ | $df_1$ | $df_2$ | $\varepsilon_{GG}$ | $p$ | $\omega^2$ |
|---|---|---|---|---|---|---|
| N2pc | 0.41 | 2 | 56 | — | .666 | 0.00 |
| Pd | 12.47 | 1.65 | 46.18 | 0.82 | **< .001** | 0.03 |
| P3b | 2.23 | 1.59 | 44.66 | 0.80 | .129 | 0.00 |
| LPC | 22.76 | 1.50 | 41.90 | 0.75 | **< .001** | 0.04 |
| CDA | 2.38 | 2 | 56 | — | .102 | 0.01 |
| N2pc onset latency | 0.87 | 2 | 56 | — | .426 | 0.00 |

***Note.*** $n = 29$. A: amplitudes are *M* (*SD*), in µV; onset latencies are *M* (jackknife-corrected *SEM*), in ms. B: GG-corrected *df* and *p* values are reported for Pd, P3b, and LPC; $\varepsilon_{GG}$ is the correction factor. N2pc, CDA, and onset latency use uncorrected *df* and *p* values because their Mauchly tests were nonsignificant. Latency statistics remain jackknife-corrected. $\omega^2$ is omega squared. Significant *p* values are bold.

**C. Post hoc comparisons**

| Measure | Comparison | Difference [95% CI] | $t$ | $p$ | $p_{\text{Holm}}$ | $d$ | $BF_{01}$ |
|---|---|---|---|---|---|---|---|
| N2pc | T − TD | −0.10 [−0.47, 0.28] | −0.66 | .512 | 1.000 | −0.05 | 4.14 |
| N2pc | T − TT | 0.01 [−0.33, 0.34] | 0.05 | .959 | 1.000 | 0.00 | 5.06 |
| N2pc | TD − TT | 0.10 [−0.16, 0.37] | 1.00 | .326 | .977 | 0.06 | 3.21 |
| Pd | T − TD | 0.43 [−0.11, 0.97] | 2.02 | .053 | .053 | 0.22 | 0.86 |
| Pd | T − TT | 0.88 [0.50, 1.26] | 5.90 | **< .001** | **< .001** | 0.45 | < 0.01 |
| Pd | TD − TT | 0.45 [0.04, 0.86] | 2.81 | **.009** | **.018** | 0.23 | 0.20 |
| P3b | T − TD | 0.00 [−0.27, 0.27] | 0.00 | 1.000 | 1.000 | 0.00 | 5.07 |
| P3b | T − TT | −0.19 [−0.51, 0.13] | −1.53 | .136 | .273 | −0.09 | 1.78 |
| P3b | TD − TT | −0.19 [−0.39, 0.01] | −2.47 | **.020** | .060 | −0.09 | 0.39 |
| LPC | T − TD | 0.01 [−0.32, 0.33] | 0.06 | .953 | .953 | 0.00 | 5.06 |
| LPC | T − TT | −0.98 [−1.52, −0.45] | −4.68 | **< .001** | **< .001** | −0.46 | < 0.01 |
| LPC | TD − TT | −0.99 [−1.40, −0.59] | −6.23 | **< .001** | **< .001** | −0.46 | < 0.01 |
| CDA | T − TD | −0.37 [−0.89, 0.16] | −1.76 | .089 | .267 | −0.30 | 1.29 |
| CDA | T − TT | −0.04 [−0.40, 0.32] | −0.30 | .770 | .770 | −0.03 | 4.87 |
| CDA | TD − TT | 0.32 [−0.17, 0.82] | 1.66 | .108 | .267 | 0.26 | 1.50 |
| N2pc onset latency | T − TD | −1.90 [−4.36, 0.55] | −1.59 | .124 | — | −0.16 | — |
| N2pc onset latency | T − TT | −0.44 [−3.73, 2.85] | −0.27 | .785 | — | −0.04 | — |
| N2pc onset latency | TD − TT | 1.46 [−2.01, 4.93] | 0.86 | .395 | — | 0.12 | — |

***Note.*** Differences are first minus second ($df = 28$), in µV for amplitudes and ms for latency. Amplitude $p_{\text{Holm}}$ and Bonferroni CIs cover three pairs per component; latency $p$ values and 95% CIs are unadjusted for multiplicity. Latency $t$ and CIs are jackknife-corrected. $d$ follows JASP's ANOVA post hoc standardizer; $BF_{01}$ is reported for amplitudes only. Significant $p$ values are bold.

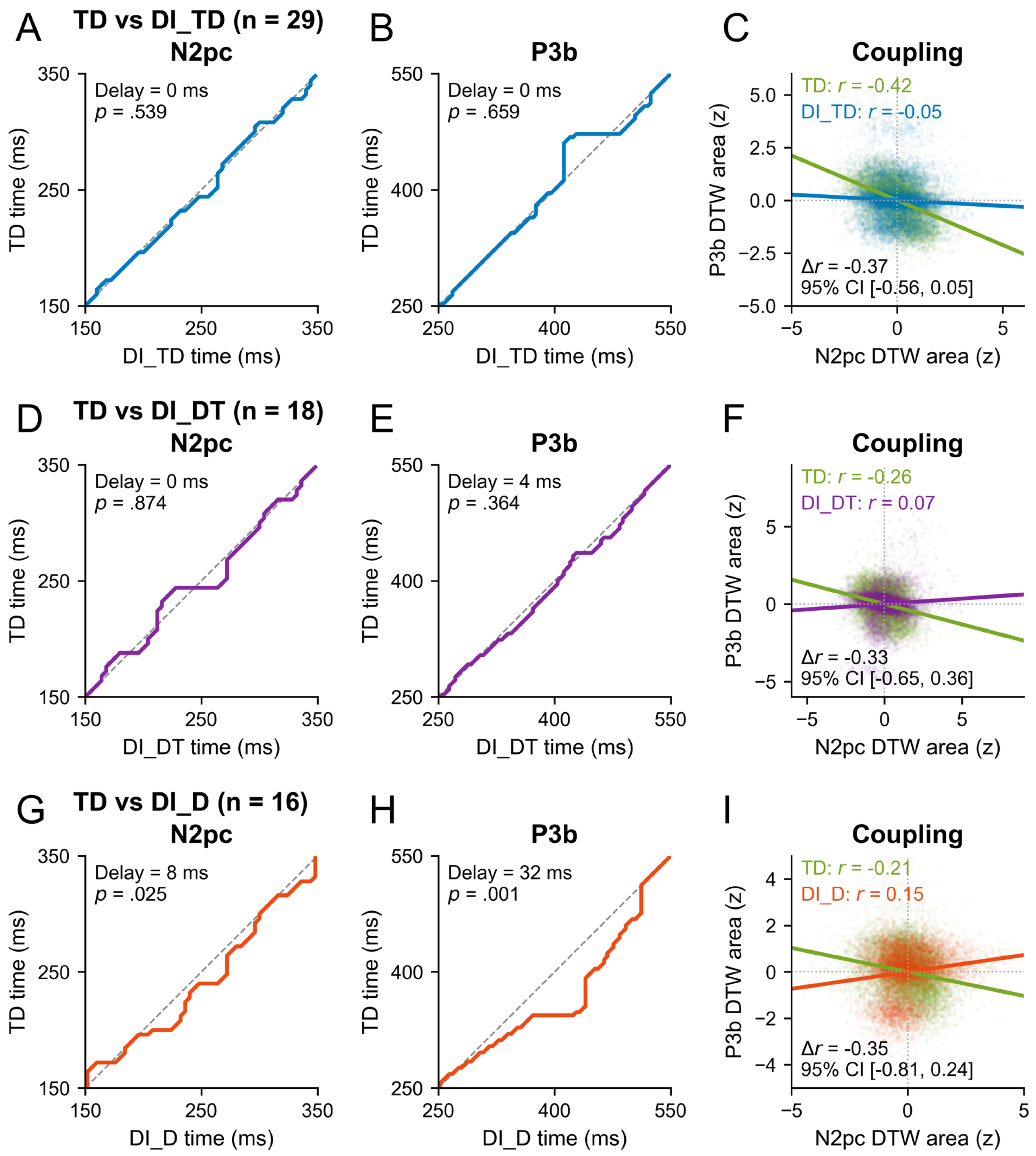


**Figure 2—figure supplement 6. N2pc and P3b timing and their coupling across intrusion outcomes.**

Dynamic time warping (DTW) compared correct TD with DI_TD (**A–C**; $n = 29$), DI_DT (**D–F**; $n = 18$), and DI_D (**G–I**; $n = 16$). Columns 1–2 show N2pc (150–350 ms) and P3b (250–550 ms) alignment paths; dashed diagonals indicate equal timing, and positive delays indicate later intrusion activity. Raw $p$ values test signed DTW area. Column 3 overlays correct TD (green) and intrusion (blue, purple, or orange) estimates. Each point is one participant-bootstrap grand average; axes show standardized signed DTW areas. Lines and $r$ values summarize Pearson coupling. **Δ*r*** is TD minus intrusion, with paired nested-bootstrap 95% CIs (**Table S7**).

**Table S7. DTW timing and N2pc–P3b coupling (Figure 2—figure supplement 6).**

**A. DTW timing comparisons.**

| Comparison | *n* | Measure | Window (ms) | $\Delta t$ (ms) | Signed area | *p* |
|---|---|---|---|---|---|---|
| TD vs DI_TD | 29 | N2pc | 150–350 | 0.00 | 0.01 | .539 |
| TD vs DI_TD | 29 | P3b | 250–550 | 0.00 | −0.02 | .659 |
| TD vs DI_DT | 18 | N2pc | 150–350 | 0.00 | 0.01 | .874 |
| TD vs DI_DT | 18 | P3b | 250–550 | 4.00 | 0.03 | .364 |
| TD vs DI_D | 16 | N2pc | 150–350 | 8.00 | 0.08 | **.025** |
| TD vs DI_D | 16 | P3b | 250–550 | 32.00 | 0.19 | **.001** |

***Note.*** $\Delta t$ is the descriptive median path displacement (intrusion minus TD); *p* values test normalized signed DTW area using 20,000 paired label permutations. ERPs were equally weighted and *z*-standardized. P3b was low-pass filtered at 25 Hz; N2pc received no additional filter. Minimum valid-trial counts were 15. Significant *p* values are bold.

**B. Within-condition N2pc–P3b coupling.**

| Comparison sample | *n* | Condition | Method | *r* | 95% CI | Coupling |
|---|---|---|---|---|---|---|
| TD–DI_TD | 29 | TD | Pearson | −0.42 | [−0.62, 0.04] | Not detected |
| | | | Spearman | −0.44 | [−0.65, 0.01] | Not detected |
| | | DI_TD | Pearson | −0.05 | [−0.36, 0.21] | Not detected |
| | | | Spearman | −0.08 | [−0.42, 0.20] | Not detected |
| TD–DI_DT | 18 | TD | Pearson | −0.26 | [−0.55, 0.27] | Not detected |
| | | | Spearman | −0.25 | [−0.56, 0.26] | Not detected |
| | | DI_DT | Pearson | 0.07 | [−0.38, 0.30] | Not detected |
| | | | Spearman | −0.15 | [−0.43, 0.29] | Not detected |
| TD–DI_D | 16 | TD | Pearson | −0.21 | [−0.54, 0.39] | Not detected |
| | | | Spearman | −0.23 | [−0.56, 0.40] | Not detected |
| | | DI_D | Pearson | 0.15 | [−0.30, 0.54] | Not detected |
| | | | Spearman | 0.11 | [−0.35, 0.55] | Not detected |

***Note.*** Each *r* is evaluated against zero by its 95% CI; “Not detected” means the CI includes zero. TD estimates use the corresponding comparison sample. Correlations use 10,000 participant-bootstrap grand averages; CIs use 2,000 outer resamples with 5,000 inner resamples. Components and conditions are jointly resampled. Bootstrap points are not independent observations for inference. No reliable coupling was detected for correct TD or any intrusion condition, and none of the three intrusion–TD coupling differences was reliable.

**C. Between-condition differences in N2pc–P3b coupling.**

| Comparison | *n* | Method | Δ*r* | 95% CI |
|---|---|---|---|---|
| TD–DI_TD | 29 | Pearson | −0.37 | [−0.56, 0.05] |
| TD–DI_TD | 29 | Spearman | −0.36 | [−0.59, 0.10] |
| TD–DI_DT | 18 | Pearson | −0.33 | [−0.65, 0.36] |
| TD–DI_DT | 18 | Spearman | −0.11 | [−0.62, 0.43] |
| TD–DI_D | 16 | Pearson | −0.35 | [−0.81, 0.24] |
| TD–DI_D | 16 | Spearman | −0.34 | [−0.84, 0.27] |

***Note.*** Δ*r* is TD minus intrusion; paired nested-bootstrap CIs follow **B**. All CIs include zero.

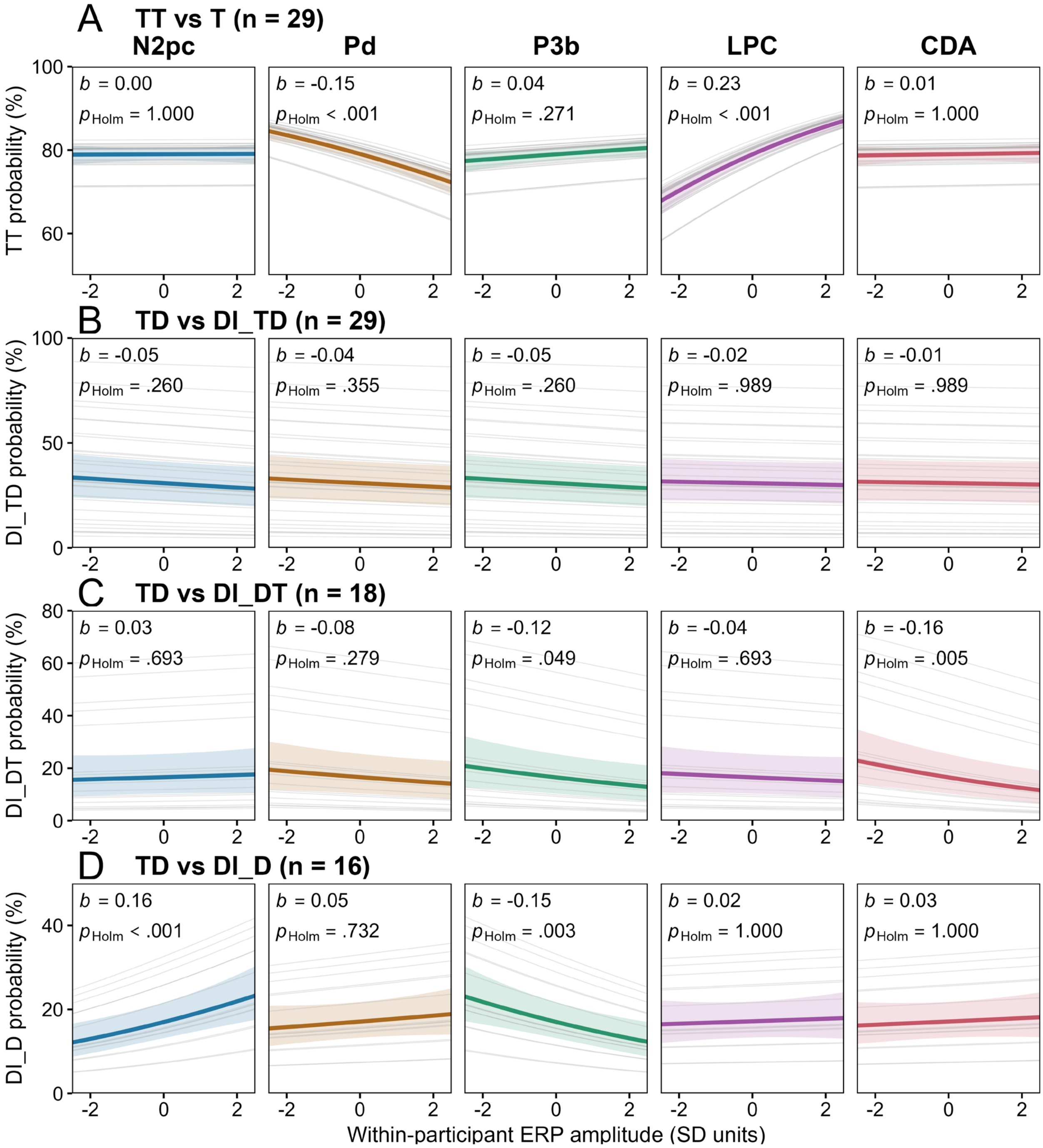


**Figure 2—figure supplement 7. Single-trial ERP associations with report outcomes.**

Binomial GLMMs compared TT with T (**A**; $n = 29$), and TD with DI_TD (**B**; $n = 29$), DI_DT (**C**; $n = 18$), or DI_D (**D**; $n = 16$); columns show N2pc, Pd, P3b, LPC, and CDA. Each model included one within-participant-centered amplitude, scaled by the pooled *SD*, and a participant random intercept; no location or distance covariates were included. Colored curves show fixed-effect predictions at random intercept zero, shading unadjusted pointwise 95% CIs, and gray lines participant-specific predictions. Annotations give $b$ and $p_{\mathrm{Holm}}$ across five ERP tests within each comparison (**Table S8**).

**Table S8. Single-trial ERP associations with report outcomes (Figure 2—figure supplement 7).**

**A. Correct TT versus T (*n* = 29).**

| Component | *b* | *SE* | *z* | *p* | $p_{Holm}$ | *OR* | 95% CI for *OR* |
|---|---|---|---|---|---|---|---|
| N2pc | 0.00 | 0.02 | 0.11 | .916 | 1.000 | 1.00 | [0.96, 1.05] |
| Pd | −0.15 | 0.02 | −6.65 | **< .001** | **< .001** | 0.86 | [0.82, 0.90] |
| P3b | 0.04 | 0.02 | 1.69 | .090 | .271 | 1.04 | [0.99, 1.09] |
| LPC | 0.23 | 0.02 | 10.02 | **< .001** | **< .001** | 1.26 | [1.20, 1.32] |
| CDA | 0.01 | 0.02 | 0.34 | .737 | 1.000 | 1.01 | [0.96, 1.05] |

**B. Correct TD versus DI_TD (*n* = 29).**

| Component | *b* | *SE* | *z* | *p* | $p_{Holm}$ | *OR* | 95% CI for *OR* |
|---|---|---|---|---|---|---|---|
| N2pc | −0.05 | 0.03 | −1.94 | .052 | .260 | 0.95 | [0.90, 1.00] |
| Pd | −0.04 | 0.03 | −1.56 | .118 | .355 | 0.96 | [0.91, 1.01] |
| P3b | −0.05 | 0.02 | −1.92 | .055 | .260 | 0.95 | [0.91, 1.00] |
| LPC | −0.02 | 0.02 | −0.68 | .494 | .989 | 0.98 | [0.94, 1.03] |
| CDA | −0.01 | 0.03 | −0.52 | .604 | .989 | 0.99 | [0.94, 1.04] |

**C. Correct TD versus DI_DT (*n* = 18).**

| Component | *b* | *SE* | *z* | *p* | $p_{Holm}$ | *OR* | 95% CI for *OR* |
|---|---|---|---|---|---|---|---|
| N2pc | 0.03 | 0.05 | 0.64 | .520 | .693 | 1.03 | [0.94, 1.13] |
| Pd | −0.08 | 0.05 | −1.68 | .093 | .279 | 0.93 | [0.85, 1.01] |
| P3b | −0.12 | 0.05 | −2.50 | **.012** | **.049** | 0.89 | [0.81, 0.98] |
| LPC | −0.04 | 0.05 | −0.94 | .347 | .693 | 0.96 | [0.87, 1.05] |
| CDA | −0.16 | 0.05 | −3.31 | **< .001** | **.005** | 0.85 | [0.77, 0.94] |

**D. Correct TD versus DI_D (*n* = 16).**

| Component | *b* | *SE* | *z* | *p* | $p_{Holm}$ | *OR* | 95% CI for *OR* |
|---|---|---|---|---|---|---|---|
| N2pc | 0.16 | 0.04 | 3.76 | **< .001** | **< .001** | 1.17 | [1.08, 1.27] |
| Pd | 0.05 | 0.04 | 1.16 | .244 | .732 | 1.05 | [0.97, 1.14] |
| P3b | −0.15 | 0.05 | −3.39 | **< .001** | **.003** | 0.86 | [0.79, 0.94] |
| LPC | 0.02 | 0.04 | 0.46 | .643 | 1.000 | 1.02 | [0.94, 1.11] |
| CDA | 0.03 | 0.04 | 0.66 | .507 | 1.000 | 1.03 | [0.95, 1.12] |

***Note.*** Separate binomial GLMMs included one ERP predictor and a participant random intercept, without location or distance covariates. **A**: TT = 1, T = 0; **B–D**: intrusion = 1, TD = 0. Amplitudes were centered within participant and divided by the pooled centered *SD* across all 29 participants. *b* and OR represent a one-*SD* increase in signed amplitude. Two-sided Wald $p_{Holm}$ values cover five components per comparison; OR CIs are unadjusted 95% Wald intervals. Minimum valid-trial counts were 15. Significant raw or adjusted *p* values are bold.

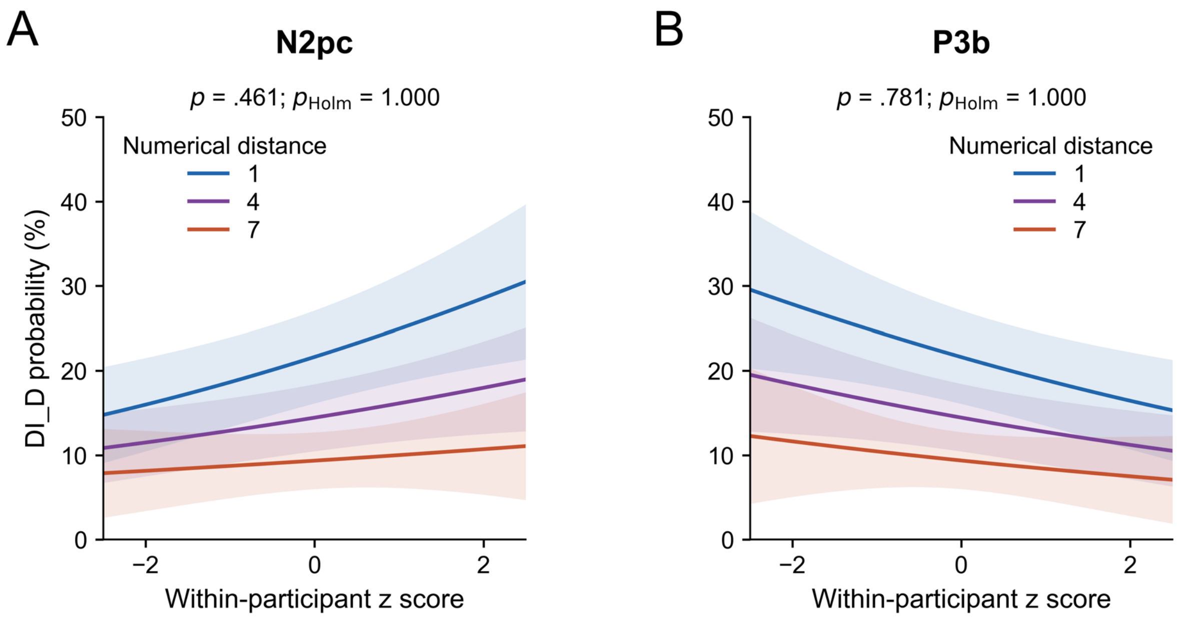


**Figure 2—figure supplement 8. Numerical-distance moderation of ERP associations with DI_D.**

Follow-up binomial GLMMs for DI_D versus correct TD ($n$ = 16) tested numerical-distance interactions with N2pc (**A**) and P3b (**B**). Curves show fixed-effect predictions at distances 1, 4, and 7, averaged equally over left and right target locations; shading shows unadjusted pointwise 95% CIs. The random intercept is set to zero. Annotations report likelihood-ratio $p$ values for the interaction and Holm-adjusted $p$ values across five ERP components. Differences between curve heights also reflect the main effect of distance and do not demonstrate an interaction. Full results for all five components are in **Table S9**.

**Table S9. Numerical-distance interactions with ERP predictors of DI_D (Figure 2—figure supplement 8).**

| Component | *b* | *SE* | $\chi^2(1)$ | *p* | $p_{Holm}$ | OR | 95% CI for OR |
|---|---|---|---|---|---|---|---|
| N2pc | −0.02 | 0.03 | 0.54 | .461 | 1.000 | 0.98 | [0.93, 1.03] |
| Pd | −0.01 | 0.02 | 0.10 | .758 | 1.000 | 0.99 | [0.95, 1.04] |
| P3b | 0.01 | 0.03 | 0.08 | .781 | 1.000 | 1.01 | [0.95, 1.07] |
| LPC | 0.03 | 0.03 | 0.90 | .342 | 1.000 | 1.03 | [0.97, 1.08] |
| CDA | −0.03 | 0.03 | 1.09 | .296 | 1.000 | 0.97 | [0.93, 1.02] |

***Note.*** $n = 16$; 3,706 trials (3,061 correct TD, 645 DI_D). Each binomial GLMM included one ERP amplitude, numerical distance, their interaction, target location, and a participant random intercept; DI_D = 1, TD = 0. ERP scaling follows **Table S8**; distance is |T1 − PTD|, centered at 4. *b* and OR describe the change in the ERP slope per unit increase in distance. *p* values are likelihood-ratio tests of the interaction (1 *df*); $p_{Holm}$ adjusts five components. OR CIs are unadjusted 95% Wald intervals. All models converged without singular fits. **Figure 2—figure supplement 8** displays N2pc and P3b.

### 3. Supplementary multivariate results

Additional multivariate analyses assess diagonal decoding, temporal generalization, classification covariance patterns, and DI_DT. Analyses use 29 participants; DI_D comparisons use 16 participants with ≥15 valid trials. DI_DT analyses use 18 participants with ≥15 valid trials; DI_DT–DI_D comparisons use the 11-participant intersection. Marginal clusters ($.05 \leq p_{\text{cluster}} < .10$) do not meet the .05 threshold. Saved temporal, matrix, or joint-model corrections are retained, without additional Holm correction.

Trial-pair sensitivity analysis. Decoding was repeated after averaging two trials within each class. Original trials were assigned to training and test partitions before averaging, and trial pairs were constructed separately within each partition. Training classes were balanced by random subsampling. Stratified threefold cross-validation was repeated 20 times, with new random partitions and pairings; performance was averaged over folds and repetitions. No original trial entered both training and test data in the same fold. Results are reported in **Table S10C** and, for DI_DT, in **Figure 3—figure supplement 7F and Table S13B**.

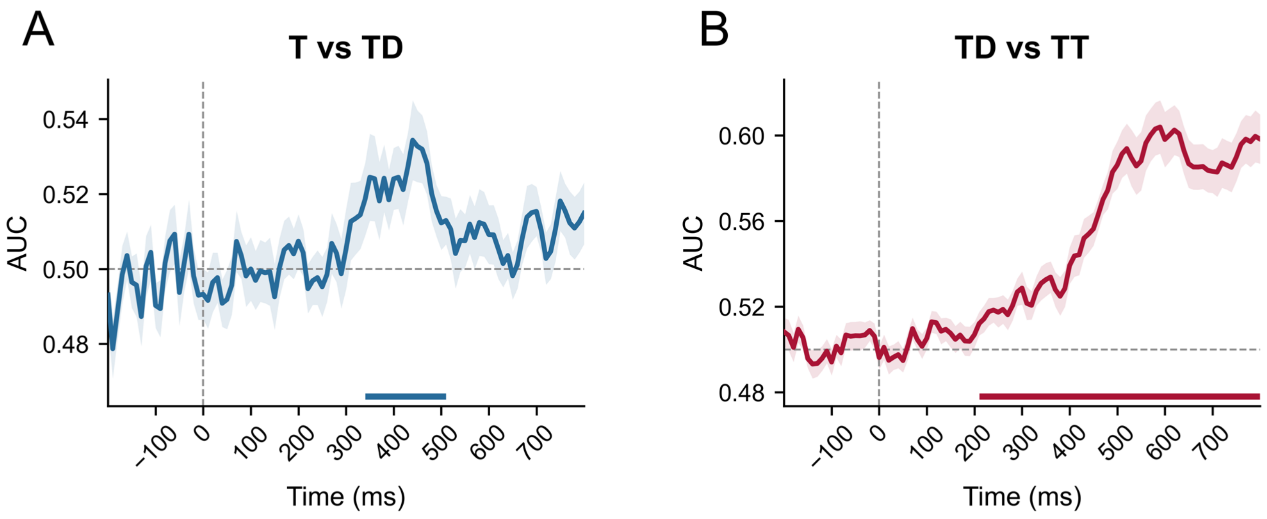


**Figure 3—figure supplement 1. Additional diagonal decoding of correct-report conditions.**

(**A**) Correct T versus TD: 340–510 ms ($p_{\text{cluster}} = .003$). (**B**) Correct TD versus TT: 210–800 ms ($p_{\text{cluster}} < .001$). Decoding performance is the area under the receiver operating characteristic curve (AUC). Lines show *M* with ± *SEM* shading. Solid colored bars mark significant clusters; the gray horizontal line marks chance (.50). No marginal cluster was detected in either comparison. $n = 29$; tests use 20,000 participant sign flips with temporal cluster correction, separately for each comparison. Complete diagonal results are in **Table S10**.

**Table S10. Pairwise diagonal decoding (Figures 3A–C and Figure 3—figure supplement 1).**

**A. Single-trial decoding of target load and intrusion outcomes.**

| Comparison | *n* | Time (ms) ($p_{\text{cluster}}$) |
|---|---|---|
| T vs TT | 29 | 300–800 (< **.001**) |
| TD vs DI_TD | 29 | 230–800 (< **.001**) |
| TD vs DI_D | 16 | 220–350 (**.005**); 420–470 (**.045**); 600–670 (**.045**); 690–740 (.069) |
| DI_TD vs DI_D | 16 | 260–380 (**.012**); 450–590 (**.010**) |
| TT vs DI_TD | 29 | 230–800 (< **.001**) |

***Note.*** Entries are time intervals in ms with $p_{\text{cluster}}$ in parentheses. Bold $p$ values are < .05; .05 ≤ $p$ < .10 denotes marginal results; a dash indicates no cluster with $p_{\text{cluster}}$ < .10. Diagonal tests use 20,000 one-sided participant sign flips with maximum-cluster correction over 0–800 ms, without additional Holm correction. Training classes were balanced; single-trial and trial-pair decoding used five and three folds, respectively, with 20 repeats.

**B. Additional single-trial condition comparisons.**

| Comparison | *n* | Time (ms) ($p_{\text{cluster}}$) |
|---|---|---|
| T vs DI_TD | 29 | 300–520 (**.003**); 580–800 (**.003**) |
| TT vs DI_D | 16 | 210–800 (< **.001**) |
| T vs DI_D | 16 | 260–380 (**.006**); 400–470 (**.019**); 490–530 (.090); 640–750 (**.012**) |
| T vs TD | 29 | 340–510 (**.003**) |
| TT vs TD | 29 | 210–800 (< **.001**) |

***Note.*** Formatting and tests follow **Table S10A**.

**C. Trial-pair sensitivity analysis.**

| Comparison | *n* | Time (ms) ($p_{\text{cluster}}$) |
|---|---|---|
| T vs TT | 29 | 310–800 (< **.001**) |
| TD vs DI_TD | 29 | 230–800 (< **.001**) |
| TD vs DI_D | 16 | 210–350 (**.006**); 420–470 (.065); 620–670 (.080) |
| DI_TD vs DI_D | 16 | 260–380 (**.009**); 450–550 (**.018**); 580–640 (.0505); 670–800 (**.019**) |
| TT vs DI_TD | 29 | 230–800 (< **.001**) |
| T vs DI_TD | 29 | 330–560 (**.003**); 580–800 (**.003**) |
| TT vs DI_D | 16 | 200–800 (< **.001**) |
| T vs DI_D | 16 | 200–470 (**.001**); 600–670 (**.042**); 710–780 (.056) |
| T vs TD | 29 | 340–490 (**.001**) |
| TT vs TD | 29 | 210–800 (< **.001**) |

***Note.*** Trial-pair sensitivity analysis; formatting and tests follow **Table S10A**.

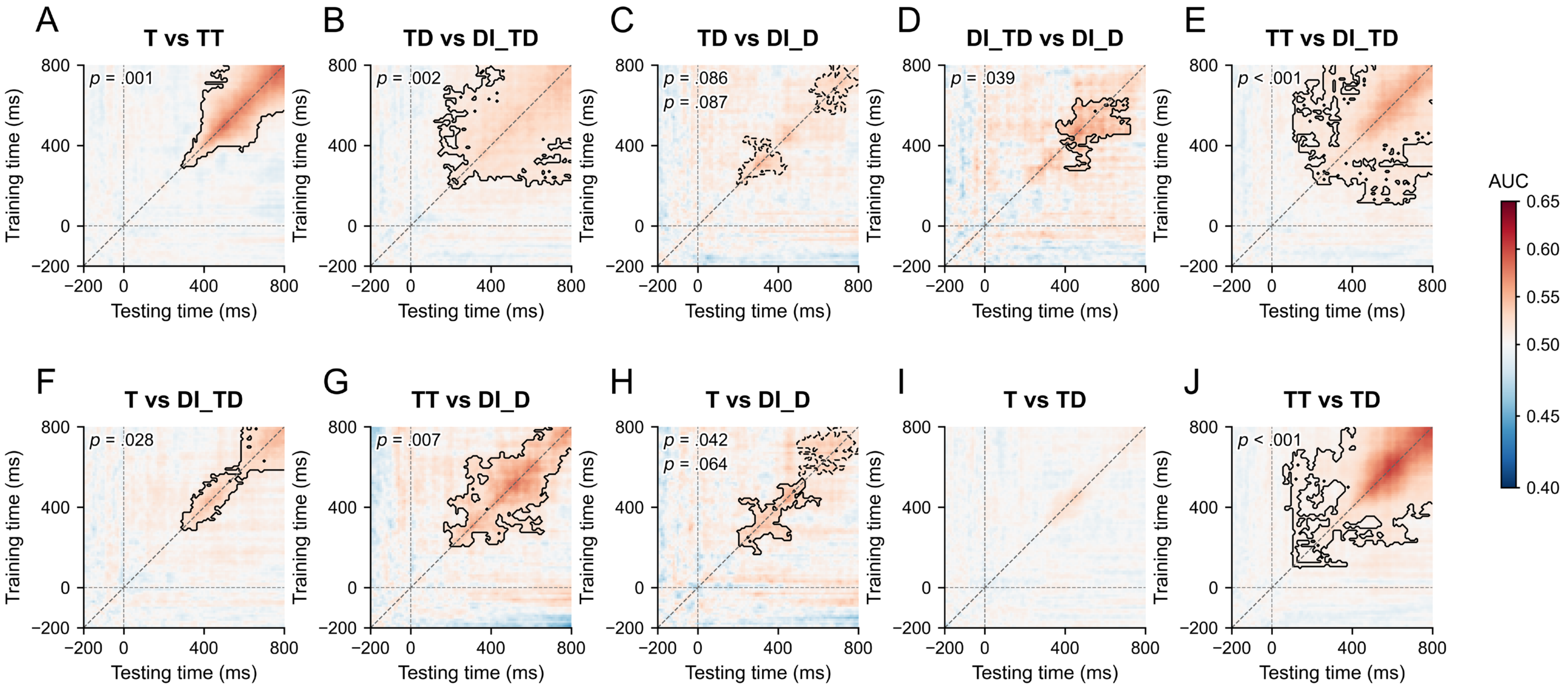


**Figure 3—figure supplement 2. Temporal generalization across target conditions and intrusion outcomes.**

AUC (*M*) for comparisons **A–J**. Solid contours show $p_{\text{cluster}} < .05$; dashed contours show $.05 \leq p_{\text{cluster}} < .10$. Cluster *p* values are printed in each panel. Gray diagonals mark matching training and testing times. Each −200–800 ms matrix is corrected separately; *n* = 16 for DI_D comparisons and 29 otherwise.

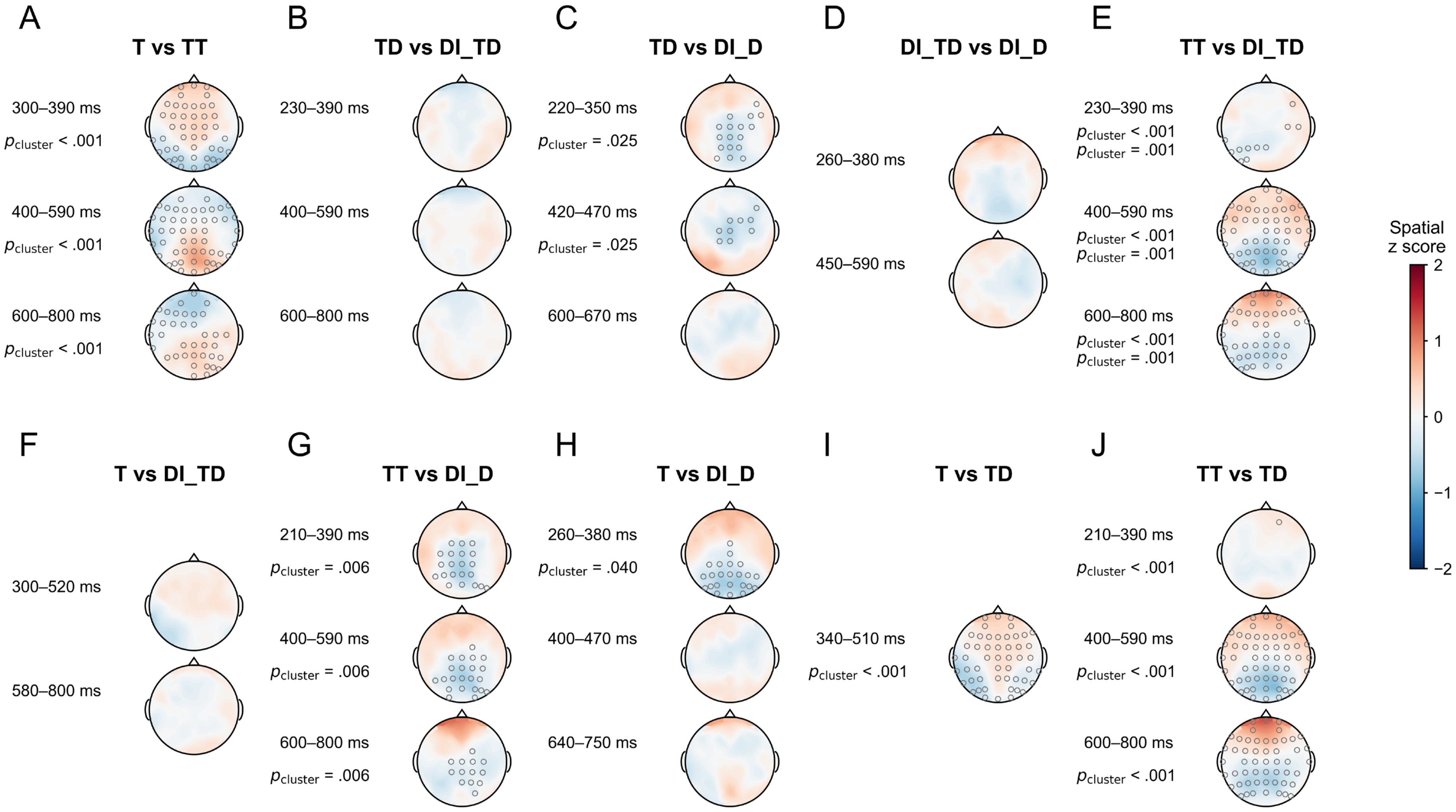


**Figure 3—figure supplement 3. Classification covariance patterns across target conditions and intrusion outcomes.**

Classification covariance maps are within-participant spatial $z$ scores; the second condition is the positive class. Small gray circles mark membership in a significant space–time cluster within the displayed interval. Annotated $p_{\text{cluster}}$ values apply to clusters, not individual electrodes. $n$ = 16 for DI_D comparisons and 29 otherwise.

**Table S11. Time-resolved T–TT reference-axis contrasts (Figures 3D and F).**

| Analysis | *n* | Time (ms) ($p_{cluster}$) |
|---|---|---|
| TD Relative distance | 29 | 470–800 (< **.001**) |
| DI_TD Relative distance | 29 | 500–620 (**.007**) |
| DI_D Relative distance | 16 | 490–590 (**.016**); 730–800 (**.032**) |
| DI_TD − TD Coordinate | 29 | 590–800 (< **.001**) |
| DI_TD − TD Relative distance | 29 | 630–800 (**.002**) |

***Note.*** Formatting follows **Table S10A**. Two-sided participant sign-flip tests control temporal clusters. Negative distance contrasts favor T; positive coordinate changes indicate a TT-directed shift.

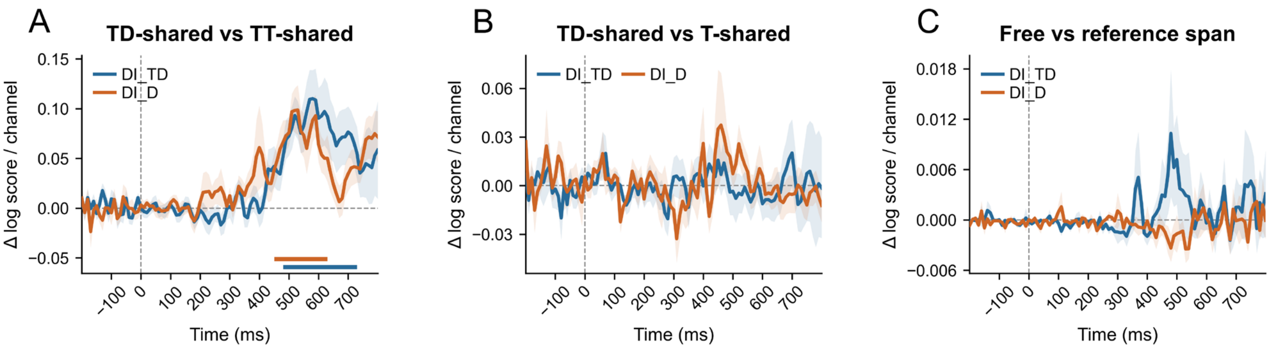


**Figure 3—figure supplement 4. Additional PCM comparisons of intrusion-related patterns.**

Pattern component modeling (PCM) compares held-out log scores; positive differences favor the first model. (**A**) TD-shared versus TT-shared: DI_TD, 480–730 ms ($n$ = 29, $p_{\text{cluster}}$ = .005); DI_D, 450–630 ms ($n$ = 16, $p_{\text{cluster}}$ = .012). Neither **B** (TD-shared versus T-shared) nor **C** (free versus reference span) had a cluster with $p_{\text{cluster}} < .10$. Tests jointly control eight primary contrasts, both samples, and time. Shading is ± *SEM*; solid and dashed bars mark significant and marginal clusters, respectively (**Table S12**).

**Table S12. PCM comparisons for DI_TD and DI_D (Figures 3E and Figure 3—figure supplement 4).**

| Intrusion | *n* | First vs second model | First favored: ms ($p_{\text{cluster}}$) | Second favored: ms ($p_{\text{cluster}}$) |
|---|---|---|---|---|
| DI_TD | 29 | TD vs T | — | — |
| DI_TD | 29 | TD vs TT | 480–730 (**.005**) | — |
| DI_TD | 29 | T vs TT | 480–680 (**.008**) | — |
| DI_TD | 29 | Free vs TD | — | 0–270 (< **.001**); 450–610 (**.028**) |
| DI_TD | 29 | Free vs T | — | 0–370 (< **.001**); 490–660 (**.008**) |
| DI_TD | 29 | Free vs TT | — | 0–370 (< **.001**) |
| DI_TD | 29 | Correct span vs One axis | — | 0–100 (**.039**) |
| DI_TD | 29 | Free vs Correct span | — | — |
| DI_D | 16 | TD vs T | — | — |
| DI_D | 16 | TD vs TT | 450–630 (**.012**) | — |
| DI_D | 16 | T vs TT | 480–630 (**.027**) | — |
| DI_D | 16 | Free vs TD | — | 0–240 (< **.001**); 570–800 (**.003**) |
| DI_D | 16 | Free vs T | — | 0–210 (< **.001**); 640–800 (**.007**) |
| DI_D | 16 | Free vs TT | — | 0–110 (**.003**); 130–200 (**.037**) |
| DI_D | 16 | Correct span vs One axis | — | — |
| DI_D | 16 | Free vs Correct span | — | — |

***Note.*** Formatting follows **Table S10A**. PCM jointly controls eight primary contrasts, both samples, and time. Columns identify the favored model. Shared models constrain covariance structure without implying identical activity.

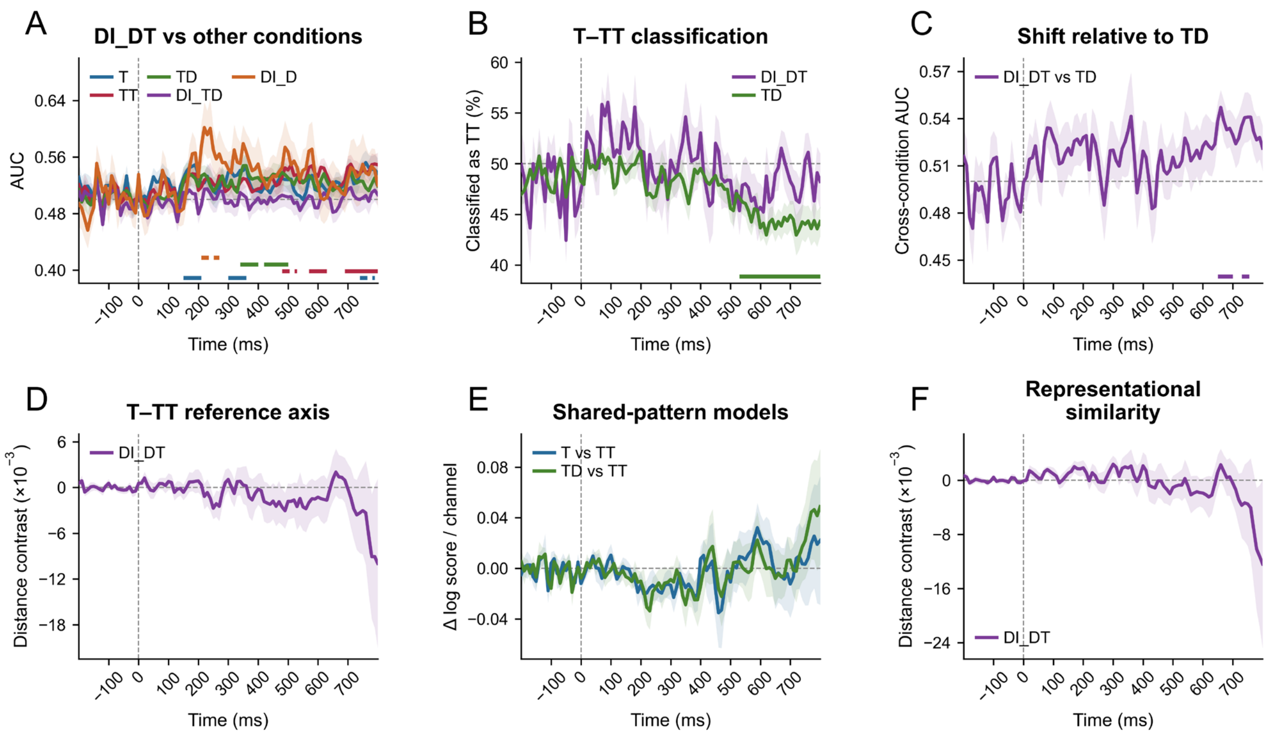


**Figure 3—figure supplement 5. Multivariate characterization of DI_DT.**

(**A**) DI_DT decoding versus T: 150–210 ms ($p_{\text{cluster}}$ = .023), 300–360 ms (.046), and 740–790 ms (.057); versus TT: 480–530 ms (.068), 570–630 ms (.037), and 690–800 ms (.010); versus TD: 340–400 ms (.043) and 420–500 ms (.032); versus DI_D: 210–270 ms (.051); versus DI_TD: no cluster with $p_{\text{cluster}} < .10$. (**B**) TT-classification proportions: TD, 530–800 ms ($p_{\text{cluster}} < .001$); DI_DT, no cluster with $p_{\text{cluster}} < .10$. (**C**) DI_DT versus TD on the T–TT decoder: 650–700 ms ($p_{\text{cluster}}$ = .034) and 730–770 ms (.076); AUC > .50 indicates a TT-directed shift. (**D–F**) Reference-axis preference, PCM sharing contrasts, and representational similarity analysis (RSA) preference had no cluster with $p_{\text{cluster}} < .10$. Negative distance contrasts favor T. $n$ = 18, except DI_D in A ($n$ = 11). Shading is ± *SEM*; solid and dashed bars mark significant and marginal clusters, respectively. PCM retains joint model correction; RSA preference is tested separately from absolute distances (**Tables S13–S14**).

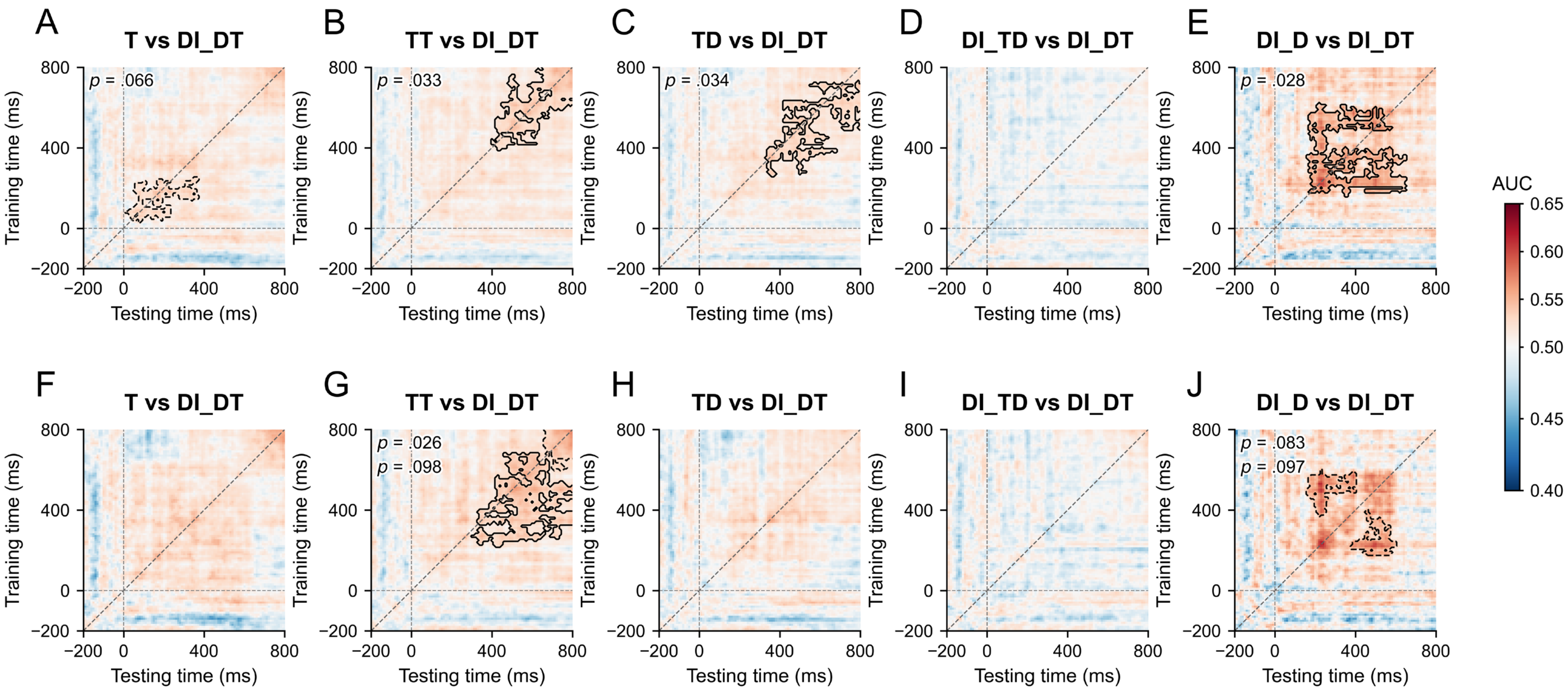


**Figure 3—figure supplement 6. Temporal generalization of DI_DT pairwise decoding.**

DI_DT versus T (**A, F**), TT (**B, G**), TD (**C, H**), DI_TD (**D, I**), and DI_D (**E, J**). The upper row uses single trials; the lower row uses independent trial pairs. Contour and *p*-value conventions follow **Figure 3—figure supplement 2**. Each −200–800 ms matrix is corrected separately. $n$ = 18, except DI_D comparisons ($n$ = 11).

**Table S13. DI_DT pairwise diagonal decoding (Figure 3—figure supplement 5A and Figure 3—figure supplement 7F).**

**A. Single-trial decoding.**

| Comparison | *n* | Time (ms) ($p_{cluster}$) |
|---|---|---|
| T vs DI_DT | 18 | 150–210 (**.023**); 300–360 (**.046**); 740–790 (.057) |
| TT vs DI_DT | 18 | 480–530 (.068); 570–630 (**.037**); 690–800 (**.010**) |
| TD vs DI_DT | 18 | 340–400 (**.043**); 420–500 (**.032**) |
| DI_TD vs DI_DT | 18 | — |
| DI_D vs DI_DT | 11 | 210–270 (.051) |

***Note.*** Formatting and decoding tests follow **Table S10A**; *n* = 18, except DI_DT–DI_D (*n* = 11).

**B. Trial-pair sensitivity analysis.**

| Comparison | *n* | Time (ms) ($p_{cluster}$) |
|---|---|---|
| T vs DI_DT | 18 | 150–200 (.074); 740–800 (.069) |
| TT vs DI_DT | 18 | 470–540 (**.044**); 570–630 (**.0497**); 700–800 (**.015**) |
| TD vs DI_DT | 18 | 340–390 (.075) |
| DI_TD vs DI_DT | 18 | — |
| DI_D vs DI_DT | 11 | 210–270 (.053) |

***Note.*** Trial-pair sensitivity analysis; conventions follow **Table S13A**.

**Table S14. Cross-condition and representational analyses of DI_DT (Figure 3—figure supplement 5B–F and Figure 3—figure supplement 7A–E).**

**A. Classification and reference-distance tests.**

| Analysis | Time (ms) ($p_{cluster}$) |
|---|---|
| Held-out T–TT AUC | 380–800 (< **.001**) |
| DI_DT TT-classification proportion | — |
| DI_DT vs TD: T–TT decoder AUC | 650–700 (**.034**); 730–770 (.076) |
| DI_DT − TD: TT-classification proportion | 650–680 (**.034**) |
| Reference-axis distance contrast | — |
| Reference distance to T | 160–200 (.084) |
| Reference distance to TT | 480–510 (.093); 750–800 (.053) |

***Note.*** Formatting follows **Table S10A**. Cross-decoder AUC uses within-participant label permutations; other contrasts use participant sign flips. Temporal cluster correction is retained. RSA distances jointly control three references, with preference tested separately.

**B. Axis-coordinate and RSA tests.**

| Analysis | Time (ms) ($p_{cluster}$) |
|---|---|
| DI_DT − TD: axis coordinate | — |
| DI_DT − TD: axis-orthogonal signal | 140–200 (**.035**) |
| RSA distance to T | 160–230 (.061) |
| RSA distance to TD | 430–510 (.062) |
| RSA distance to TT | 730–800 (.087) |
| RSA distance contrast | — |
| DI_DT − DI_D: axis coordinate ($n$ = 11) | 730–770 (.091) |
| DI_DT: axis-orthogonal signal | 160–200 (.075) |

***Note.*** Tests and formatting follow **Table S14A**; DI_DT–DI_D uses $n$ = 11.

**C. PCM sharing-model comparisons.**

| First vs second model | First favored: ms ($p_{cluster}$) | Second favored: ms ($p_{cluster}$) |
|---|---|---|
| TD vs T | — | — |
| TD vs TT | — | — |
| T vs TT | — | — |
| Free vs TD | — | 0–210 (**.003**) |
| Free vs T | — | 0–210 (**.004**); 520–650 (**.027**) |
| Free vs TT | — | 0–400 (< **.001**) |
| Correct span vs One axis | — | — |
| Free vs Correct span | — | — |
| T vs Null | 470–720 (**.015**) | 0–260 (< **.001**) |
| TT vs Null | 460–770 (**.005**) | 0–50 (**.037**); 80–200 (**.016**) |
| TD vs Null | 580–710 (.061) | 0–50 (**.034**); 80–290 (< **.001**) |
| One axis vs Null | 610–710 (.091) | 0–210 (< **.001**) |

***Note.*** Formatting follows **Table S10A**. Eight primary PCM contrasts and six model-versus-null diagnostics form separate corrected families. An early advantage over the free model need not indicate shared signal if both models underperform the null.

**D. T-shared versus TT-shared PCM sensitivity analysis across intrusion types.**

| Intrusion | *n* | Time (ms) ($p_{cluster}$) |
|---|---|---|
| DI_TD | 29 | 480–680 (**.018**) |
| DI_D | 16 | 480–630 (**.046**) |
| DI_DT | 18 | — |

***Note.*** Formatting follows **Table S10A**. Entries report intervals favoring the T-shared over the TT-shared model. This sensitivity test jointly controls three intrusion samples, eight model contrasts, and time, using a larger family than **Table S12**.

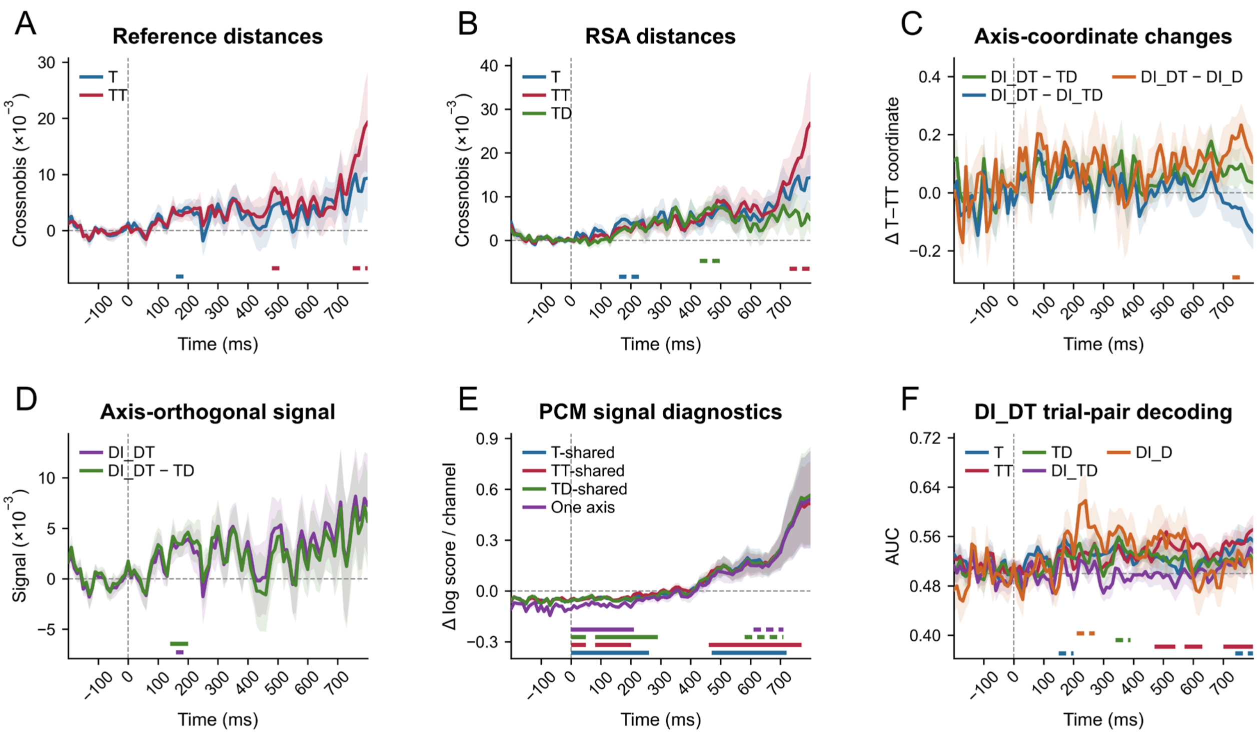


**Figure 3—figure supplement 7. Additional DI_DT reference tests and sensitivity analyses.**

(**A**) Reference distances: T, 160–200 ms ($p_{cluster}$ = .084); TT, 480–510 ms (.093) and 750–800 ms (.053). (**B**) RSA distances: T, 160–230 ms ($p_{cluster}$ = .061); TT, 730–800 ms (.087); TD, 430–510 ms (.062). (**C**) Axis shifts: DI_DT minus DI_D, 730–770 ms ($n$ = 11, $p_{cluster}$ = .091); the other contrasts had no cluster with $p_{cluster}$ < .10. (**D**) Axis-orthogonal signal: DI_DT, 160–200 ms ($p_{cluster}$ = .075); DI_DT minus TD, 140–200 ms (.035). (**E**) PCM versus null: T-shared was higher at 470–720 ms ($p_{cluster}$ = .015) and lower at 0–260 ms (< .001); TT-shared was higher at 460–770 ms (.005) and lower at 0–50 ms (.037) and 80–200 ms (.016); TD-shared was higher at 580–710 ms (.061) and lower at 0–50 ms (.034) and 80–290 ms (< .001); one-axis was higher at 610–710 ms (.091) and lower at 0–210 ms (< .001). (**F**) Trial-pair decoding versus T: 150–200 ms ($p_{cluster}$ = .074) and 740–800 ms (.069); versus TT: 470–540 ms (.044), 570–630 ms (.0497), and 700–800 ms (.015); versus TD: 340–390 ms (.075); versus DI_D: 210–270 ms (.053); versus DI_TD: no cluster with $p_{cluster}$ < .10. $n$ = 18, except DI_D comparisons ($n$ = 11). Shading is ± *SEM*; solid and dashed bars mark significant and marginal clusters, respectively. RSA distances jointly control three references; PCM null comparisons jointly control six models (**Tables S13–S14**).

### 4. Supplementary time-frequency results

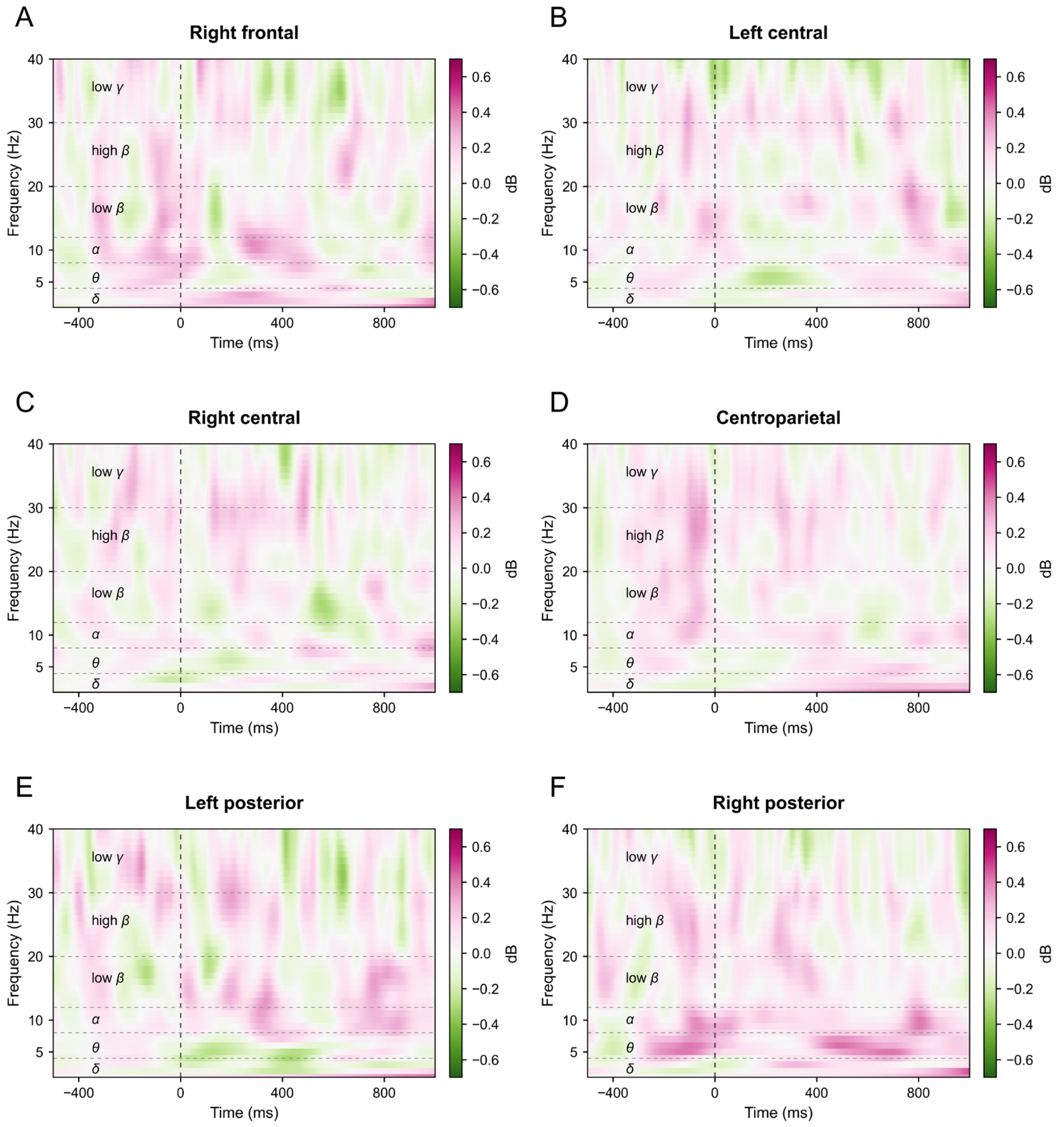


**Figure 4—figure supplement 1. Time–frequency power differences between correct TD and DI_TD trials outside the left frontal region.**

Correct TD minus DI_TD power ($n$ = 29) at right frontal (**A**), left central (**B**), right central (**C**), centroparietal (**D**), left posterior (**E**), and right posterior (**F**) electrodes. Power is expressed in decibels (dB) relative to −500 to −200 ms; magenta indicates greater TD power and green greater DI_TD power. Dashed lines mark T1 onset and frequency-band boundaries. Two-sided cluster-permutation tests used 20,000 randomizations over 1–40 Hz and −500–1,000 ms (cluster-forming $p < .05$). None of these regions had a significant cluster (all $p_{\text{cluster}} \geq .068$, $p_{\text{Holm}} \geq .410$). Holm correction covered seven regions, including the left frontal region in **Figure 4A**.

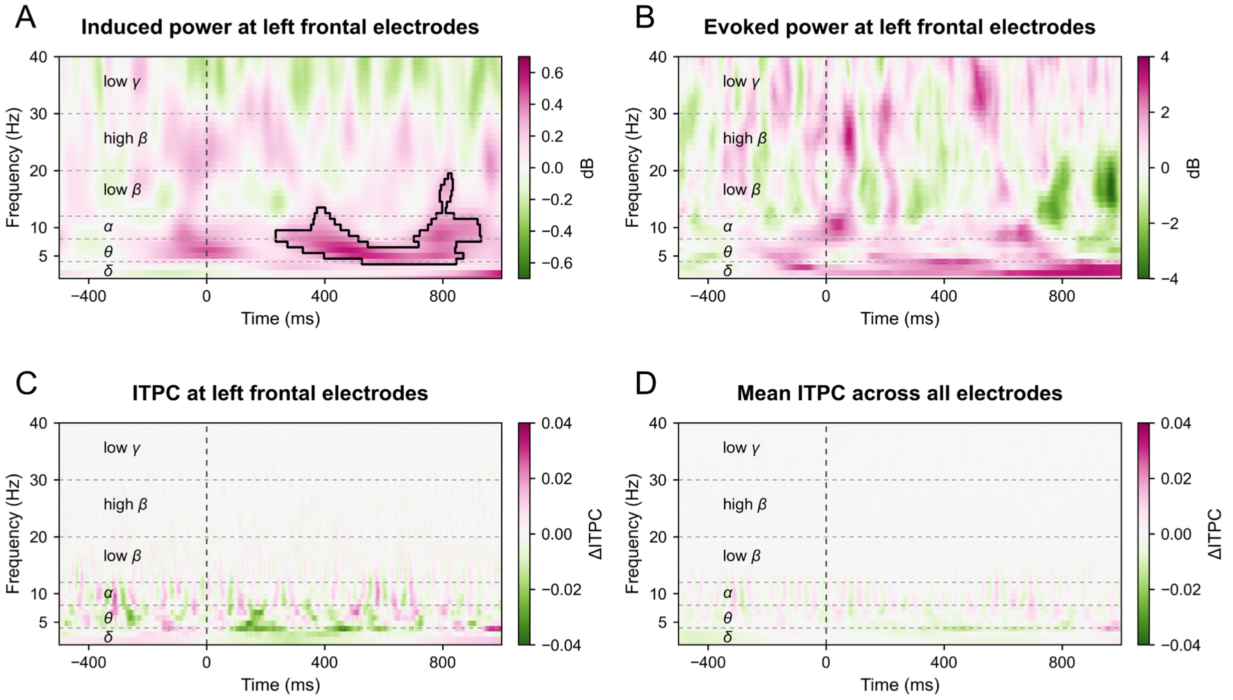


**Figure 4—figure supplement 2. Induced and evoked power and intertrial phase coherence for correct TD and DI_TD trials.**

Correct TD minus DI_TD differences ($n = 29$) in induced power (A), evoked power (B), and intertrial phase coherence (ITPC) at left frontal electrodes (C), and mean ITPC across electrodes (D). Induced power follows subtraction of the condition ERP from each trial; evoked power uses the condition ERP. Color conventions and the power baseline follow **Figure 4—figure supplement 1**; ITPC is unitless. The outline in A marks a significant cluster ($p_{\text{cluster}} = .006$), mainly in theta with extension into adjacent frequencies. The other measures had no significant cluster (all $p_{\text{cluster}} \geq .168$); each measure was tested separately over 1–40 Hz and −500 to 1,000 ms using 20,000 two-sided within-participant condition-label permutations, with cluster correction within each map and no additional Holm correction across the four measures.

**Table S15. Single-trial frontal-power associations with DI_TD (Figure 4C–E).**

| Model | Predictor | *b* | *SE* | *z* | *p* | OR | 95% CI for OR |
|---|---|---|---|---|---|---|---|
| M1 | Intercept | −0.80 | 0.23 | −3.52 | < **.001** | 0.45 | [0.29, 0.70] |
| M1 | Left frontal power | −0.10 | 0.03 | −3.78 | < **.001** | 0.91 | [0.86, 0.95] |
| M2 | Intercept | −0.83 | 0.23 | −3.60 | < **.001** | 0.44 | [0.28, 0.69] |
| M2 | Left − right frontal power | −0.12 | 0.03 | −4.63 | < **.001** | 0.89 | [0.85, 0.93] |
| M2 | Target location (right vs left) | 0.05 | 0.05 | 0.93 | .350 | 1.05 | [0.95, 1.16] |
| M2 | Bilateral mean power | −0.02 | 0.03 | −0.96 | .337 | 0.98 | [0.93, 1.03] |
| M3 | Intercept | −0.83 | 0.23 | −3.60 | < **.001** | 0.44 | [0.28, 0.69] |
| M3 | Left frontal power | −0.12 | 0.03 | −4.38 | < **.001** | 0.89 | [0.84, 0.94] |
| M3 | Right frontal power | 0.09 | 0.03 | 3.31 | < **.001** | 1.10 | [1.04, 1.16] |
| M3 | Target location (right vs left) | 0.05 | 0.05 | 0.93 | .350 | 1.05 | [0.95, 1.16] |

***Note.*** Binomial GLMMs use 29 participants, with DI_TD = 1 and TD = 0. All include a participant random intercept. M1 uses the full left frontal cluster; M2 and M3 use the full hemisphere-interaction cluster (**Figure 4A–B**). M2 includes left-minus-right power, bilateral mean power, and location; M3 includes both hemispheres and location. Power is centered within participant and scaled by its pooled *SD*; M3 uses a common hemispheric scale. *p* values are unadjusted two-sided Wald tests with 95% Wald CIs. All models converged without singular fits.

**Table S16. Nested frontal-power models for DI_TD (Figure 4E).**

| Reduced model | Added predictor | $\chi^2$ | *df* | *p* | ΔAIC |
|---|---|---|---|---|---|
| Left power + target location | Right power | 10.93 | 1 | **< .001** | 8.93 |
| Right power + target location | Left power | 19.22 | 1 | **< .001** | 17.22 |

***Note.*** Nested models use the same interaction-cluster trials, location covariate, and participant random intercept as M3 in **Table S15**. *p* values are unadjusted likelihood-ratio tests. AIC is the Akaike information criterion; ΔAIC is reduced minus full-model AIC, so positive values favor the full model.

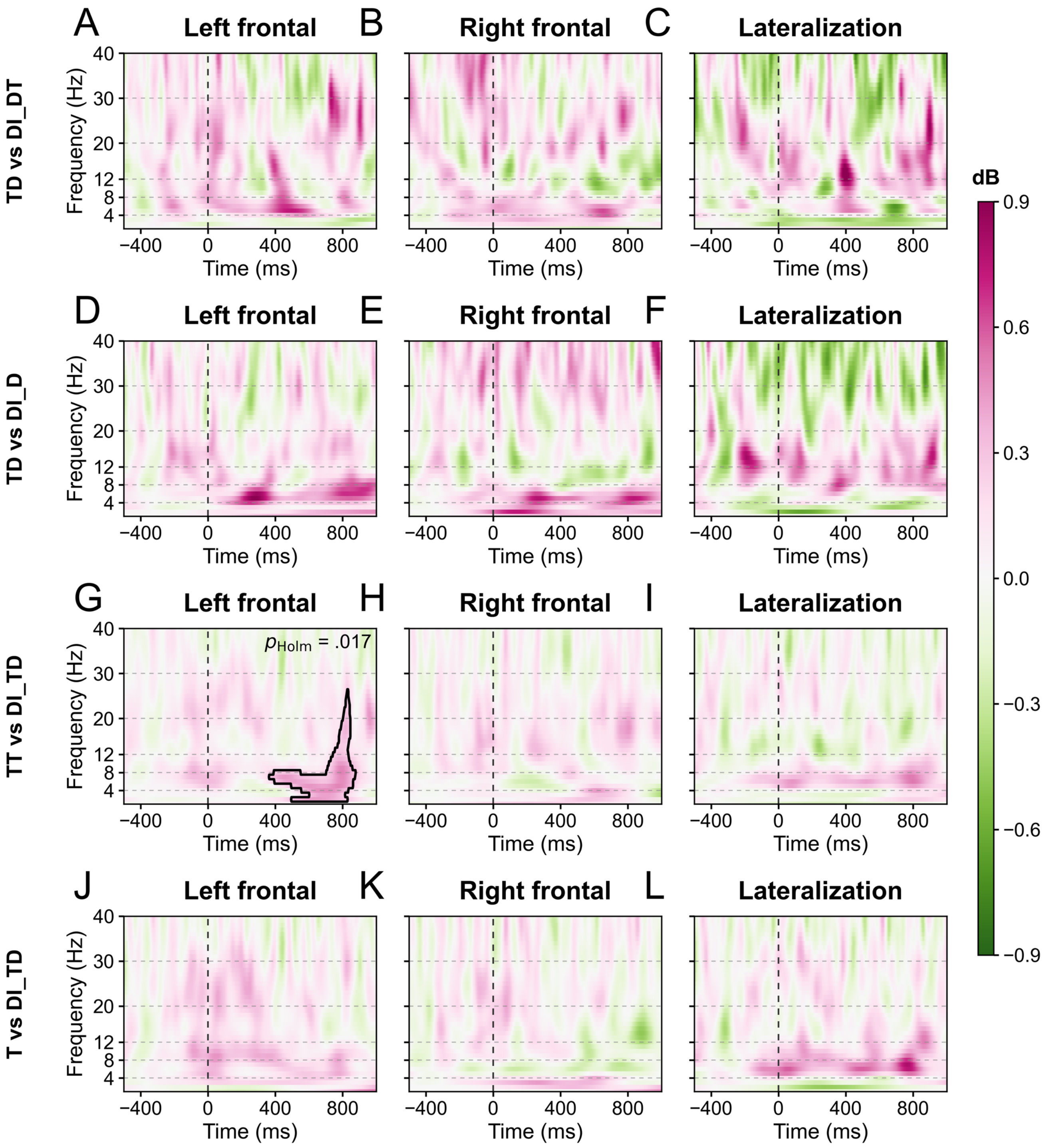


**Figure 4—figure supplement 3. Supplementary frontal-power comparisons for intrusion outcomes.**

Total-power differences compare correct TD with DI_DT (**A–C**; $n = 18$) and DI_D (**D–F**; $n = 16$), and correct TT (**G–I**; $n = 29$) or T (**J–L**; $n = 29$) with DI_TD. Columns show left frontal (lFC), right frontal (rFC), and frontal lateralization. The two regional measures average seven electrodes each; lateralization uses five homologous pairs (F3/F4, F5/F6, F7/F8, FC3/FC4, FC5/FC6) and represents the between-condition difference in left-minus-right power. Maps subtract the second condition from the first; colors and baseline follow **Figure 4—figure supplement 1**. Tests used 20,000 two-sided paired cluster permutations over 1–40 Hz and −500 to 1,000 ms. Holm correction covers the seven comparisons in **Figure 4—figure supplements 3–4** separately within each measure. Only corrected-significant clusters are outlined: TT minus DI_TD at lFC (**G**; $p_{Holm} = .017$; **Table S17**).

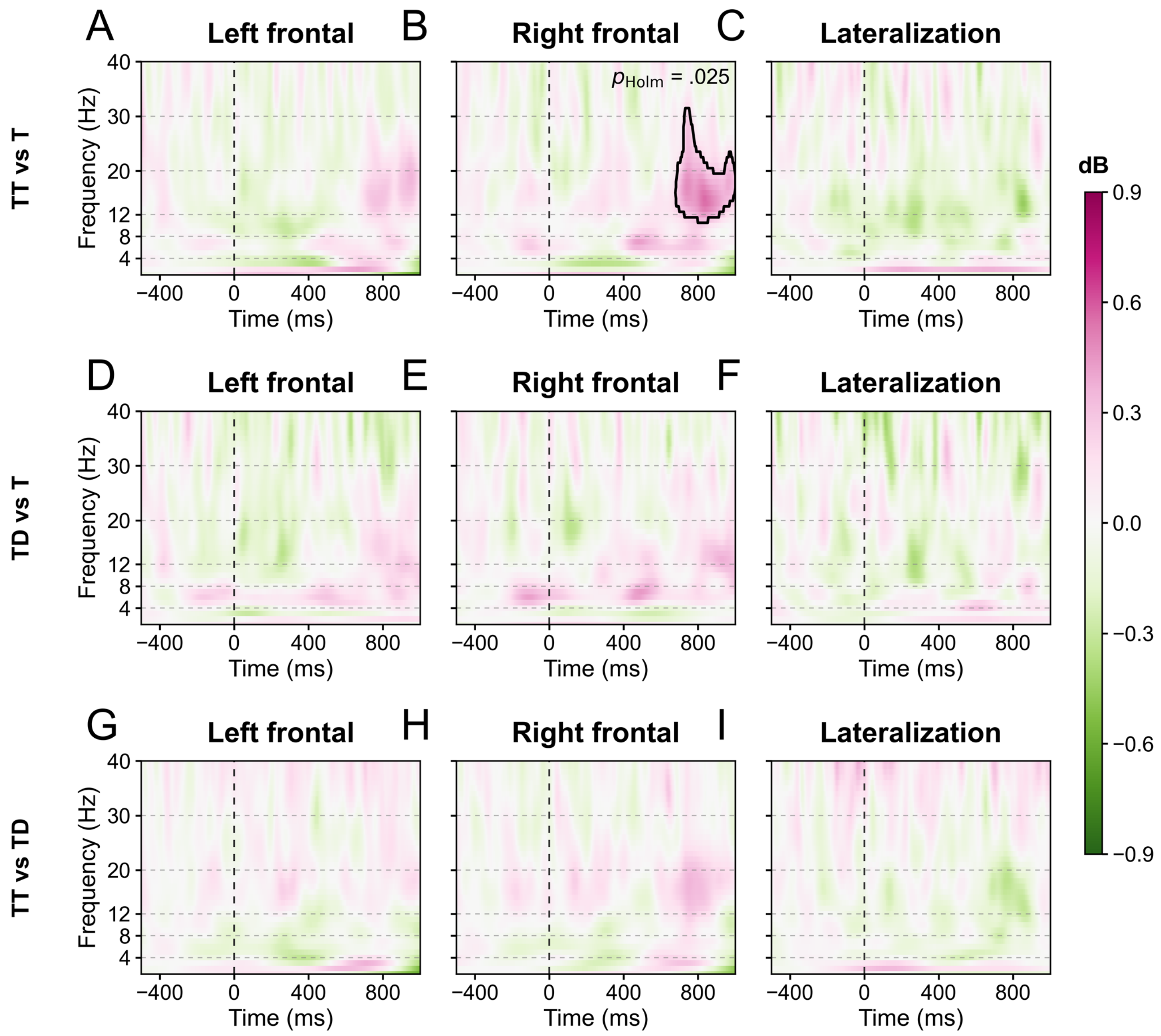


**Figure 4—figure supplement 4. Frontal-power comparisons among correct-report conditions.**

Correct-report power differences ($n$ = 29): TT minus T (**A–C**), TD minus T (**D–F**), and TT minus TD (**G–I**). Measures, colors, baseline, and statistical procedures follow **Figure 4—figure supplement 3**. Only the TT-minus-T rFC cluster survived correction (**B**; $p_{\mathrm{cluster}}$ = .0036, $p_{\mathrm{Holm}}$ = .025; **Table S17**). No other regional or lateralization test was significant after correction.

**Table S17. Supplementary frontal time–frequency differences surviving Holm correction (Figure 4—figure supplements 3–4).**

| Comparison | $n$ | Measure | Frequency (Hz) | Time (ms) | $p_{\text{cluster}}$ | $p_{\text{Holm}}$ |
|---|---|---|---|---|---|---|
| TT − DI_TD | 29 | lFC | 2.00–26.00 | 364.00–876.00 | **.0025** | **.017** |
| TT − T | 29 | rFC | 11.00–31.00 | 680.00–1000.00 | **.0036** | **.025** |

***Note.*** Only corrected-significant clusters are listed. $p_{\text{cluster}}$ controls time–frequency comparisons within each map; $p_{\text{Holm}}$ additionally adjusts the seven condition comparisons separately within lFC, rFC, and lateralization, using each map's minimum cluster $p$ value. All other tests had $p_{\text{Holm}} \geq .278$. Frequency and time values are outer bounds of irregular clusters, not uniformly significant rectangles. Definitions and sample sizes follow **Figure 4—figure supplements 3–4**. Significant $p$ values are bold.

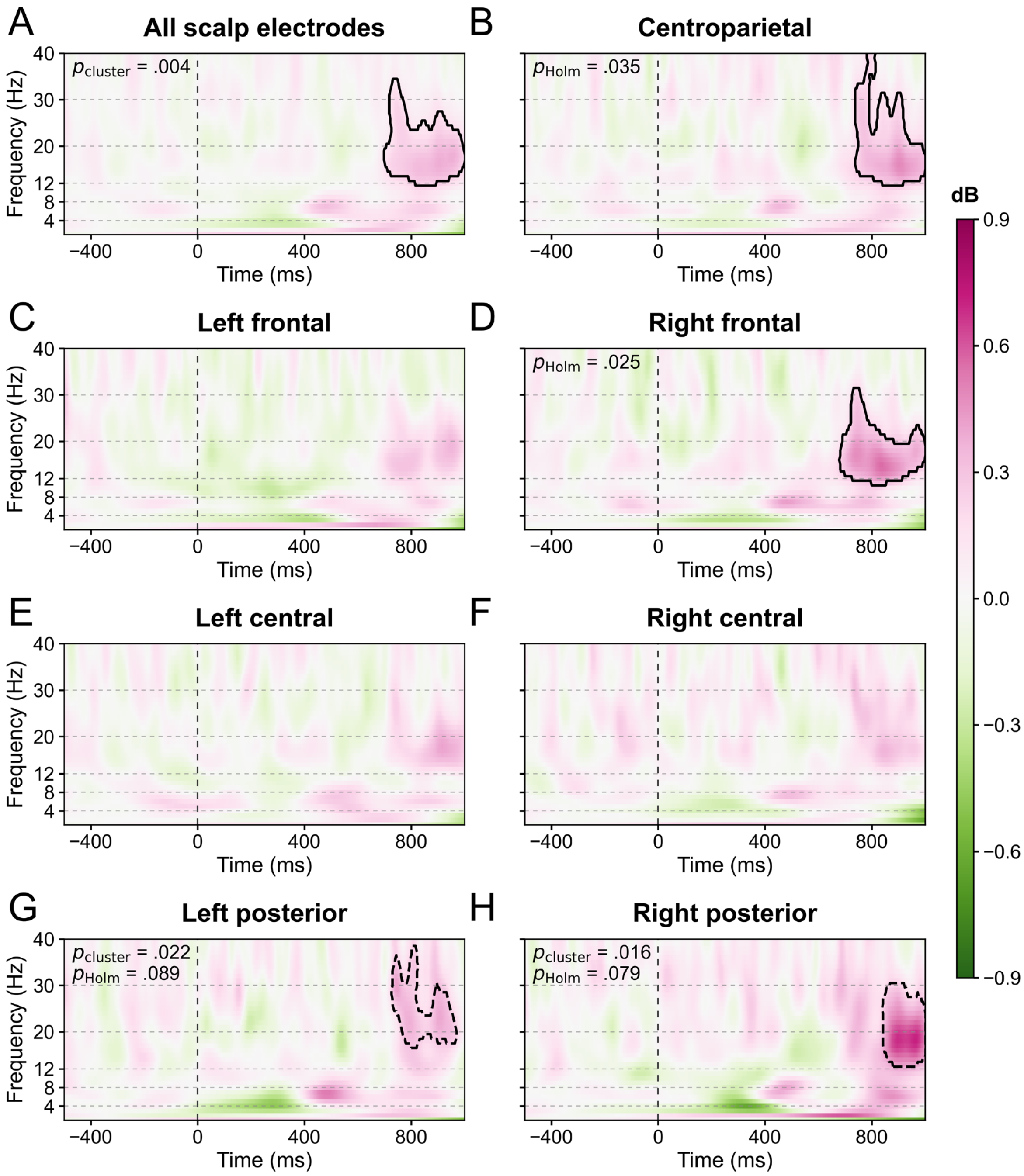


**Figure 4—figure supplement 5. Total-power differences between correct TT and T trials across the scalp.**

Correct TT minus correct T total power ($n$ = 29): all 60 scalp electrodes (**A**), centroparietal (**B**), left and right frontal (**C–D**), left and right central (**E–F**), and left and right posterior (**G–H**) regions. Colors and baseline follow **Figure 4—figure supplement 1**. Tests used 20,000 two-sided cluster permutations over 1–40 Hz and −500 to 1,000 ms. Holm correction covers seven regions; the all-electrode test uses cluster correction only. Solid outlines mark significant clusters: all electrodes, 12–34 Hz and 700–1,000 ms ($p_{cluster}$ = .0039); centroparietal, 12–40 Hz and 740–1,000 ms ($p_{cluster}$ = .0059, $p_{Holm}$ = .035); right frontal, 11–31 Hz and 680–1,000 ms ($p_{cluster}$ = .0036, $p_{Holm}$ = .025). Dashed outlines mark clusters with $p_{cluster} < .05$ that did not survive Holm correction: left posterior, 17–38 Hz and 728–968 ms ($p_{cluster}$ = .022, $p_{Holm}$ = .089); right posterior, 13–30 Hz and 840–1,000 ms ($p_{cluster}$ = .016, $p_{Holm}$ = .079). Ranges are outer bounds of irregular clusters, not uniformly significant rectangles.

### 5. Supplementary results for joint neural correlates

A joint multinomial GLMM included five ERP amplitudes, lFC power, rFC power, numerical distance, and target location, with correct TD as the reference and DI_TD, DI_D, and DI_DT as the other outcomes. Other errors were excluded. The full sample comprised 29 participants. The sensitivity sample comprised 11 participants with at least 15 valid trials for DI_D and DI_DT. Neural predictors were centered within participant over retained TD trials and scaled using the full-sample pooled centered *SD*; distance was standardized using the full-sample trial mean and *SD*. Neither model included participant mean neural values. The lFC and rFC features used the regional and frontal-lateralization masks, respectively. The same centering and scaling were retained in the sensitivity analysis.

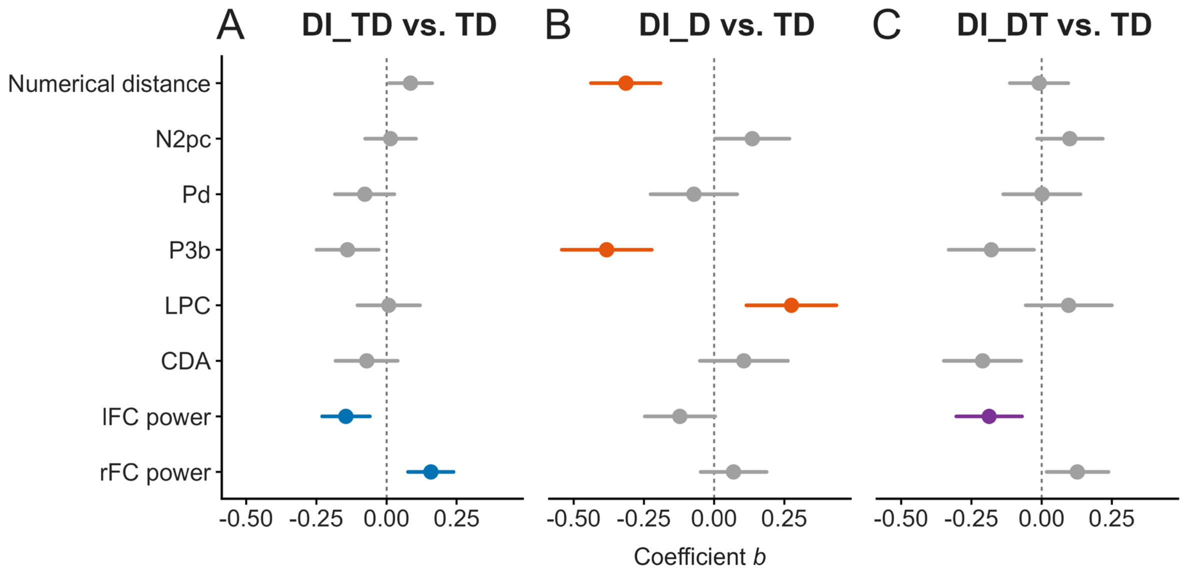


**Figure 5—figure supplement 1. Joint neural associations in the intersection sample.**

The 11-participant sensitivity analysis follows **Figure 5**. Panels **(A–C)** show DI_TD, DI_D, and DI_DT relative to correct TD. Target location is controlled but not shown; numerical distance appears first. Points show *b* and lines show unadjusted 95% CIs. Colored estimates survive Holm correction across 27 slopes. The coefficient axes match **Figure 5**. Full statistics are reported in **Table S18B**.

**Table S18. Joint multinomial associations with intrusion outcomes (Figures 5 and Figure 5—figure supplement 1).**

**A. Main sample ($n$ = 29).**

| Outcome | Predictor | $b$ | $SE$ | $z$ | $p$ | $p_{\text{Holm}}$ | OR | 95% CI for OR |
|---|---|---|---|---|---|---|---|---|
| DI_TD | Numerical distance | −0.01 | 0.03 | −0.47 | .640 | 1.000 | 0.99 | [0.94, 1.04] |
| DI_TD | N2pc | −0.05 | 0.03 | −1.69 | .092 | .916 | 0.95 | [0.89, 1.01] |
| DI_TD | Pd | −0.05 | 0.04 | −1.36 | .174 | 1.000 | 0.95 | [0.89, 1.02] |
| DI_TD | P3b | −0.09 | 0.04 | −2.38 | **.017** | .277 | 0.92 | [0.85, 0.98] |
| DI_TD | LPC | 0.05 | 0.04 | 1.42 | .157 | 1.000 | 1.05 | [0.98, 1.13] |
| DI_TD | CDA | 0.03 | 0.03 | 0.87 | .385 | 1.000 | 1.03 | [0.96, 1.10] |
| DI_TD | lFC power | −0.13 | 0.03 | −4.76 | **< .001** | **< .001** | 0.88 | [0.83, 0.93] |
| DI_TD | rFC power | 0.09 | 0.03 | 3.34 | **< .001** | **.018** | 1.10 | [1.04, 1.15] |
| DI_D | Numerical distance | −0.31 | 0.04 | −7.05 | **< .001** | **< .001** | 0.73 | [0.67, 0.80] |
| DI_D | N2pc | 0.17 | 0.05 | 3.42 | **< .001** | **.014** | 1.19 | [1.08, 1.31] |
| DI_D | Pd | −0.07 | 0.06 | −1.17 | .240 | 1.000 | 0.93 | [0.84, 1.05] |
| DI_D | P3b | −0.37 | 0.06 | −5.98 | **< .001** | **< .001** | 0.69 | [0.62, 0.78] |
| DI_D | LPC | 0.32 | 0.06 | 5.34 | **< .001** | **< .001** | 1.38 | [1.23, 1.55] |
| DI_D | CDA | 0.03 | 0.05 | 0.51 | .612 | 1.000 | 1.03 | [0.93, 1.13] |
| DI_D | lFC power | −0.10 | 0.05 | −2.19 | **.028** | .396 | 0.90 | [0.83, 0.99] |
| DI_D | rFC power | 0.04 | 0.04 | 0.91 | .365 | 1.000 | 1.04 | [0.95, 1.13] |
| DI_DT | Numerical distance | −0.08 | 0.04 | −1.90 | .057 | .682 | 0.92 | [0.84, 1.00] |
| DI_DT | N2pc | 0.10 | 0.05 | 1.89 | .059 | .682 | 1.10 | [1.00, 1.22] |
| DI_DT | Pd | −0.01 | 0.06 | −0.11 | .912 | 1.000 | 0.99 | [0.88, 1.12] |
| DI_DT | P3b | −0.18 | 0.06 | −2.70 | **.007** | .123 | 0.84 | [0.74, 0.95] |
| DI_DT | LPC | 0.14 | 0.06 | 2.15 | **.031** | .408 | 1.15 | [1.01, 1.31] |
| DI_DT | CDA | −0.14 | 0.06 | −2.48 | **.013** | .225 | 0.87 | [0.78, 0.97] |
| DI_DT | lFC power | −0.15 | 0.05 | −3.17 | **.002** | **.031** | 0.86 | [0.78, 0.94] |
| DI_DT | rFC power | 0.11 | 0.05 | 2.26 | **.024** | .356 | 1.11 | [1.01, 1.22] |

***Note.*** Correct TD is the reference category. Both models jointly include the eight displayed predictors and target location, which is omitted from the table. Raw $p$ values are from two-sided Wald $z$ tests; $p_{\text{Holm}}$ adjusts all 27 slopes separately within each sample. OR CIs are unadjusted 95% Wald intervals. Neural coefficients describe a one-*SD* increase relative to the participant's mean; distance coefficients describe a one-*SD* increase in numerical distance. Centering and scaling follow **Figure 5**. Significant raw or adjusted $p$ values are bold.

**B. Intersection sample (*n* = 11).**

| Outcome | Predictor | *b* | *SE* | *z* | *p* | $p_{Holm}$ | OR | 95% CI for OR |
|---|---|---|---|---|---|---|---|---|
| DI_TD | Numerical distance | 0.09 | 0.04 | 2.19 | **.028** | .479 | 1.09 | [1.01, 1.18] |
| DI_TD | N2pc | 0.01 | 0.05 | 0.31 | .755 | 1.000 | 1.01 | [0.93, 1.11] |
| DI_TD | Pd | −0.08 | 0.05 | −1.43 | .152 | 1.000 | 0.93 | [0.83, 1.03] |
| DI_TD | P3b | −0.14 | 0.06 | −2.46 | **.014** | .279 | 0.87 | [0.78, 0.97] |
| DI_TD | LPC | 0.01 | 0.06 | 0.14 | .888 | 1.000 | 1.01 | [0.90, 1.13] |
| DI_TD | CDA | −0.07 | 0.06 | −1.25 | .212 | 1.000 | 0.93 | [0.83, 1.04] |
| DI_TD | lFC power | −0.14 | 0.04 | −3.32 | **< .001** | **.020** | 0.87 | [0.79, 0.94] |
| DI_TD | rFC power | 0.16 | 0.04 | 3.82 | **< .001** | **.003** | 1.17 | [1.08, 1.27] |
| DI_D | Numerical distance | −0.31 | 0.06 | −4.97 | **< .001** | **< .001** | 0.73 | [0.65, 0.83] |
| DI_D | N2pc | 0.14 | 0.07 | 2.01 | **.045** | .712 | 1.15 | [1.00, 1.31] |
| DI_D | Pd | −0.07 | 0.08 | −0.91 | .363 | 1.000 | 0.93 | [0.80, 1.09] |
| DI_D | P3b | −0.38 | 0.08 | −4.68 | **< .001** | **< .001** | 0.68 | [0.58, 0.80] |
| DI_D | LPC | 0.28 | 0.08 | 3.37 | **< .001** | **.018** | 1.32 | [1.12, 1.55] |
| DI_D | CDA | 0.11 | 0.08 | 1.33 | .185 | 1.000 | 1.11 | [0.95, 1.30] |
| DI_D | lFC power | −0.12 | 0.06 | −1.90 | .057 | .854 | 0.89 | [0.78, 1.00] |
| DI_D | rFC power | 0.07 | 0.06 | 1.16 | .244 | 1.000 | 1.07 | [0.95, 1.21] |
| DI_DT | Numerical distance | −0.01 | 0.05 | −0.16 | .871 | 1.000 | 0.99 | [0.89, 1.10] |
| DI_DT | N2pc | 0.10 | 0.06 | 1.69 | .091 | 1.000 | 1.11 | [0.98, 1.24] |
| DI_DT | Pd | 0.00 | 0.07 | 0.02 | .986 | 1.000 | 1.00 | [0.87, 1.15] |
| DI_DT | P3b | −0.18 | 0.08 | −2.31 | **.021** | .397 | 0.84 | [0.72, 0.97] |
| DI_DT | LPC | 0.10 | 0.08 | 1.24 | .216 | 1.000 | 1.10 | [0.95, 1.28] |
| DI_DT | CDA | −0.21 | 0.07 | −3.00 | **.003** | .058 | 0.81 | [0.71, 0.93] |
| DI_DT | lFC power | −0.19 | 0.06 | −3.12 | **.002** | **.039** | 0.83 | [0.74, 0.93] |
| DI_DT | rFC power | 0.13 | 0.06 | 2.27 | **.023** | .420 | 1.14 | [1.02, 1.27] |

***Note.*** Model specification, scaling, and statistical reporting follow part **A**. This sample meets both condition-specific trial-count criteria.

**LPC association conditional on P3b**

The positive LPC association with DI_D emerged only after controlling for P3b in the matched-trial multinomial models (**Table S19**). All full-sample models used the same trials from 29 participants, response categories, participant random-intercept structure, and neural scaling. LPC was not significant when entered alone ($b$ = 0.04, $p$ = .277), with numerical distance and target location ($b$ = 0.05, $p$ = .207), or with all full-model predictors except P3b ($b$ = 0.05, $p$ = .181). Adding P3b to the behavioral-covariate model yielded a positive LPC coefficient ($b$ = 0.33, $p$ < .001); the full model produced a similar estimate ($b$ = 0.32, $p$ < .001). The same pattern occurred in the 11-participant intersection sample. P3b and LPC were positively correlated after within-participant centering ($r$ = .745 in the full sample; .739 in the intersection sample), whereas their full-model DI_D coefficients had opposite signs. This pattern is consistent with statistical suppression: accounting for shared P3b-related variance reveals an association with relatively greater late positivity at a given P3b amplitude. It therefore characterizes the balance between earlier and later activity rather than a general LPC increase in substitution reports. In particular, it does not imply that DI_D elicited a larger condition-average LPC, that more information was maintained, or that P3b causally mediated LPC. Full-model variance inflation factors were modest (P3b = 2.26, LPC = 2.25), and all sensitivity models converged.

**Table S19. Sensitivity of the DI_D–LPC association to P3b adjustment.**

| n | Model | *b* | *SE* | *p* | $p_{\text{Holm}}$ | OR [95% CI] |
|---|---|---|---|---|---|---|
| 29 | LPC only | 0.04 | 0.04 | .277 | 1.000 | 1.04 [0.97, 1.13] |
| 29 | LPC + behavior | 0.05 | 0.04 | .207 | 1.000 | 1.05 [0.97, 1.14] |
| 29 | P3b + LPC + behavior | 0.33 | 0.06 | **< .001** | **< .001** | 1.39 [1.23, 1.56] |
| 29 | Full without P3b | 0.05 | 0.04 | .181 | 1.000 | 1.06 [0.97, 1.14] |
| 29 | Full | 0.32 | 0.06 | **< .001** | **< .001** | 1.38 [1.23, 1.55] |
| 11 | LPC only | −0.02 | 0.05 | .771 | 1.000 | 0.98 [0.88, 1.10] |
| 11 | LPC + behavior | −0.01 | 0.06 | .839 | 1.000 | 0.99 [0.89, 1.10] |
| 11 | P3b + LPC + behavior | 0.28 | 0.08 | **< .001** | **.005** | 1.32 [1.12, 1.55] |
| 11 | Full without P3b | −0.01 | 0.06 | .921 | 1.000 | 0.99 [0.89, 1.11] |
| 11 | Full | 0.28 | 0.08 | **< .001** | **.005** | 1.32 [1.12, 1.55] |

***Note.*** Each row reports the LPC slope for DI_D relative to correct TD from a four-category multinomial GLMM. The 29- and 11-participant samples contained 10,093 and 3,839 trials, respectively; every model within a sample used exactly the same trials and full-sample standardization. “Behavior” includes numerical distance and target location. “Full” also includes N2pc, Pd, P3b, CDA, and left and right frontal power. Models without P3b retain every other full-model predictor. Models were fitted using penalized quasi-likelihood with a maximum-likelihood approximation and category-specific participant random intercepts. Two-sided Wald p values are raw; $p_{\text{Holm}}$ adjusts the 10 displayed LPC tests. OR CIs are unadjusted 95% Wald intervals. The primary full-model correction across all 27 slopes is retained in **Table S18**; this sensitivity table uses its own explicitly defined family.

### 6. Additional ERP analyses

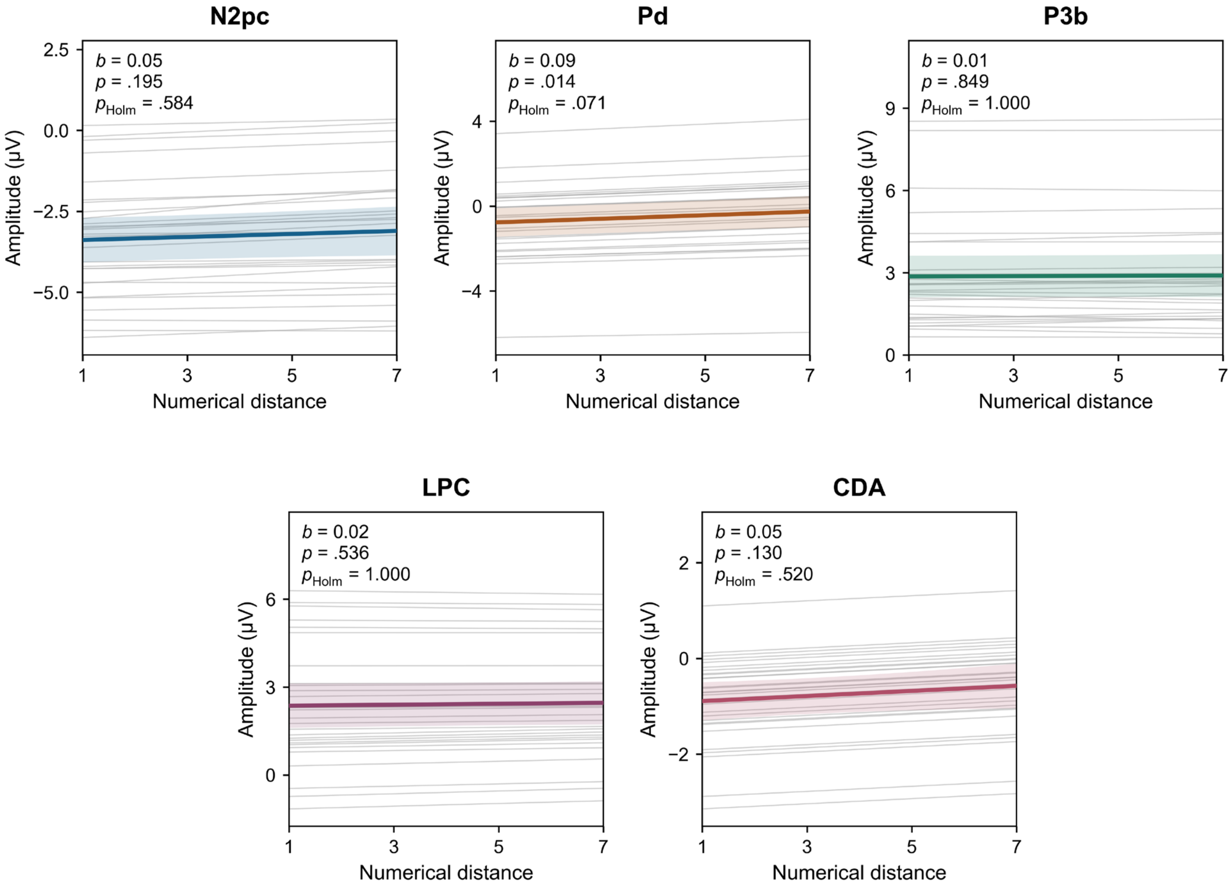


**Figure 2—figure supplement 9. Exploratory trial-level numerical-distance effects on ERP amplitudes.**

Linear mixed-effects models (LMMs) related absolute target–PTD numerical distance to N2pc, Pd, P3b, LPC, and CDA amplitudes in all artifact-free TD trials ($n = 29$). Models jointly adjusted for target location and report type (correct TD, DI_TD, DI_D, DI_DT, or other error). Colored lines and shading show fixed-effect predictions and unadjusted pointwise 95% CIs, standardized over the observed location and report-type distributions with equal participant weights. Gray lines show participant-specific predictions, adding each participant's random effects to the same covariate profile. Annotations show $b$, raw $p$, and $p_{\mathrm{Holm}}$ across five components. Distance was positively associated with Pd before correction ($p = .014$), but no component survived Holm correction (all $p_{\mathrm{Holm}} \geq .071$; **Table S20**).

**Table S20. Trial-level numerical-distance effects on ERP amplitudes (Figure 2—figure supplement 9).**

| Component | $b$ | $SE$ | $t$ | $p$ | $p_{Holm}$ | 95% CI for $b$ |
|---|---|---|---|---|---|---|
| N2pc | 0.05 | 0.04 | 1.33 | .195 | .584 | [−0.03, 0.12] |
| Pd | 0.09 | 0.03 | 2.61 | **.014** | .071 | [0.02, 0.15] |
| P3b | 0.01 | 0.03 | 0.19 | .849 | 1.000 | [−0.05, 0.06] |
| LPC | 0.02 | 0.03 | 0.63 | .536 | 1.000 | [−0.04, 0.07] |
| CDA | 0.05 | 0.04 | 1.51 | .130 | .520 | [−0.02, 0.12] |

***Note.*** $n = 29$. Models jointly adjusted for target location and report type, separating within- and between-participant distance. $b$ is the within-participant amplitude change in µV per numerical-distance unit. N2pc, Pd, P3b, and LPC included participant random intercepts and distance slopes; CDA retained a random intercept after singular random-slope fits. Final models converged without singular fits. Raw $p$ values are from two-sided Satterthwaite $t$ tests; $p_{Holm}$ adjusts five components. CIs are unadjusted; significant $p$ values are bold. This analysis is exploratory. Adjustment for report type describes a conditional association and does not establish causality.

**CDA analyses with alternative electrode definitions**

The supplementary P7/P8 and combined P7/P8–PO7/PO8 analyses did not reliably distinguish correct T from TT during the 450–700 ms CDA window **(Figure 2—figure supplement 10; Table S21)**. Using the same 29 participants and artifact-free condition averages, we measured contralateral-minus-ipsilateral activity at P7/P8 and the mean of the P7/P8 and PO7/PO8 difference waves. Neither mean-amplitude comparison was significant (P7/P8: $t(28) = -0.06$, $p = .956$; combined: $t(28) = -0.16$, $p = .870$; both $p_{\text{Holm}} = 1.000$). Waveform tests found no significant cluster overlapping the CDA window. The combined definition did show an earlier difference at 308–372 ms (joint $p_{\text{cluster}} = .014$), which preceded the CDA window and should not be interpreted as a CDA effect.

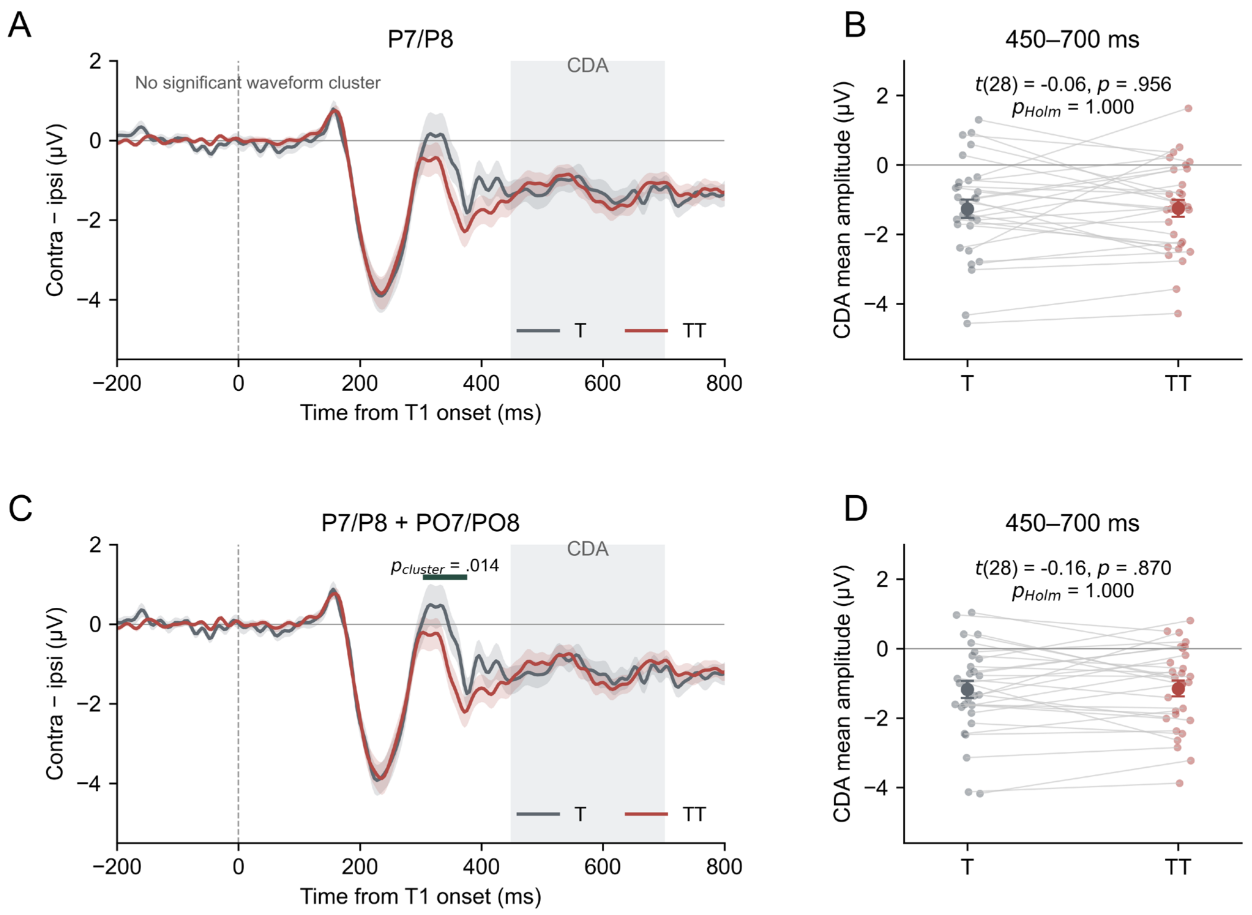


**Figure 2—figure supplement 10. CDA waveforms and amplitudes with alternative electrode definitions.**

Correct T (gray) and TT (red), $n = 29$. A–B: P7/P8; C–D: the mean of the P7/P8 and PO7/PO8 contralateral-minus-ipsilateral waves. A and C show grand averages ± *SEM*; shading marks the 450–700 ms CDA window. B and D show paired participant amplitudes, with larger points and error bars indicating $M \pm SEM$. Positive voltages are plotted upward in every panel. The green bar in C marks the early 308–372 ms cluster. Two-sided paired waveform tests used 20,000 randomizations over −200–800 ms, with a maximum-cluster correction jointly across time, both tails, and the two electrode definitions. Mean-amplitude $p_{\text{Holm}}$ values adjust the two electrode definitions. The significant early cluster lies outside the CDA window.

**Table S21. Alternative-electrode CDA analyses for correct T and TT.**

**A. Mean amplitude at 450–700 ms.**

| Electrode definition | T *M* (*SD*) | TT *M* (*SD*) | T – TT [95% CI] | *t* | *p* | $p_{\text{Holm}}$ |
|---|---|---|---|---|---|---|
| P7/P8 | −1.26 (1.43) | −1.25 (1.32) | −0.01 [−0.51, 0.49] | −0.06 | .956 | 1.000 |
| P7/P8 + PO7/PO8 | −1.18 (1.31) | −1.15 (1.21) | −0.03 [−0.41, 0.36] | −0.16 | .870 | 1.000 |

***Note.*** Amplitudes and differences are in µV. CIs are 95% familywise Bonferroni intervals across two electrode definitions; $p_{\text{Holm}}$ adjusts the same two paired tests. The mean-amplitude procedure matched the primary analysis, including prestimulus baseline correction and ERPLAB spline interpolation. Signed effect sizes were $d_z$ = −0.010 (P7/P8) and −0.031 (combined).

**B. Waveform clusters with joint $p_{\text{cluster}}$ < .10.**

| Electrode definition | Time (ms) | Joint $p_{\text{cluster}}$ | Overlaps CDA |
|---|---|---|---|
| P7/P8 | 332–372 | .061 | No |
| P7/P8 | 388–428 | .096 | No |
| P7/P8 + PO7/PO8 | 308–372 | .014 | No |
| P7/P8 + PO7/PO8 | 388–428 | .074 | No |

***Note.*** Native 250-Hz waveforms were tested without additional filtering or smoothing. Joint $p_{\text{cluster}}$ values control both electrode definitions and the full −200–800 ms search interval. No cluster with $p_{\text{cluster}}$ < .10 overlapped 450–700 ms.
